# An evolutionary timescale for Bacteria calibrated using the Great Oxidation Event

**DOI:** 10.1101/2023.08.08.552427

**Authors:** Adrián A. Davín, Ben J. Woodcroft, Rochelle M. Soo, Benoit Morel, Ranjani Murali, Dominik Schrempf, James Clark, Bastien Boussau, Edmund R. R. Moody, Lénárd L. Szánthó, Etienne Richy, Davide Pisani, James Hemp, Woodward Fischer, Philip C.J. Donoghue, Anja Spang, Philip Hugenholtz, Tom A. Williams, Gergely J. Szöllősi

## Abstract

Most of life’s diversity and history is microbial but it has left a meagre fossil record, greatly hindering understanding of evolution in deep time. However, the co-evolution of life and the Earth system has left signatures of bacterial metabolism in the geochemical record, most conspicuously the Great Oxidation Event (GOE) ∼2.33 billion years ago (Ga, (Poulton et al. 2021)), in which oxygenic photosynthesis and tectonism (Eguchi, Seales, and Dasgupta 2019) transformed Earth’s biosphere from dominantly anaerobic to aerobic. Here, we combine machine learning and phylogenetic reconciliation to infer ancestral transitions to aerobic lifestyles during bacterial evolution. Linking these transitions to the GOE provides new constraints to infer the timetree of Bacteria. We find that extant bacterial phyla are truly ancient, having radiated in the Archaean and the Proterozoic: the oldest include Bacillota (Firmicutes), which radiated 3.1-3.7 Ga, Cyanobacteria (3.3-3.5 Ga) and Patescibacteria (3-3.5 Ga). We show that most bacterial phyla were ancestrally anaerobic and that most transitions to an aerobic lifestyle post-dated the GOE. Our analyses trace oxygen production and consumption back to Cyanobacteria. From that starting point, horizontal transfer seeded aerobic lifestyles across bacterial diversity over hundreds of millions of years. Our analyses demonstrate that the diversification of aerobes proceeded in two waves corresponding to the GOE and to a second sustained rise in atmospheric O_2_ at the dawn of the Palezoic (Krause et al. 2022).

---

Establishing evolutionary timescales requires relaxed molecular clocks (Zuckerkandl and Pauling 1965; Drummond et al. 2006) in which rates of molecular evolution (the “ticking” of the clock) can be calibrated using fossils of known ages that are assigned to the internal nodes of an evolutionary tree (Ho and Duchêne 2014; dos Reis, Donoghue, and Yang 2016). The greatest problem for estimating the timetree of Bacteria is a dearth of fossil evidence for constraining the maximum age of clades. Maximum clade-age calibrations can be established for more readily fossilisable organisms, like animals and plants, based on evidence of absence qualified by preservational, environmental and biogeographic controls (J. L. Morris et al. 2018; dos Reis et al. 2015). However, this approach is not possible for most microbes since they leave few interpretable fossils, if any. As a result, the only credible maximum constraint for the great majority of lineages is the Moon-forming impact 4.52 billion years ago ((Hartmann and Davis 1975; Jacobson et al. 2014), which would have been effective in planetary-scale sterilization (Betts et al. 2018).

In the absence of fossils, traces of biological activity preserved in the geochemical record provide an alternative source of information on the age of key lineages and metabolisms (Knoll, Bergmann, and Strauss 2016). Arguably the most striking event recorded in the geochemical record is the GOE, when oxygen began to accumulate in the Earth’s atmosphere ∼2.33 Ga (Lyons, Reinhard, and Planavsky 2014; Poulton et al. 2021). The GOE was the result of the emergence of a new prokaryotic metabolism, oxygenic photosynthesis, which geochemical evidence shows had evolved by ∼3.22 Ga and has been attributed to Cyanobacteria (Johnson et al. 2013; Dismukes et al. 2001; Soo, Hemp, and Hugenholtz 2019). The consensus view is that, despite the evolution of oxygenic photosynthesis some 900 Myr prior, most life was anaerobic prior to the GOE (Fischer, Hemp, and Valentine 2016). As oxygen levels rose, anaerobic life either retreated to anoxic niches or adapted to tolerate or make use of oxygen, giving rise to the wide variety of anaerobic, microaerophilic, and aerobic lifestyles distributed across the tree of life today (R. L. Morris and Schmidt 2013). If this scenario is correct, most transitions to aerobic lifestyles would have happened during or after the GOE, potentially providing a maximum age for most aerobic lineages that can help to resolve the bacterial timetree. Here we develop and validate a probabilistic approach that constrains most aerobic lineages to after the GOE, but allows for aerobic lineages to predate it given sufficient evidence from fossils and genomic data.

## Inferring the evolution of aerobic lifestyles on the bacterial tree

To evaluate the potential of the GOE to provide a maximum age constraint on aerobic lineages, and so to calibrate the bacterial time tree, we first determined transitions to aerobic lifestyles in the species tree of Bacteria. To achieve this, we inferred a species tree of 1007 bacterial genomes representing most orders in the Genome Taxonomy Database ((Parks et al. 2018); Figure 1, Supplementary Material). To study the evolution of oxygen use in Bacteria, we retrieved experimental data on the ability of extant isolates to grow in the presence of oxygen from the BacDive database (aerobic or anaerobic; in what follows, we use “aerobe” to refer to an organism that can grow in the presence of oxygen, see discussion in Supplementary Material) for 3184 species covering 249 GTDB orders (Reimer et al. 2022). As Cyanobacteria are not included in BacDive, we established oxygen usage from the literature for this important group (see Supplementary Material, (Fujita, Tsujimoto, and Aoki 2015)). We evaluated seven different machine learning algorithms to infer the relationship between gene content and oxygen use for extant Bacteria (see Supplementary Material). Each of the classifiers predicted oxygen use for the experimentally determined cases with >95% accuracy (Figure S3, Table S2). We selected the gradient boosting method XGBoost (Ayyadevara 2018) for subsequent analyses because it had the best combination of accuracy and robustness to noise in the gene contents used as input.

**Figure 1.**
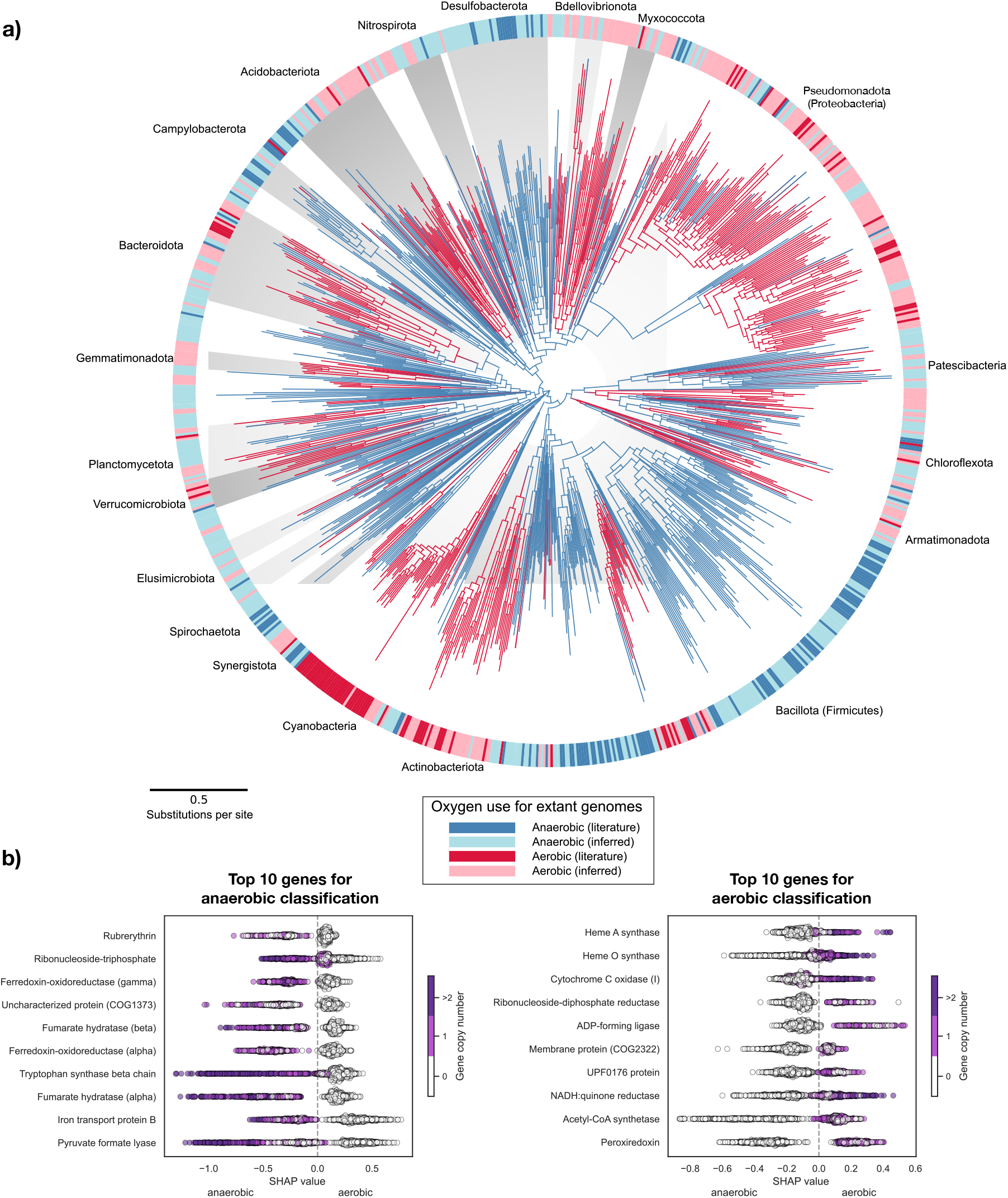
The evolution of aerobic lifestyles in Bacteria. (a) A species tree of Bacteria (inferred using 65 marker genes from 1007 genomes using a custom site-heterogeneous substitution model), with branches coloured according to predicted ability to grow in the presence of oxygen (red, aerobic; blue, anaerobic). The external disc shows observed (dark colours) or predicted (light colours) oxygen use phenotypes for extant genomes. Major bacterial phyla (with more than 15 genomes in our tree) are named and delimited by shading. Branch lengths are proportional to the expected number of substitutions per site, as indicated by the scale bar. (b) Hallmark genes of aerobic and anaerobic Bacteria. The plots show genes with the strongest contribution (SHAP values) to classification as aerobic or anaerobic with the XGBoost classifier (see main text and SM). Each dot represents one node in the tree. On the left, genes where a higher copy number predicts an anaerobic lifestyle; on the right, genes where a higher copy number predicts an aerobic lifestyle. The genes most diagnostic of aerobic lifestyles include core components of aerobic metabolism such as the heme-copper oxygen reductase (HCO) and enzymes for the biosynthesis of heme A, the co-factor of the most widely distributed O2 reductase (A-family) from the HCO superfamily. The genes most closely correlated with the prediction of an anaerobe are genes such as pyruvate formate lyase and fumarate hydratase, which are known to be important for anaerobic central metabolic pathways.

To identify the branches on the tree where transitions between anaerobic and aerobic lifestyles occurred, we reconstructed ancestral gene contents for all internal nodes of the species phylogeny using the probabilistic gene tree-species tree reconciliation method ALE (Szöllõsi et al. 2013; Szöllősi et al. 2015) and applied the XGBoost classifier to these nodes to infer the ability of ancestral organisms to grow in the presence of oxygen. Since ancestral gene contents can only be inferred with significant uncertainty, we explored the impact of false positives and negatives (spuriously present or absent gene families) on our ancestral state inferences. Even with a joint 50% false negative and false positive rate, XGBoost retained >90% accuracy (Figure S3). This robustness implies that many gene families distinguish aerobes from anaerobes, perhaps because the ability to grow in the presence of oxygen leaves a broad imprint on the genome. This is evident from the list of genes with the largest effect on the XGBoost predictions, including genes directly involved in aerobic respiration (Supplementary Material). To cross-check the results obtained by the gene content-based classifier, we reasoned that oxygen use phenotypes may also have left an imprint in the amino acid sequences of universal genes. According to this hypothesis, different amino acids may be favoured in aerobic vs anaerobic species at particular sites. To test this hypothesis, we used the concatenate of universal marker genes employed to reconstruct the tree of Bacteria and learned the relationship between amino acid state at each site and oxygen use phenotype for extant species, using elastic net logistic regression (Supplementary Material). We then reconstructed ancestral sequences for all nodes on the species tree and applied the amino acid classifier to ancestral sequences to predict ancestral oxygen use phenotype. These analyses were also accurate, with a leave-one-out prediction accuracy of 83.7%, and in good agreement with the gene content-based classifier for the ancestral nodes in the tree (overall 90% with XGBoost). This shows that oxygen use has left two distinct but consistent imprints in genomes, in gene content on one hand and, on the other, in site-specific amino acid usage in universal genes. By identifying branches in our reference tree with different XGBoost-predicted aerobic states for parental and descendant nodes, we established a map of aerobic transitions on the bacterial phylogeny, to which the GOE constraint could be applied in dating analysis (Figure 1).

## A bacterial time tree calibrated with the GOE

To infer the timescale of bacterial evolution, we expanded our species tree to include genes from mitochondria and chloroplasts from diverse eukaryotes. Mitochondria and chloroplasts are descended from Bacteria but, since they are eukaryotic organelles, their ages can be constrained using the eukaryotic fossil record, greatly improving the number of fossil calibrations available for dating the tree (of 18 total geological calibrations in our analysis, 12 are from the eukaryotic record including 8 maximum age calibrations) (Shih et al. 2013; Mahendrarajah et al. 2023). Since plants and algae have both mitochondria and chloroplasts, the same species divergences occur twice in the species tree, for example, the divergence between red and green algae appears in both the mitochondrial and plastid clades. These equivalent nodes can be “braced” (fixed to the same unknown age (Shih et al. 2013)) in the dating analysis, providing valuable additional calibration information. (Figure 2).

**Figure 2:**
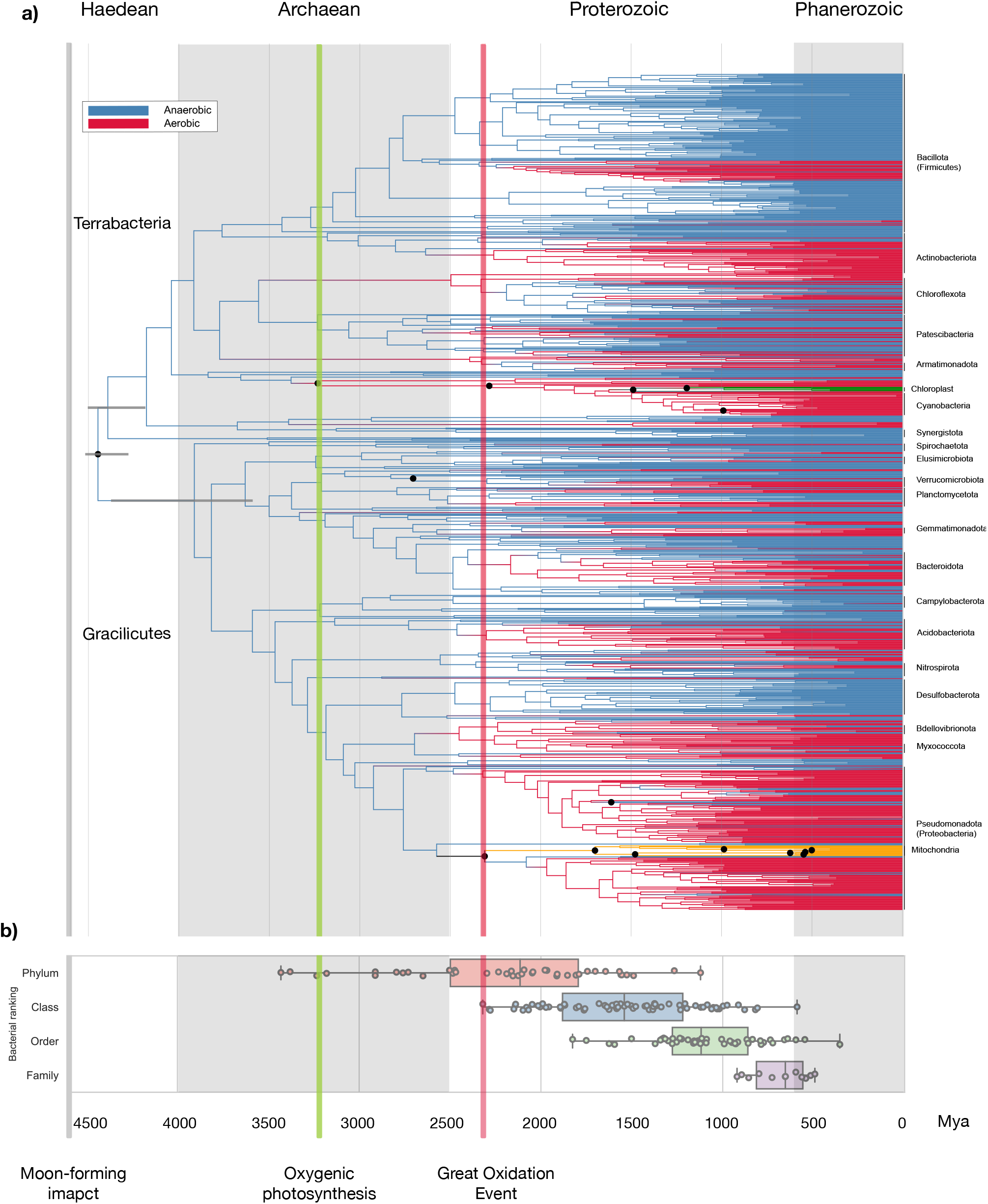
A dated phylogeny of Bacteria. **a)** dated phylogenetic tree of Bacteria. Branch colours represent anaerobic (blue) and aerobic lineages (red). Mitochondrial branches are colored in orange while chloroplast branches are coloured in green. Branch lengths are proportional to geological time (Ga). 95% highest posterior densities are shown for the root, and the two major bacterial divisions Gracilicutes and Terrabacteria. The grey line shows the Moon-forming impact at ∼4.52 Ga; the green line at ∼3.23 Ga reflects the presence of fossil and isotopic evidence for oxygenic photosynthesis; the red line at ∼2.33 Ga reflects the end of the GOE (see calibrations in Supplementary Material). The bottom panel shows the ages of taxonomic groups. The names of phyla represented by 8 or more genomes are shown. We show the age of the last common ancestors of major taxa (phyla, class, order and family) represented in our 1007 genomes dataset by at least 3 genomes. The median ages are: phyla: 2115 Ma; class: 1524 Ma; order: 11120 Ma; family: 655 Ma. Black dots denote nodes that were directly calibrated in molecular clock analyses.

We then performed Bayesian relaxed molecular clock analyses under three different conditions: with only the Moon-forming impact as a maximum age on bacterial evolution; with the Moon-forming impact as a maximum and 17 additional fossil and geochemical calibrations (see Supplementary Material), (“Fossils”); and combining those sources of information with the GOE constraint on the age of aerobic nodes (“Fossils+GOE”). Note that, while transitions to oxygen use were likely more common after the GOE, it is clear that at least some aerobic lineages evolved beforehand. Indeed, there is good evidence for a level of free oxygen indicative of cyanobacterial activity from at least 3.225 Ma based on stable Fe and U-Th-Pb isotopes in the Manzimnyama Banded Ironstone Formation (Fig Tree Group, Barberton, South Africa; (Satkoski et al. 2015)). To account for this, we implemented the GOE constraint as a soft maximum in the Bayesian analysis; this reduces the probability that aerobic nodes are older than the GOE, but they can nonetheless pre-date the GOE if there is sufficient evidence from fossils or sequence divergence to support it. The impact of the GOE constraint is generally to shift the origin of aerobic bacterial lineages towards the present, with the mean age of bacterial phyla moving 380 million years (My) towards the present when this constraint is applied. However, we recover evidence for an earlier emerging aerobic linages, as well as oxygen-producing Cyanobacteria, in both our the “fossils” and “fossils+GOE” analyses.

## The GOE constraint improves the accuracy of the bacterial time tree

We used the GOE constraint based on the prior hypothesis that aerobic metabolisms would have been more common after the GOE, than before. However, given that some aerobic bacterial lineages pre-date the GOE in both the “fossils” and “fossils+GOE” analyses, it seems reasonable to ask whether applying the GOE soft maximum improves inference of the bacterial timetree. We reasoned that the histories of gene families might, in principle, carry phylogenetic signal to distinguish the dated trees because gene transfers can only occur between species that exist at the same time (Daubin and Szöllősi 2016; Davín et al. 2018). We harnessed this information using a novel time-constrained gene tree-species tree reconciliation model that only allows transfer between branches that overlap in time to calculate the likelihood of each gene family under each candidate dated species tree (see Supplementary Material). As the different dated species trees vary in their branch lengths, the possible reconciliation scenarios differ among the trees. The summed log-likelihood of all reconciled gene families therefore provides a metric for choosing among different dated trees, much as site likelihoods can be used to choose among tree topologies in traditional phylogenetics (Shimodaira and Hasegawa 2001; Szöllosi et al. 2012; Harris, Sheridan, et al. 2022). To validate the GOE soft maximum calibration on aerobic nodes we compared support for the three inferred timescales using 4731 gene families from the 1007 genomes. As expected, the time tree calibrated only with the Moon-forming impact had the lowest likelihood (was least compatible with gene family histories) under the time-constrained gene tree species tree reconciliation model. Calibrating the tree with fossils substantially improved the likelihood, while combining fossils with the GOE constraint further increased the likelihood and produced the best results (Table 1). The improvement in time-constrained likelihood is greatest when using XGBoost to predict node phenotypes, though a substantial improvement in likelihood was observed for all predictors of aerobicity tested (Table 1, Figure S13).

**Table 1.** Comparison of likelihoods under the time-constrained DTL model. 4731 families were reconciled with different consensus chronograms. We show both the total LL (resulting from adding all the LLs from every reconciliation) and the difference with the least likely chronogram (Moon-forming impact as the only calibration implemented as a maximum age calibration on the root).

| Condition | Summed LL<br>(4731 gene families) | LL improvement over MFI<br>(4731 gene families) |
| --- | --- | --- |
| Moon-forming impact (MFI) | -4807906 | 0 |
| MFI + Fossils | -4807655 | 251.7682 |
| MFI + Fossils + GOE<br>(Concatenate) | -4807626 | 280.6226 |
| MFI + Fossils + GOE<br>(ExtraTrees) | -4807410 | 496.0085 |
| MFI + Fossils + GOE (XGBoost) | -4807223 | 683.0737 |

## A revised chronology of bacterial evolution

Our analyses support the view that the deepest divergences within the bacterial domain occurred early in Earth history (Figure 3). We estimate that the last bacterial common ancestor (LBCA) lived 4.5-4.3 Ga, while the major clades of Gracilicutes and Terrabacteria radiated 4.4-3.6 Ga and 4.5-4.2 Ga (respectively), substantially earlier than previous estimates (Battistuzzi and Hedges 2009). The extant phyla with the oldest crown groups are Bacillota (formerly Firmicutes; 3.7-3.1 Ga), Patescibacteria (3.5-3.0 Ga), Actinobacteriota (3.5-2.8 Ga) and Cyanobacteria (3.5-3.3 Ga; as defined by GTDB taxonomy, including the three classes Cyanobacteriia, Vampirovibrionia and Sericytochromatia; (Oren, Mareš, and Rippka 2022; Garcia-Pichel et al. 2020). However, the crown of the photosynthetic lineage (Class Cyanobacteria) is estimated to be substantially younger (2.5-2.0 Ga, Figure S18), in agreement with other studies that have estimated that the three classes diverged between 2.8-2.0 Ga (Shih et al. 2017; Magnabosco et al. 2018; Oliver et al. 2021; Boden et al. 2021); published age ranges are reviewed in (Sánchez-Baracaldo et al. 2022). Pseudomonadota (formerly Proteobacteria; crown age 3.0-2.6 Ga) are somewhat younger than Cyanobacteria (3.0-2.6 Ga) and other major terrabacterial phyla despite their extant phylogenetic and metabolic diversity.

**Figure 3.**
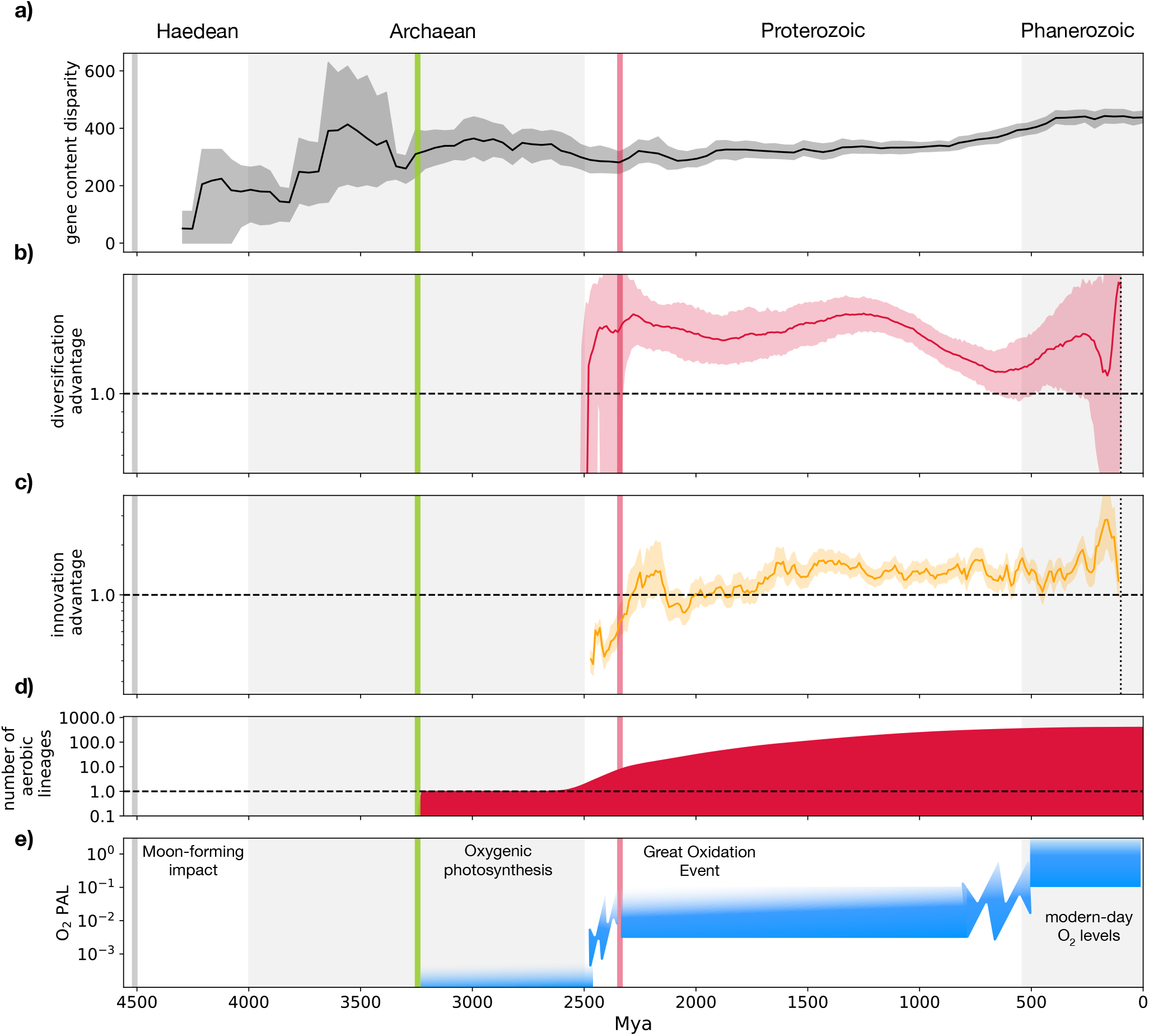
Bacterial evolution through geological time. (a) The early evolution of bacterial genetic diversity in the Archaean (David and Alm 2010) is reflected by rapid increase in gene content disparity (see Supplementary Material). (b) A persistent higher diversification rate of aerobes since the GOE. The line charts the relative diversification rate (the ratio of the inverse of the waiting time to the next speciation event) for aerobic and anaerobic lineages, revealing a persistent higher diversification rate of aerobes for most of the Earth’s history since the GOE. Rates equilibrated prior to the NOE, before a return to higher diversification rates for aerobes since the GOE. The dotted line denotes a ratio of 1, where the diversification rate of aerobic and anaerobic lineages is equal. Terminal branches were omitted and ratios since 100 Mya were not calculated do to a low number of nodes. (c) Spread of aerobic metabolism by HGT. Transitions from anaerobic to aerobic lifestyles have outnumbered transitions in the reverse direction/back transitions/returns to anaerobicity for most of Earth history since the GOE. The plots show the ratio of aerobic to anaerobic transition rates on the dated species tree during each time period. The dotted line denotes a ratio of 1, where transitions occur in both directions at the same rate. The small number of nodes since ∼200 Ma greatly increases uncertainty for both analyses, as reflected in the wide confidence intervals of posterior node age estimates. Ratios since 100 Mya were not calculated due to a low number of nodes. (d) shows the number of aerobic lineages according to Figure 2, e) qualitative sketch of atmospheric oxygen levels through time based on (Lyons, Reinhard, and Planavsky 2014; Poulton et al. 2021; Krause et al. 2022). *The grey line shows the Moon-forming impact at ∼4*.*52 Ga; the green line at ∼3*.*23 Ga reflects the presence of fossil and isotopic evidence for oxygenic photosynthesis; the red line at ∼2*.*33 Ga reflects the end of the GOE (see calibrations in Supplementary Material)*.

We find that the relative evolutionary divergence (RED) metric used by GTDB to partition prokaryotic diversity into taxonomic groups of similar age correlates well with absolute geological time, with a Spearman correlation coefficient (ρ) of 0.93 between RED values and the age of nodes (Figure S6). A previous study focusing on fungal taxonomy found a similar consistency between RED and relative divergence times using a relaxed molecular clock approach (Li et al. 2021). Our analysis makes plain the enormous difference in evolutionary timescales for prokaryotic and eukaryotic diversity. For example, we estimate that the last common ancestor of extant mitochondria (and therefore of extant eukaryotes) lived 1880-1502 Ma. The eukaryotic domain of life is therefore comparable in age to a bacterial class, while the crown ages of groups such as metazoans (animals) 833-650 Ma (dos Reis et al. 2015; Dohrmann and Wörheide 2017)) and embryophytes (land plants) 515-494 Ma (Harris, Clark, et al. 2022)) correspond roughly to bacterial families whose ages range from 899-457 Ma in our analysis (Figure 2). Indeed, many modern bacterial families were already established in the Precambrian (before the Phanerozoic and Proterozoic Eras boundary in Fig. 2.). These results are important since they inform attempts to standardise prokaryote systematics as it is revised by molecular phylogenetics, to achieve temporal consistency among clades of equivalent taxonomic rank (Parks et al. 2018). Taken together, our observations highlight the sparsity of bacterial ranks in geological time relative to eukaryotes and indicate that temporal harmonization of prokaryotic and eukaryotic taxonomy would require introduction of additional ranks in prokaryotic taxonomy and commensurate pruning of eukaryotic ranks (Avise and Johns 1999). Furthermore, Figure 2 reveals extensive phylogenetic structure and evolutionary time above the level of bacterial phylum, which supports the recent proposal for additional ranks between domain and phylum (Göker and Oren 2023).

## Mechanisms underlying transitions between anaerobic and aerobic lifestyles

Returning to the history of adaptation to aerobic conditions across the bacterial tree, LBCA and the common ancestors of Gracilicutes and Terrabacteria are inferred to be anaerobic. Indeed, most (38 out of 49) bacterial phyla are inferred to be ancestrally anaerobic, consistent with their emergence prior to the GOE. As expected, the majority (68 of 84) of transitions to aerobic metabolism happened after the GOE; however, the observation that 16 transitions occurred pre-GOE supports the view that some niches existed for aerobes before the oxygenation of the atmosphere. Interestingly, the earliest transition to aerobic metabolism mapped to the bacterial species tree is the common ancestor of the cyanobacterial classes Cyanobacteriia and Vampirovibrionia, which implies that aerobic lifestyles and oxygenic photosynthesis evolved very close together in time (Fig S17-19).

To investigate the biochemical potential that is strongly associated with extant aerobes, we identified the genes that are correlated with a prediction of aerobicity (Figure 1, SHAP values). The best predictor is related to the A-family heme copper oxygen (HCO) reductase, the most widely distributed terminal O2 reductase in bacteria (Pereira, Santana, and Teixeira 2001; Murali, Hemp, and Gennis 2022), its constituent catalytic subunit I, as well as enzymes (heme O and heme A synthases) involved in the synthesis of its cofactor, heme A. In aerobes, another enzyme involved in O2 respiration, ndh-II (type-II NADH dehydrogenase) is also found to be important and is preferred over type-I NADH dehydrogenase during aerobic growth (Meng, Green, and Guest 1997). Proteins that protect against oxidative damage such as peroxiredoxin and ferritin are also commonly found in aerobes. To identify the metabolic drivers of transitions to an aerobic lifestyle that may have changed over time, we compared genes and gene functions gained and lost in the 84 independent aerobic transitions identified. To do so, we performed enrichment analysis of Gene Ontology (GO) terms on transition branches, considering transitions that occurred before and after the GOE separately. While we did not find a significant enrichment of GO terms among genes that were lost, we did find a number of GO terms enriched among genes gained by horizontal transfer on transition branches. Consistent with their presence in modern aerobes, the A-family O2 reductase, and heme O and A synthases were found to be often enriched in transition branches. Curiously, more recent transitions to aerobes from anaerobes are associated with different markers for aerobic respiration such as the less widely distributed HCO (C-family O2 reductase), members of the other O2 reductase superfamily, the cytochrome bd oxygen reductases and the cytochrome bc1 complex. Other metabolic markers for aerobic transition are the presence of enzymes within the oxidative part of the TCA cycle, beginning with the oxidation of 2-oxoglutarate all the way to CO2 and with some monooxygenase reactions such as those involved in aerobic degradation of phenyl acetate. Biosynthesis pathways for cofactors such as siroheme and ubiquinone are also associated with transition to aerobicity; ubiquinone has been previously identified as a marker for high-potential metabolism (B Schoepp-Cothenot et al. PNAS 2009). Transporters and chaperones for the use of transition metals such as copper and molybdenum are enriched in the aerobic transitions, consistent with the evidence that both the bioavailability and use of these transition metals increased during the oxidation of the Earth in the early Proterozoic (David and Alm 2010; Zerkle, House, and Brantley 2005). In fact, copper is essential for aerobic respiratory pathways that involve the HCO enzymes ((Pereira, Santana, and Teixeira 2001; Murali, Hemp, and Gennis 2022)). Enzymes involved in mediating redox stress such as catalase are also important for anaerobes to become aerobes. Another unexpected association with the ancient transition to aerobic lifestyles is with many large subunits in the ribosome and aminoacyl tRNA synthetases. The sensitivity of protein translation machinery to oxidative stress has been noted (Ling and Söil Biochemistry, 2010, Chan et al. 2012 Nature Communications) and our results suggest that protein translation must have adapted in response to the rise of oxygen in the atmosphere.

## Aerobes diversified faster than anaerobes

We estimated rates of gene content evolution by combining the dated tree with the gene tree-species tree reconciliations. Normalised by genome size, numbers of gene transfers and losses per million years were broadly similar in aerobes and anaerobes, with about one transfer and one loss per million years, while inferred rates of gene duplication were about 100-fold lower (see Figure S16-17).

To evaluate how bacterial gene repertoires have evolved over time, we applied non-metric multidimensional scaling to quantify the disparity among the gene content of modern and ancestral Bacteria, using the reconstructed ancestral gene repertoires as ancestral states. Plotting disparity through time (Figure 3a) provided clear support for an “Archaean expansion” (David and Alm 2010) during which most variation in bacterial gene repertoires was already established prior >3.5Ga during the radiation of the Terrabacteria and Gracilicutes clades.

Next, we investigated the impact of the transition to aerobic metabolisms on bacterial evolution. To compare the relative diversification rate of aerobes and anaerobes, we used a metric based on the time intervals between speciation events (Figure 3b). This analysis suggests two main periods of adaptation to aerobic life: a rapidly established diversification advantage of aerobes that began at the GOE (cf. Figure 3e) and was sustained for the following ∼1.5Gyr, that waned and then rebounded during the last 500 million years. A higher diversification rate for aerobes is also evident in a complementary metric that measures the directionality of transitions between aerobic and anaerobic metabolism (Figure 3c). We interpret this pattern to reflect a combination of the creation of aerobic niches at the GOE and additional higher-oxygen aerobic niches as levels of atmospheric free oxygen underwent a second sustained rise in the Paleozoic (Krause et al. 2022) and the selective extinction of early anaerobic lineages as a consequence of oxygenation, reduction and fragmentation of their niche space and competition for resources with aerobes.

In addition to the two periods of aerobic advantage, both diversification metrics also record a late reversion of the earlier trends, in which anaerobes are at an advantage ∼200Ma (Figure 4). These inferences are uncertain because confidence intervals are wider in this time period owing to the low number of speciation nodes. Nonetheless, it seems possible that this signal reflects adaptation to new anoxic niches, such as the gastrointestinal tracts of early animals or anaerobic niches within decaying plant biomass. Consistent with this hypothesis, many of the detected reversions to anaerobic metabolism are seen in lineages that are now obligately host-associated.

## Conclusions

Our analyses demonstrate a new way to link bacterial gene content to phenotype, and to trace those relationships back in time. By combining phylogenetic reconciliation with machine learning classification on extant and ancestral gene repertoires and sequences, we predicted the oxygen use phenotypes of ancestral nodes in the bacterial tree. This allowed us to calibrate bacterial evolution to the record of biospheric oxygenation, greatly augmenting the limited fossil record of early life and bringing a new level of resolution to the study of evolution in the Precambrian. Our timetree highlights the disparity between bacterial and eukaryotic classification: most extant bacterial phyla radiated in the Archaean and early Proterozoic, pre-dating the eukaryotic domain of life. The earliest aerobes that left extant descendants lived in the Archaean (3.22-3.25 Ga), pre-dating the GOE by about 900 Mys and gave rise to the Cyanobacteriia and Vampirovibrionia. Spread of aerobic metabolism occurred later, proceeding in two main phases coinciding with the GOE and NOE. This was enabled by the widespread transfer of respiratory complexes, and lineages that became aerobic subsequently experienced a higher diversification rate than contemporary anaerobes. While aerobic metabolism was an obvious choice, any trait might be explored with these new dating approaches. We anticipate that high-resolution time trees of microbial life will greatly enrich our understanding of the connection between biological and geological evolution.

## Data availability

All data used is described in detail in the Supplementary Material and available in the figshare repository <u>10.6084/m9.figshare.23899299</u> associated with the submission.

## Code availability

All code used is described in detail in the Supplementary Material and available either openly on github or other relevant public repository, as indicated or included in the figshare repository <u>10.6084/m9.figshare.23899299</u> associated with the submission in the case of more specialised scripts.

## Acknowledgements

We thank Julia A. Palacios for providing R scripts to compute the D2 distance for ranked trees. This work was supported by the Gordon and Betty Moore Foundation through grant GBMF9741 to TAW, AS, and GJSz. AD, DS, LLSz DS and GJSz received funding from the European Research Council under the European Union’s Horizon 2020 research and innovation program under Grant Agreement (grant agreement No. 714774, GENECLOCKS). AS has received funding from the European Research Council (ERC) under the European Union’s Horizon 2020 research and innovation programme (grant agreement No. 947317, ASymbEL), the Moore–Simons Project on the Origin of the Eukaryotic Cell, Simons Foundation 735929LPI (https://doi.org/10.46714/735929LPI), and a Gordon and Betty Moore Foundation’s Symbiosis in Aquatic Systems Initiative (GBMF9346). Further, this work was supported by a Royal Society University Research Fellowship to TAW. ERRM, DP, PCJD and TAW were supported by the John Templeton Foundation (62220). BJW is supported by an Australian Research Council Future Fellowship (FT210100521). AAD and PH were supported in part by an Australian Research Council Laureate Fellowship (FL150100038).

