## Supplementary Material for "An evolutionary timescale for Bacteria calibrated using the Great Oxidation Event"

### Methods

1. Species tree inference
    - Genome preselection
    - Marker filtering
    - Genome sampling refinement
    - Markers preselection
    - Species tree inference with an archeal outgroup
    - Bacterial species tree inference using the dataset-specific EDM model
    - Adding mitochondrial and plastid genomes to the dataset

2. Gene tree-species tree reconciliations
    - Gene family alignments
    - Alignment preparation
    - Inference of bacteria-specific site heterogeneous substitution models
    - Comparison of model fit
    - Refinement of the alignments based on the EDM trees
    - Reprocessing of the subclusters
    - Inference of subcluster's gene trees
    - Reconciliations with ALE

3. Phenotype prediction
    - Prediction via gene content
    - Aerobicity prediction using a concatenate of marker genes

4. Calibrations
5. Molecular dating analyses
6. Analyses with time-constrained reconciliations
    - Using the time-constrainted DTL ALE model to distinguish dated trees
    - Simulations
    - Robustness analysis
    - Patterns of Gene Content evolution
7. Enrichment analysis
8. Origin of core photosynthesis genes
9. Analysis of bacterial disparity through time
10. Analysis of aerobic vs anaerobic diversification

### 1. Species tree inference

#### Genome selection

We targeted a dataset of approximately one thousand genomes, as a compromise between having a representative sample of bacterial diversity and a small enough dataset so that we could use the best computational methods available. We used the GTDB taxonomy version r95 (Parks et al. 2018, 2020) since it was the most recent version at the time we started our analyses. The genomes involved in our study were identified using a blend of manual curation and automated processes.

We first extracted available information on growth under aerobic and anaerobic conditions for GTDB genomes from BacDive, downloaded  (Reimer et al. 2019). Following this, we ranked the genomes based on three criteria: firstly, whether the genome was the species type of the genus; secondly, the availability of information concerning the oxygen tolerance of the genome; and lastly, the quality of the genome assembly (defined as the CheckM-completeness – 5* CheckM-contamination). This ranking was employed to create a representative subsample of the GTDB database.

Our methodology involved grouping all organisms by a given taxonomic level and only retrieving the top-ranked organism for each group. Through this subsampling strategy, we generated three separate sets of genomes that were ultimately consolidated into a singular dataset. The first set was comprised of species possessing HCOs (Online data supplement HCOr95.zip), and we subsampled this list at the class level, selecting the top-ranked species within each class. This step ensured that our final dataset adequately represented bacterial groups, capturing the diversity of this gene family. The second set included the top-ranked species per class in GTDB version r95, ensuring that our dataset encapsulated a comprehensive sample of known bacterial diversity. Lastly, we sampled the top-ranked species at either the order or family level for the ten most species-rich bacterial phyla, guaranteeing the most speciose phyla (as determined by GTDB) were well-represented in our final dataset.

Upon assembling these three datasets, we manually incorporated two additional species GB_GCA_013390565.1 and GB_GCA_013390945.1, which were taken from the r202 version of GTDB and chosen due to their status as recent discoveries of genomes using Type I reaction centre for phototrophy, offering potential insights into the evolution of aerobic metabolism. The resulting sample included 1118 genomes, exceeding our initial target. Consequently, we decided to refine our selection further by filtering out certain genomes based on the criteria elucidated in the section entitled 'genome sampling refinement'. The final genome selection is presented in `GenomesInfo.tsv` in the Online Data Supplement.

Ultimately, after applying the XGBoost gene-based prediction algorithm (see below), we found that of the final chosen set of 1007 genomes, 445 of them (44%) were predicted to be aerobic. For comparison, from a randomly sampled set of 57,085 species representatives from GTDB 08-RS214, 30,360 (53%) were predicted to be aerobic.

### Marker preselection

We used 71 markers widely distributed in Bacteria. We took all the markers used in Coleman et al. 2021 (Coleman et al. 2021). We limited the selection to markers that were represented in more than 600 different genomes and had less than 1350 gene copies, to balance the ubiquity of the marker (which we want to maximize) and the number of multicopy genes (which we want to minimize). We based the marker selection on the COG clusters, but we first filtered sequences from every gene family if both the PFAM and the TIGR annotation were different from the most common annotation in the gene family. We also removed families with a 30% difference in size (measured as the number of amino acids) from the median, to avoid having sequences too small or too big introducing noise in the alignments. The markers can be found in the Online Data Supplement with the name `MarkersInfo.xlsx`

#### Marker filtering

For every marker, we computed the alignments (using MAFFT (Minh et al. 2020; Katoh and Standley 2013), with the command `mafft --maxiterate 1000 --localpair`), trimmed them with BMGE (Criscuolo and Gribaldo 2010) and inferred the individual trees with IQ-TREE 2 (Minh et al. 2020) using the model LG+G4+F. The sequences, alignments, trimmed alignments and trees can be found in the Online Data Supplement (compressed folder `Markers_prefiltering.tar.gz`). Some of the markers have multiple copies of some genomes. To decide which copy we would keep we use a Python script that uses the protein distance matrices obtained by IQ-TREE to infer, for all the multicopy sequences, those whose position in the phylogeny is closer to the mean position of that gene across all markers. The script can be found in the Only Data Supplement with the name `sequence_picker.py`. We retained for every multicopy gene the one with the highest score given by this script, removing in total 1313 sequences from a total of 73168 (~1.8%).

### Refining genome selection based on marker gene analysis

To further evaluate the quality of the different markers, we marked as problematic those sequences that meet any of these criteria:

- Sequences whose branch is more than 5 times the median branch distance of leaf branches (gene trees were rooted using MAD (Fernando Domingues Kümmel Tria, Landan, and Dagan 2017))
- Sequences that are twice the median distance or more from the root for all sequences.
- Sequences with 60% of more gaps in the alignment
- Sequences that fail a composition chi-square test ($p < 0.05$), determined by IQ-TREE

We noticed that, across all marker genes examined, problematic sequences often came from the same genomes. We used this information to remove some genomes from the dataset. To do so, we counted how many sequences were marked as problematic. The most problematic genomes were all Patescibacteria. We, therefore, decided not to remove the Patescibacteria according to this criterion because it would greatly reduce the taxonomic coverage of this group. Instead, we ignored the Patescibacteria in this step and, for other groups, removed the genomes that had 3 or more copies marked as problematic (which removed a total of 77 genomes). After this filtering step, we also removed the genomes that were the least present across all the 71 markers (removing those that appear in 50 markers or less). These additional filtering steps resulted in a dataset of 1007 genomes.

#### Marker gene set refinement using an archaeal outgroup

To identify inter-domain gene transfers in the 71 marker genes, we retrieved archaeal homologues, where available, for each gene family, and inferred single gene trees for the combined archaea-bacteria dataset. We then manually inspected these gene trees to identify cases in which bacterial sequences appeared to be the result of horizontal acquisition from archaea (because such sequences will not correctly track the evolutionary history of the bacterial lineage encoding them). Based on manual inspection, 6 of the 71 markers (COG0150, COG0255, COG0460, COG0504, COG0525, COG0540) were removed from the dataset because they showed extensive interleaving of archaeal and bacterial clades, such that it was difficult to establish the number and direction of inter-domain transfers. For two additional markers, COG0343 and COG0496, the marker as a whole was kept but several sequences that had clearly been acquired from archaea were deleted. The resulting 65-gene marker dataset along with the concatenate can be found in the Online Data Supplement with the name: `Concatenate65.tar.gz.`

These marker genes were aligned with `mafft` (`l-ins-i`) (Katoh and Standley 2013) and poorly aligning regions were removed using BMGE (Criscuolo and Gribaldo 2010), with the BLOSUM30 matrix. An LG+G4+F tree was inferred from the 65-gene concatenation using IQTREE 2.1.3 (`-m LG+G+F -B 1000`). The analysis can be found in the Online Data Supplement with the name 65gene_LGGF_analysis.tar.gz. MAD rooting of this tree (using the -g option, to optimise root position within branches) recovers a Gracilicutes-Terrabacteria root. The results from this rooting analysis can be found in the Online Data Supplement with the name 65gene_lggf.treefile.rooted.

Next, we inferred a tree from the 65-gene concatenation using LG+C20+G via the PMSF (Posterior Mean Site Frequency) approximation, which greatly reduces RAM use and computational time for fitting site-heterogeneous mixture models to large alignments (Wang et al. 2018). We first inferred a fixed frequency profile for each site in the alignment using the LG+C20+G mixture model:

```
iqtree2 -s $concatenation -m LG+C20 -ft $LGGF_guidetree -n 0
```

Then performed an ML tree search and bootstrapping using the fixed site-specific profiles:

```
iqtree2 -s $concatenation -m LG+C20 -fs $sitefreq -B 10000
```

The results from this analysis can be found in the Online Data Supplement with the name:
`65gene_analysis_LGC20G_PMSF.tar.gz.`

The MAD-rooted tree (Gracilicutes-Terrabacteria; obtained using mad.py with the -g option) can be found in the Online Data Supplement with the name:

`LGC20GPMSF_65gene.treefile.rooted.`

#### Species tree inference with archaeal outgroup

To test the effect of including an archaeal outgroup on the inferred bacterial topology, we constructed alignments (MAFFT L-INS-I, v.7.4 (Katoh and Standley 2013) ) with the archeal homologues (using the COG identification for these markers from (Moody et al. 2022)) outlined above, for the 71 bacterial marker genes, resulting in 52 alignments with representation from Archaea and Bacteria. We trimmed these 52 alignments using BMGE 1.12 (with default settings in addition to a BLOSUM30 matrix) (Criscuolo and Gribaldo 2010) and inferred initial trees using IQ-TREE 2.0.3 (Minh et al. 2020) with the LG+G+F model, with 10,000 ultrafast bootstraps. Our initial trees were inspected manually, and 11 of the resulting trees were removed due to multiple independent transfer events, paralogous gene families, or possible long-branch-attraction. After removing these genes and pruning sequences which violated domain monophyly, we re-alignedand subsequently trimmed the remaining sequences before another round of manual inspection. We then concatenated these updated 41 alignments, and inferred a tree using IQ-TREE 2 (2.0.3) with LG+G+F with 10000 bootstrap replicates. The concatenate and the individual markers can be found in the Online Data Supplement with the name: `Concatenate41.zip.`

#### Bacterial species tree inference using the dataset-specific EDM model

As part of our subsequent analyses of bacterial gene families, we inferred new, bacteria-specific site-heterogeneous models based on 3064 single-gene alignments using the EDCluster algorithm (https://github.com/dschrempf/EDCluster, (Schrempf, Lartillot, and Szöllösi 2020)). Analysis of model fit suggested that LG+EDM models were the best-fitting models for the majority of alignments, with LG+EDM0128+G the most popular model overall. The EDM models have the added benefit of mixture weights estimated from the 10% sample of alignments, so that weights do not need to be inferred during single gene analysis. We therefore used these models to infer an unrooted species tree from the 65-gene concatenate using the PMSF approach in IQ-TREE 2.1.1. To do so, we inferred a guide tree using LG+G, then used the guide tree to estimate site-specific frequencies using the LG+EDM0064+G model. Model fitting with more than 64 components was impractical on the concatenate, with 1.16TB RAM required for fitting the 128 component mixture model. The LG+G guide tree, the LG+EDM0064+G ML treefile (10000 UFBoot bootstraps) and the MAD-rooting on the LG+EDM0064+G (PMSF) can be found in the Online Data Supplement with the name PMSFtree.tar.gz. As with the LG+C20+G analysis reported above, the optimal MAD root was recovered between Gracilicutes and Terrabacteria; Spirochaetes grouped with Gracilicutes, while Wall-, Muir- and Fusobacteriota grouped with Terrabacteria.

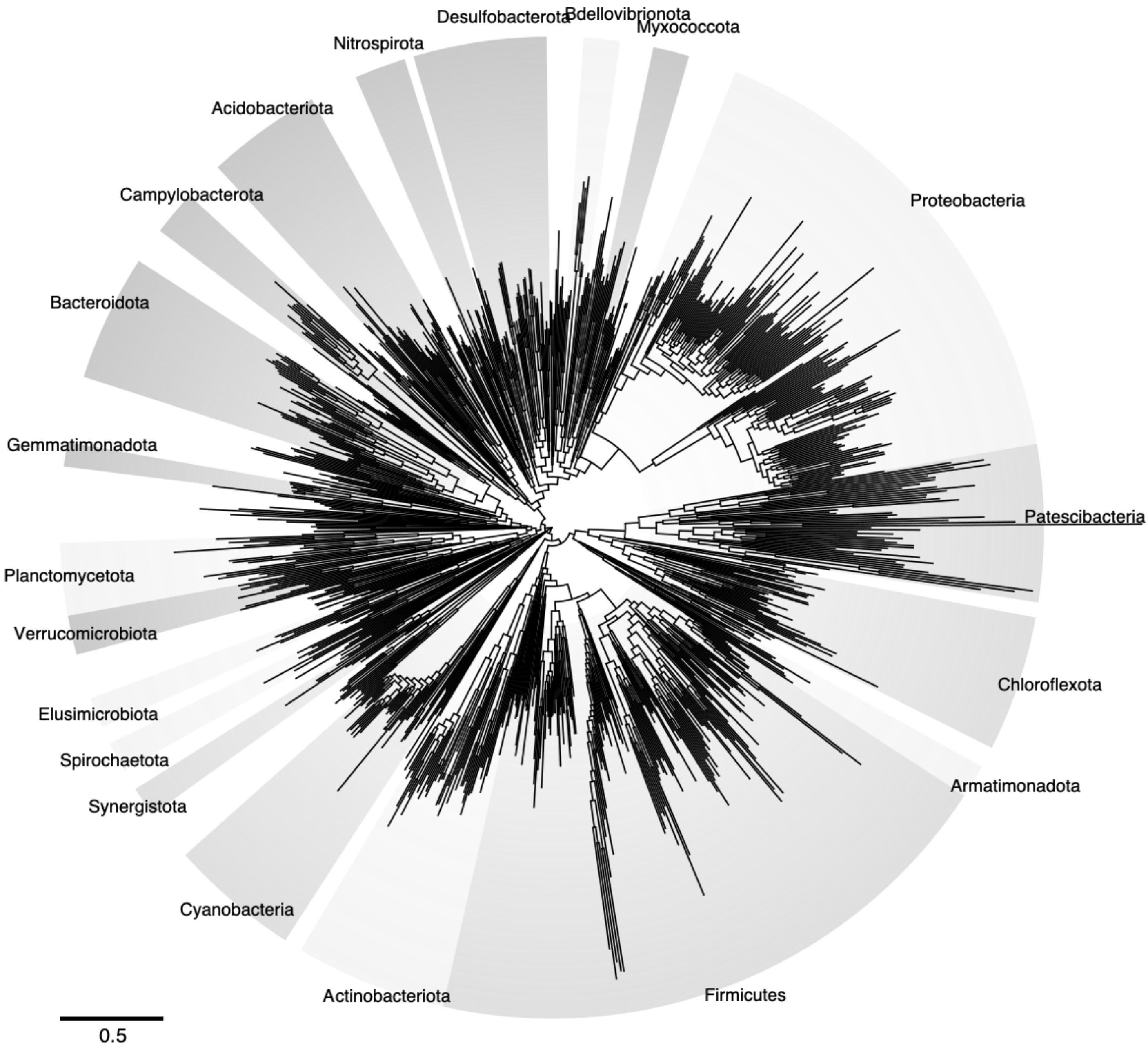


***Figure S1: Rooted Species Tree of Bacteria*** *including the longer branches that were removed for clarity in Figure 1 of the main text.*

#### Species tree inference after removing compositionally-biased sites from the concatenated alignment

Variation in nucleotide and amino acid composition across the tree has been shown to affect inferences of bacterial relationships, but substitution models that account for compositional heterogeneity across branches are computationally demanding on large datasets, such as the concatenate analysed here. As an alternative, we explored site-stripping approaches to evaluate the impact on the phylogeny of the most compositionally-biased sites. We ranked sites in the concatenation by the extent to which their composition differed between high-GC and low-GC taxa, using the approach of (Muñoz-Gómez et al. 2019), implemented in the Python script [https://github.com/Tancata/phylo/blob/master/rank_taxa_by_aa_compo.py](https://github.com/Tancata/phylo/blob/master/rank_taxa_by_aa_compo.py). We then deleted the top 50% most biased sites. A phylogeny inferred under the LG+EDM0064+G model (PMSF) was similar to the analysis on the full data. The phylogeny resulting from this analysis can be found in the Online Data Supplement with the name: `65genes_LGEDM0064LCLRG_50bh_PMSF`.

#### Adding mitochondrial and plastid genomes to the dataset

To leverage the fossil record of eukaryotes, we performed a molecular dating analysis in which we added the mitochondrial and plastid clades to the bacterial species tree, branching with Alphaproteobacteria and Cyanobacteria, respectively. To do so, we searched the genomes of 13 eukaryotes (*Arabidopsis thaliana*, *Capitella teleta*, *Limulus polyphemus*, *Lottia gigantea*, *Micromonas pusilla*, *Naegleria gruberi*, *Neurospora crassa*, *Paramecium tetraurelia*, *Physcomitrium patens*, *Porphyra umbilicalis*, *Priapulus caudatus*, *Selaginella moellendorffii*, *Tribolium castaneum*), and a representative sample of 38 archaea (representing: Heimdallarchaeota, Helarchaeota, Lokiarchaeota, Odinarchaeota, Thorarchaeota, Aigarchaeota, Bathyarchaeota, Crenarchaeota, Geothermarchaeota, Korarchaeota, Marsarchaeota, Nezhaarchaeota, Thaumarchaeota, Verstraetearchaeota, Euryarchaeota) for orthologues of the 65 marker genes using HMM (HMMER 3.3.2 (Eddy 2011), e-value of 1e-10) searching. Sequences found this way were added to the bacterial sequences before alignment through MAFFT v7.505 with mode L-INS-I (Katoh and Standley 2013), and subsequently trimmed using BMGE v1.12 (Criscuolo and Gribaldo 2010) with default settings aside from a BLOSUM30 matrix. We then performed iterative inference of single gene trees inferred with the options:

```
-m MFP -B 10000 -wbtl -madd
LG+C60+F+G,LG+C50+F+G,LG+C40+F+G,LG+C30+F+G,LG+C20+F+G,LG+C10+F+G,
LG+F+G,LG+R+F --score-diff ALL
```

to identify the orthologous (mitochondrial- or plastid-origin) eukaryotic genes in each case. These alignments were then concatenated and used to infer a species tree using the PMSF LG+C20+F+G model in IQ-TREE 2, with a guide tree inferred using LG+F+G), with 1000 ultrafast bootstrap replicates. The files underlying this analysis can be found in the Online Data Supplement with the name `PlastidAnalyses.tar.gz`.

### 2. Gene tree-species tree reconciliation pipeline

#### Gene family alignments

We used EggNOG-mapper (Huerta-Cepas et al. 2017) to annotate gene models on the 1007 bacterial genomes with the following flags '`-m diamond -i AllProteins.faa --query_cover 50.0 --evalue 0.0000001 --cpu 24 -o dimnd_mega_harsh_no_o2o`'. The resulting table can be found in Online Data Supplement with the name `dimnd_mega_harsh_no_o2o.emapper.annotations`. We also obtained the KO annotations using the table HMMER3/f [3.1b2 | February 2015]. In every COG cluster inferred by EggNOG, we included only those sequences whose KO annotation was present and equal to the most common KO annotation within that cluster. We assign a KO to a sequence only if the value is < 0.01 and we only assign one annotation to every gene, the one with the lowest e-value and the highest bitscore. The KO annotations can be found in the table Annotation_KEGGs.tsv. The command used was:

```
hmmsearch --tblout KEGG/sequence_results.txt -o KEGG/results_all.txt --domtblout KEGG/domain_results.txt --notextw --cpu 58 ./KEGG/All_KOs.hmm AllProteins.faa
```

The resulting sequences grouped into different COG clusters (3500) can be found in Online Data Supplement with the name `Fastas1.zip`.

#### Alignment preparation

From this initial clustering, we performed an iterative filtering process to retain full-length homologous sequences amenable to phylogenetic analysis and subsequent gene tree-species tree reconciliation. We started obtaining alignments using MAFFT (auto). An initial visual inspection of the alignments revealed that some genes that appear to have multiple copies in a single genome, would sometimes show a pattern suggestive of actually belonging to the same single tree that had been split into different genes, potentially due to the genes being present in different contigs or to an incorrect inference in the gene calling step. We developed a small pipeline to automatically identify and correct these cases in which the gene has been fragmented into pieces.

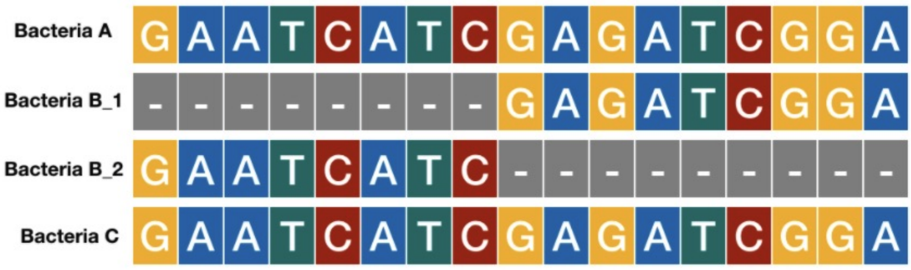


***Figure S2: Identified genes split into fragments.*** *In the example 4 sequences are compared. Bacteria B has two genes, with a pattern of amino acid presence/absence that is suggestive of some error in the annotation pipeline that has artifactually split a homologous sequence into two different ones.*

We use a custom made script in Python named fragment_detector.py (Online Data Supplement) that iterates the columns of the alignment one by one, making all the pairwise comparisons between multicopy genes, assigning a score to those sequences. The score is obtained by adding one every time two columns from the sequences compared contain one gap and one amino acid, and subtracting 5 in the case that the two sequences have both amino acids. The resulting score is divided by the length of the two sequences. Some manual iterations led us to use a threshold of 80% to successfully identify the sequences that we could clearly recognize as different fragments of the same gene. As an additional quality control, we determined the position of every pair of possible fragments within their contigs and selected only those fragments that were, either contiguous in the genome, either in different contigs and at the extreme of them. Once we identified the fragments to stitch, we used a script in Python named stitcher.py (Online Data Supplement) to put together the fragments. We recomputed the alignments of all the clustering with the new corrected sequences, using MAFFT linsi with the command `mafft --thread -1 --maxiterate 1000 --localpair` and trimming the sequences with BMGE (BLOSUM 30). We filter from these alignments those sequences with an occupancy lower than 50%. The alignments can be found in the Online Data Supplement with the name: `FilteredFinal2.zip`.

#### Inference of bacteria-specific site-heterogeneous substitution models

We used EDCluster (https://github.com/dschrempf/EDCluster, (Schrempf, Lartillot, and Szöllösi 2020)) to infer a dataset-specific site-heterogeneous substitution model for inferring bacterial gene family trees. To do so, we randomly sampled 12% of the gene families in the dataset, then discarded the largest and smallest 1% of the sample to give 10% of the families with intermediate size. This first sample can be found in the Online Data Supplement with the name `RandomSample1ED.zip`. For these 334 families, we inferred ML trees under the LG+G model (iqtree 2.1.3: `iqtree2 -s $alignment -m LG+G`). The results of this analysis can be in the Online Data Supplement with the name randomsample_LG.tar.gz.

These trees were then used as fixed guide trees to sample site profiles for each alignment under the CAT+Poisson model in Phylobayes (pb 4.1b: `pb -cat -poisson -d $alignment -T $iqtree_treefile chainname`). Two chains were run for each pair of alignment and guide topology until at least 80000 samples were collected. Only the first chain of those runs were considered in the further analysis, where proper convergence was reached for the posterior log-likelihood (effective sample size above 50 and relative difference between the chains smaller than 0.3). This left 314 families. For these families the posterior site profile was exported from the first chain using the command:

```
readpb -ss -x $20_percent_burnin 1 chainname.
```

The site profiles are the input for the program EDCluster which generates all the mixture models with cluster numbers from 4 to 4096 and transformations none, CLR, and LCLR

(EDCluster *.sitestat). All the relevant files can be found in the Online Data Supplement in the file named `EDClusters.tar.gz`.

#### Comparison of model fit

We used the model-testing mode in IQ-TREE 2.1.1 to evaluate the fit of the dataset-specific EDM models to all the alignments using the command (with COG0002 as an example):

```
iqtree2 -s COG0002.faa.aln.trimmed.filtered.faa.aln.stitched.trimmed.filtered -mdef models/128_models.nex -m TESTONLY -mset LG -madd Poisson+EDM0008LCLR+G,Poisson+EDM0016LCLR+G,Poisson+EDM0032LCLR+G,Poisson+EDM0064LCLR+G,Poisson+EDM0128LCLR+G,LG+EDM0008LCLR+G,LG+EDM0016LCLR+G,LG+EDM0032LCLR+G,LG+EDM0064LCLR+G,LG+EDM0128LCLR+G,C20+G,C40+G,C60+G,LG+C20+G,LG+C40+G,LG+C60+G --score-diff ALL
```

LG+EDM models were the most popular overall (2564/3064 alignments), with LG+EDM0128+G the most popular model (2557). Poisson+EDM models were the next most popular choice (250 alignments). LG+C60+G (52 alignments) and variants on the site-homogeneous LG model (LG+G, 50; LG+F, 31; LG+I, 28 alignments) were also occasional choices. As the mixture weights were inferred from the sample of 314 alignments, the number of parameters did not differ between EDM models with different numbers of mixture components. BIC scores increased consistently with the number of mixture components up to 128 components, beyond which model fitting was impractical due to RAM limitations.

#### Refinement of the alignments based on the EDM trees

Upon a visual inspection of the trees obtained in the previous step, we decided to further improve the pipeline to avoid the inference of very long branches (>= 2 mean substitutions per site). We decided to “cut” the phylogenetic trees by removing those branches, which could result in the total tree divided into different subclusters if the cut branch was an internal one. We used two custom made scripts that can be found in the Online Data Supplement (cut_gene_tree.py, create_cluster_fastas.py). This generated a total of 5051 subclusters. We decided to name the clusters using the following schema. If the original family COGXXX was broken into n subclusters, they would be ordered from the largest (in terms of number of sequences) to the smallest, and named: COGXXX_0, COGXXX_1, COG_(n-1). Gene clusters that were not subdivided were named with the name of the original COG family plus “_X”. All the relevant files here, including the trees that were cut into pieces, the fasta of the original clusters and the fasta files of the subclusters can be found in the Online Data Supplement with the name `SubclusterProcessing.zip`.

#### Reprocessing of subclusters

The resulting fasta files were again realigned with MAFFT (linsi), trimmed with BMGE (BLOSUM30) and filtered again to remove those sequences with 50% or less of amino acid occupancy. The final alignments that were used to infer the gene trees that were reconciled with the species tree can be found in the Online Data Supplement with the name `FinalAlignments.zip`.

#### Inference of subcluster's gene trees

We use the same procedure as before,

```
iqtree2 -s ./supercurated/COG0002.faa.aln.trimmed.filtered.faa.aln.stitched.trimmed.filtered -mdef /scratch/tw15962/projectB/edm/models/all_models.nex -m TESTONLY -mset LG -madd Poisson+EDM0008LCLR+G,Poisson+EDM0016LCLR+G,Poisson+EDM0032LCLR+G,Poisson+EDM0064LCLR+G,Poisson+EDM0128LCLR+G,LG+EDM0008LCLR+G,LG+EDM0016LCLR+G,LG+EDM0032LCLR+G,LG+EDM0064LCLR+G,LG+EDM0128LCLR+G,C20+G,C40+G,C60+G,LG+C20+G,LG+C40+G,LG+C60+G --score-diff ALL
```

Then, we compute the guide tree

```
iqtree2 -s ${JOB_ARRAY_FILE} -m MFP --prefix ${JOB_ARRAY_FILE}_guide
```

Finally, we would launch the command

```
iqtree2 -s COG0002_0.faa.aln.trimmed.filtered.faa.aln.stitched.trimmed.filtered.treefile.subclusters2.faa.aln.stitched.trimmed.filtered -mdef

edm/models/all_models.nex -m LG+EDM0128LCLR+G -ft COG00002_0.faa.aln.trimmed.filtered.faa.aln.stitched.trimmed.filtered.treefile.subclusters2.faa.aln.stitched.trimmed.filtered_guide.treefile -pers 0.2 -nstop 500 -nm 100000 -B 10000 -wbtl -
```

#### Reconciliations with ALE

The basic commands used to perform gene tree-species tree reconciliations using ALE (Szöllõsi et al. 2013) were:

```
ALEobserve COGXXXX.ufboot

ALEml_undated species.tre COGXXXX.ale fraction_missing=fm.txt
```

The fraction missing file can be found in the Online Data Supplement with the name: `fraction_missing.txt`.

### 3. Phenotype Prediction

#### Prediction via gene content

Inspired by some previous approaches (Weimann et al. 2016; Jabłońska and Tawfik 2019), we developed a machine learning classifier to predict whether genomes are aerobic or anaerobic. The workflow code is available at: https://github.com/wwood/bacterial_dating_aerobic_predictor.

##### Gold standard dataset collection

A collection of 3184 tuples of (GTDB_genome_accession, growth in oxygen) was collected, where a genome is defined as a GTDB r202 accession, and "growth in oxygen" is defined as either aerobic or anaerobic (Online Data Supplement, "`AerobicPredictionGoldStandard`"). Of these, 2129 were annotated aerobic, and 1055 annotated anaerobic.

The majority of the dataset was derived from BacDive, accessed March 15, 2023 (Reimer et al. 2022). GTDB 06-RS202 genus representatives (metadata column "gtdb_type_species_of_genus") were selected, including both Bacteria and Archaea. Of those, the genus of the "ncbi_taxonomy" column was parsed out. These genus names were used to search BacDive through the API searching by "taxonomy" (https://pypi.org/project/bacdive/), accessed March 15, 2023. For each species returned by that query, the "oxygen tolerance" entries were tabulated. This list of species and growth characteristics, where the species was also a GTDB species representative, and was the originally described species (i.e. did not have a postfix _A, _B, etc), was further condensed. First, "anaerobe" and "obligate anaerobe" were classed as "anaerobe", while "aerobe" and "obligate aerobe" were classed as "aerobe". Species annotated as "microaerophile", "facultative aerobe", "facultative anaerobe", "aerotolerant" and "microaerotolerant" were excluded. Species classed as both aerobe and anaerobe were also excluded. Second, to balance the species representation across the tree, a maximum of three species were chosen at random from each genus, but including the type species of the genus where possible.

Cyanobacteria are not included in BacDive, so were incorporated through a separate procedure. From the class c__Cyanobacteriia, also known as Oxyphotobacteria, one genome from each genus was chosen with CheckM completeness 95% and the contamination < 5%. At most 10 were chosen from each family. Together with *Vampirovibrio chlorellavorus*, which has been cultured (Coder 1981), and each of the c__Cyanobacteriia from the 1007 genomes included in the species tree here, all were included in the dataset as aerobic.

Before training, 20% of the data was excluded to be used as a test set. To increase confidence that accuracy observed in the test set is representative of the accuracy of the predictors in lineages outside of the training set, and accuracy on reconstructed ancestral states, genomes from each family were included in either the test set, or the set used for training / cross validation.

To determine the gene families present in each genome, the amino acid sequences of genes were annotated using eggNOG-mapper v2.1.3 (Cantalapiedra et al. 2021) and HMMSEARCH v3.3.2 (Finn, Clements, and Eddy 2011) against KOFAM 2022-01-30 (Aramaki et al. 2020), using the following commands:

```
emapper.py -m diamond --target_orthologs one2one --query_cover 50.0 --evalue 0.0000001 --cpu 24
```

```
hmmsearch --tblout ... kofam/2022-01-30/profiles.hmm
```

The KEGG orthologous group with the lowest e-value and the root eggNOG-mapper annotations were collected for each gene. Only those pairs in a predetermined whitelist were considered as features for the prediction (3507 genes, Online Data Supplement, “ModalKEGGs.tsv”). This whitelist only contained annotations considered suitable for the ancestral state reconstruction. During training and testing, only entire gene families (as opposed to subclusters, as above), were considered.

To increase robustness of the predictor, each genome was included multiple times in the training and test datasets after adding gene annotations (simulating “contamination” in the original genomes or extra genes being erroneously included as part of the ancestral state reconstruction algorithm) and/or removing gene annotations (simulating “incompleteness” in the original genomes or inferred ancestral states). Specifically, 0%, 10%, 20%, 30%, 40% and 50% of genes were removed at random (the “gene removal rate”). Then, 0%, 10%, 20%, 30%, 40% and 50% extra genes were included (the “extra genes rate”), chosen from the set of all genes. Note that this percentage was a percentage of the number of genes included in the ancestral reconstruction, not a percentage of the genes in each genome. Selection of genes to add was based on a weighted random sampling, where the weights were the total copy number of each gene amongst the sum of all genes’ copy numbers. In total, 88,164 entries were used for training / cross-validation.

##### Machine learning model training and benchmarking

Several machine learning algorithms were applied to this training set, assessed firstly using 5-fold cross validation: XGBoost (Chen and Guestrin 2016) as well as RandomForestClassifier, GradientBoostingClassifier, AdaBoostClassifier, ExtraTreesClassifier, LogisticRegression, and Perceptron classifiers from scikit-learn ( version 1.2.2, (Kramer 2016)). The input to the model was copy numbers of each gene. Before training, a MaxAbsScaler scaling was performed, so that all inputs were valued between 0 and 1. RandomForestClassifier and ExtraTreesClassifier algorithms used n_estimators=1000. The GradientBoostingClassifier had learning_rate=0.1. As with the partitioning of the training and test sets above, genomes from the same taxonomic family were only included in either the training partition of the cross validation or the validation, implemented using sci-kit learn’s GroupKFold. Final predictors were trained using the entirety of the training dataset.

SHapley Additive exPlanation (SHAP) values (Lundberg et al. 2020) were used to indicate the contribution each gene family made to the overall prediction in these datasets (Online Data Supplement, “`xgboost_shap_values.csv`”). Amongst the 20 genes which contributed most to the the predictions according to these SHAP values, some were more prevalent in aerobes, while some were more prevalent in anaerobes (**Figure S4**).

Plotting (using ggplot2, (Wickham 2016)) the accuracy scores of each of the algorithms with increasing quantities of noise in their inputs (either addition of extra genes or removal of genes encoded in the genome) showed most algorithms were robust (**Figure S3**). XGBoost showed the highest accuracy (96.6%), where accuracy was defined as (true_positive+true_negative) / (true_positive+true_negative+false_positive+false_negative) amongst the entire training dataset (**Table S1**).

| **Predictor** | **Cross-validation accuracy (%)** | **Hold-out (test) dataset accuracy** |
|---|---|---|
| XGBoost | 96.6 | 97.2 |
| LogisticRegression | 96.0 | 97.1 |
| GradientBoosting | 95.4 | 96.8 |
| Perceptron | 95.3 | 97.0 |
| RandomForest | 94.7 | 97.0 |
| AdaBoostClassifier | 94.2 | 94.6 |
| ExtraTrees | 94.0 | 90.7 |

***Table S1****. Accuracy of trained models assessed through cross validation and by the hold-out test dataset.*

After training each algorithm on the full training dataset, they were applied to the ancestral states and leaf nodes of the species tree (Online Data Supplement, "`gene_based_prediction_on_ancestors_and_leaves.csv`"). The input dataset was the gene family copy number (which, for ancestral nodes, was inferred using gene tree-species tree reconciliation) at each node, which for ancestral states was a fractional value rather than a whole number. In cases where gene families were broken into subclusters for ancestral state reconstruction, the copy numbers of each subfamily were summed to estimate the copy number of the entire family.

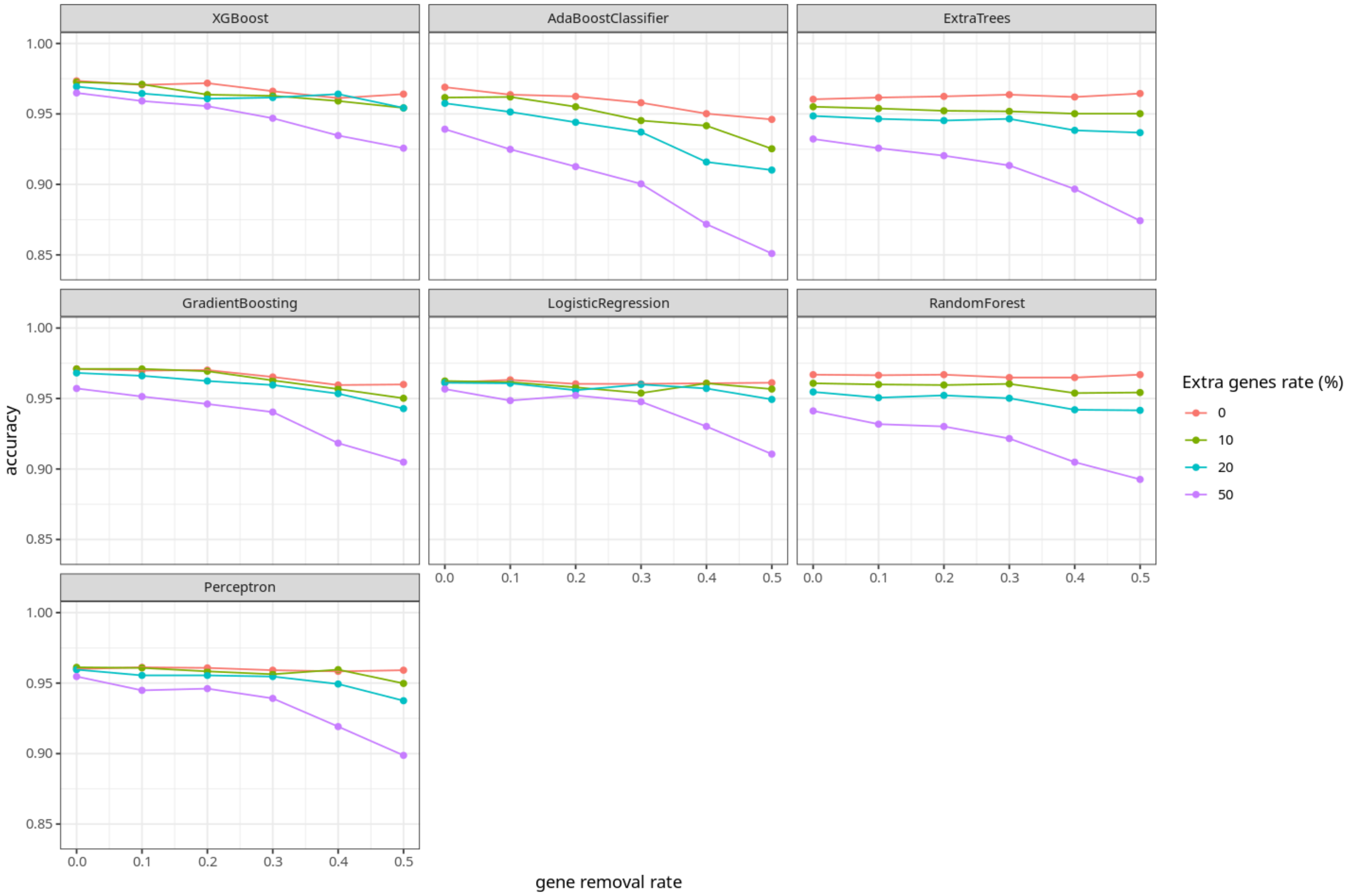


***Figure S3**: **Accuracy of trained machine learning models assessed using cross-validation.** Points show the average accuracy obtained for data collected (left-most red points) or simulated (all other points). Simulated data was generated by adding randomly chosen genes (coloured lines, "extra genes rate") or removing genes at random (x-axis, "gene removal rate") from each model's inputs. Given its robustness to noise in its inputs and overall best accuracy (**Table S1**), we prioritised the use of the XGBoost model in subsequent analyses.*

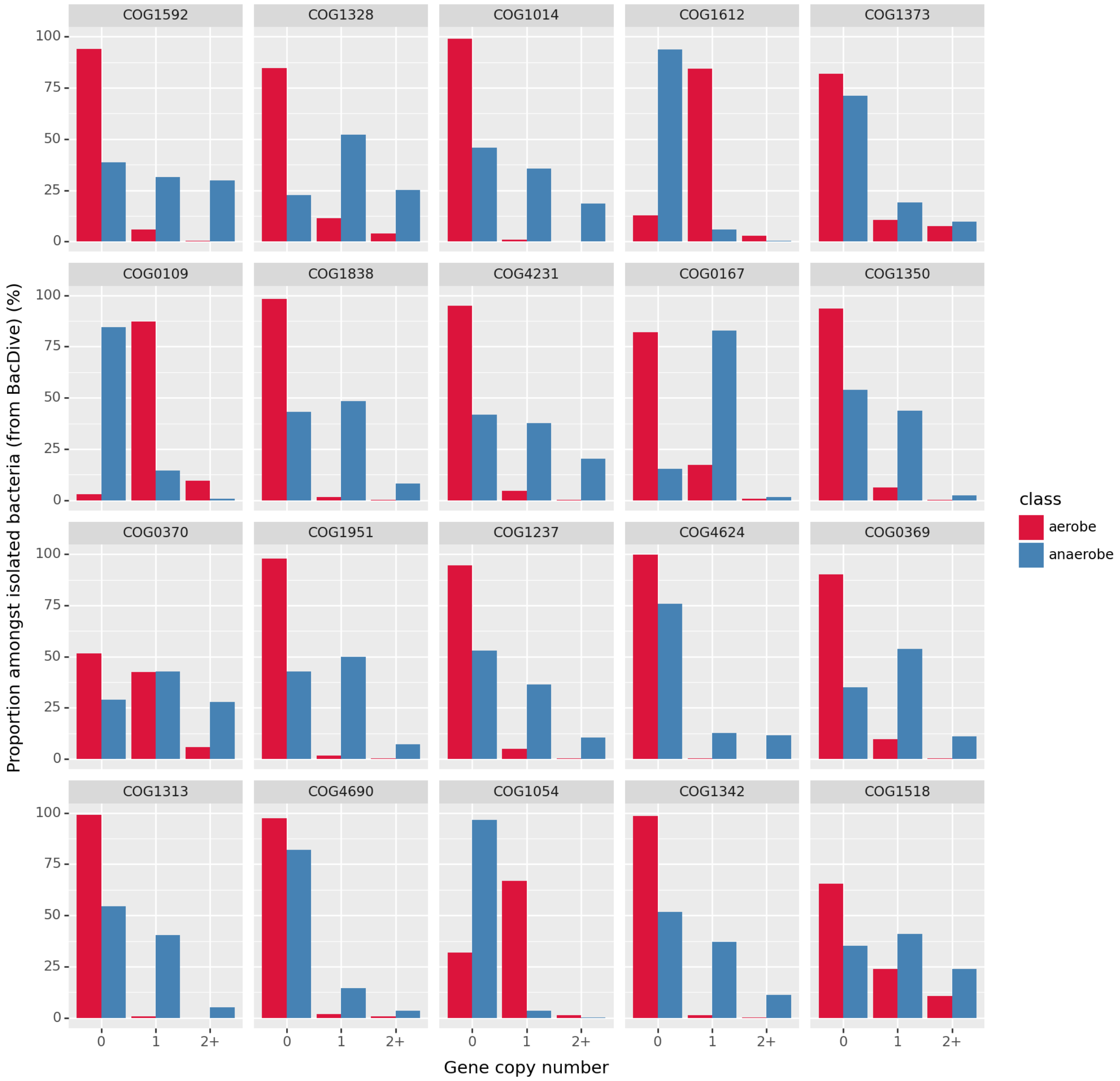


***Figure S4**: **Copy numbers of genes with large contributions to aerobicity prediction.** The gene families with the highest absolute SHAP values are plotted, with counts amongst aerobic (red) and anaerobic (blue) genomes in the gold standard shown. These counts show that no single gene family is perfectly predictive of aerobicity.*

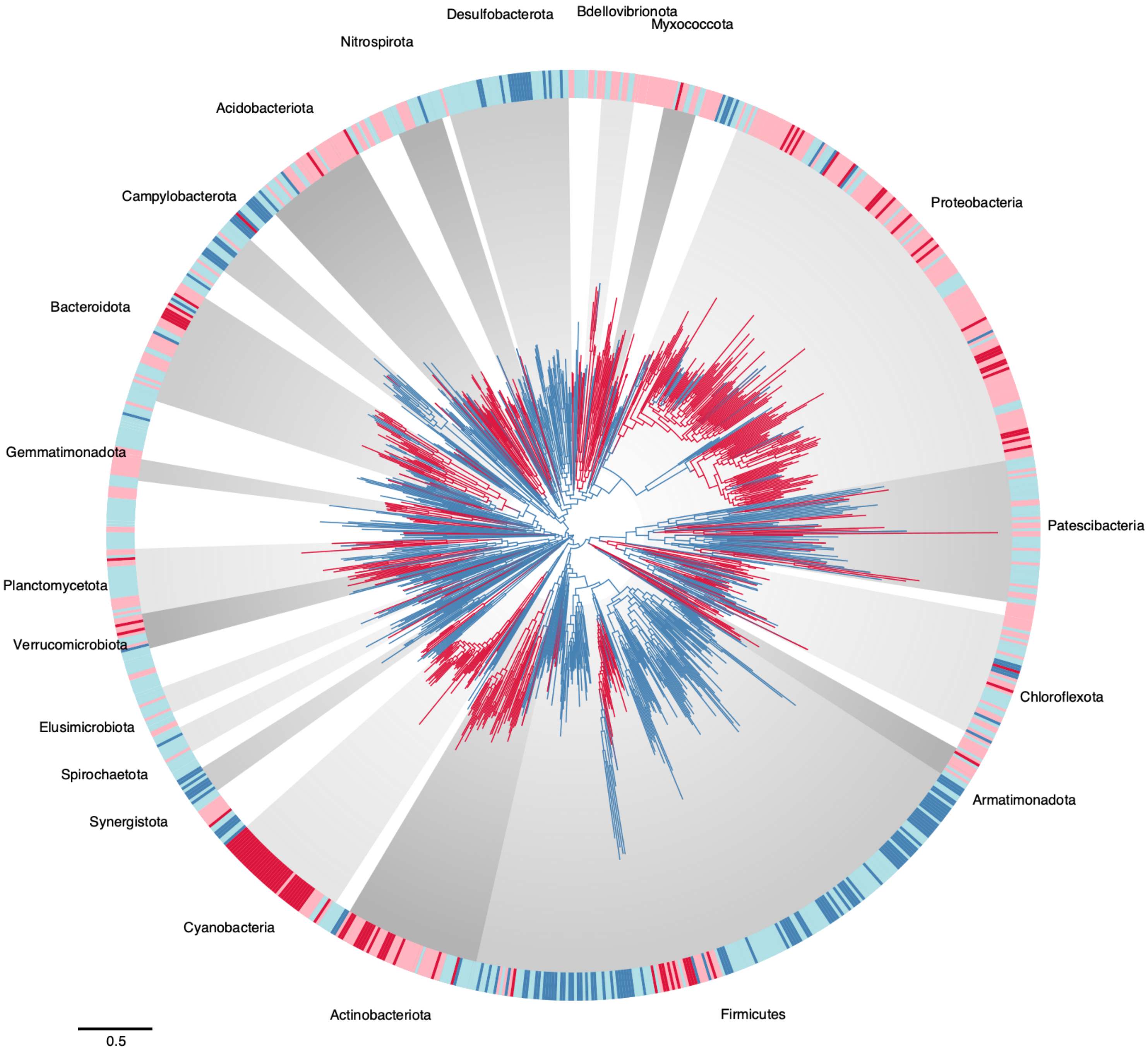


***Figure S5 Reconstruction of aerobic metabolism.*** *Tree from Fig. S1 with branches coloured according to aerobic predictions according to XGBoost and including long branches omitted in Fig 1 of the main text.*

#### Comparison with predictions made by the presence of heme-copper oxygen reductases (HCO) and cytochrome bd oxygen reductases (cytbd)

We considered a rule-based predictor which equates the presence of any of the two terminal respiratory oxygen reductases in bacteria (heme-copper oxygen reductase (HCO) and cytochrome bd oxygen reductase (cytbd)) with a prediction of "aerobic" and lack of HCO/cytbd genes with a prediction of "anaerobic". Alternative oxidases (AOX) are also found in bacteria but these enzymes are very rare, and do not contribute to the generation of proton motive force (Dunn 2023) so we did not include them in our analysis. The HCO and cytbd dataset was manually curated to ensure that genes were allocated to the correct subfamily, which can be challenging for this large protein family (OnlineData Supplement, "`aerobic_respiration_by_species_in_gtdb_r202.tsv`"). Manual curation was performed by extracting all HCO and cytbd sequences from our dataset of 1007 species with HMMER v3 and a curated database of HMMs available on GitHub (https://github.com/ranjani-m/cytbd-superfamily, https://github.com/ranjani-m/HCO) (Murali, Gennis, and Hemp 2021; Murali, Hemp, and Gennis 2022). Thereafter, sequences classified into different families were verified by manual inspection, and sequences that did not contain conserved active site features were removed, as previously described (Murali, Gennis, and Hemp 2021; Murali, Hemp, and Gennis 2022) to avoid mis-annotation of $O_2$ reduction function.

We found that predictions based solely on the presence of HCO and cytbd were not as accurate as those made by integrating signals across many genes as above. The reasons for the comparatively poor accuracy of this rule-based predictor are two-fold. Firstly, it will likely perform poorly when the input set of genes is not entirely reliable (i.e. when the extra genes rate or gene removal rate is non-zero), since the prediction is made on a very small number of genes. Secondly, we found that a substantial fraction of species whose growth condition was measured as being "anaerobic" in BacDive encoded at least one predicted HCO or cytbd $O_2$ reductase (448 of 1055, 42% of anaerobic species in the training dataset were in this category). Aerobic species, meanwhile, almost uniformly encoded at least one HCO or cytbd $O_2$ reductase (2123 of 2129, 99.7%). The most frequent $O_2$ reductases encoded in genomes marked as anaerobic were HCO A, C and qOR1, though other subclasses are also represented (**Table S2**). The implication of the observation of $O_2$ reductases in genomes annotated as anaerobic isn't clear. In some cases, these enzymes may simply never be used or turned on. It is also possible that the experimental observations in BacDive do not include conditions that reflect every type of ecological niche where $O_2$ reductases are needed. For e.g., while BacDive lists the oxygen tolerance of *Desulfovibrio vulgaris* as "anaerobe", it has been shown to reduce oxygen to water, uncoupled to growth (Ramel et al. 2015, PLoS ONE). However, the BacDive experimental data likely reflects the number of species whose lifestyles are primarily "aerobic" and we think there is sufficient statistical resolving power here for us to infer the evolution of aerobic lineages and constrain the bacterial timetree with the coevolution of bacteria and aerobic metabolism. Despite the less accurate predictions made by observation of terminal $O_2$ reductases in each genome, $O_2$ reductases (and HCO enzymes in particular) nonetheless provide some predictive utility, with several ranking highly in the SHAP analysis of the XGBoost predictor (**Figure 1,** Online Data Supplement, "`xgboost_shap_values.csv`").

| Oxygen reductase | Number of anaerobic genomes in gold standard that encode at least one copy |
|---|---|
| A (HCO) | 107 |
| B1 (HCO) | 12 |
| B3 (HCO) | 3 |
| B4 (HCO) | 1 |
| B5 (HCO) | 1 |
| B6 (HCO) | 1 |
| C (HCO) | 100 |
| OR-C1a (cytbd) | 77 |
| OR-N3a (cytbd) | 1 |
| qOR1 (cytbd) | 400 |
| qOR2 (cytbd) | 6 |
| qOR3 (cytbd) | 7 |
| qOR4a (cytbd) | 14 |

***Table S2.*** *Counts of anaerobic species containing HCO and cytbd genes.*

#### Aerobic prediction in the node immediately ancestral to the classes Cyanobacteriia (Oxyphotobacteria) and Vampirovibrionia

The geological isotope record contains signatures stable of oxidized Fe and U-Th-Pb isotopes at 3.2Ga that have been interpreted as evidence of free oxygen consistent with oxygenic photosynthesis by stem Cyanobacteria (Satkoski et al. 2015; Betts et al. 2018). This evidence constrains the common ancestor of Cyanobacteriia and their closest relatives, Vampirovibrionia, to be at least this old. Interestingly, this ancestral node is robustly predicted to be aerobic. This is despite the fact that extant Cyanobacteriia and Vampirovibrionia harbour distinct sets of genes for aerobic respiration, which was previously interpreted as evidence that aerobic respiration evolved following the diversification of Cyanobacteriota classes (Soo et al. 2017). The aerobic prediction is based on the inference that some key genes of aerobic metabolism including components of the cytochrome C complex and heme synthase genes, were already present at this node. The analysis therefore suggests that the ancestor of Vampirovibrionia and Cyanobacteriia was already capable of aerobic metabolism.

To investigate the aerobic prediction for the common ancestor of Cyanobacteriia and Vampirovibrionia (which we refer to below as node 1963, in line with the numbering in our species tree), we first inspected the SHAP values to ascertain which genes underlie the aerobic prediction, finding in particular that near-zero inferred copy number of ribonucleoside-triphosphate reductase (formate), CRISPR-associated protein Cas1 pushed the prediction towards “aerobic”, alongside the presence of cytochrome c oxidase subunit I and heme a synthase genes. In contrast, lack of heme o synthase pushed the prediction “anaerobic”.

The copy number of each gene inferred through ancestral reconstruction is not a whole number, but rather encodes an estimate of that gene’s true copy number. To interpret these numbers more directly as a probability, we modelled the presence of each gene as a binomial distribution, and sampled from these distributions for each gene, arriving at a copy number for each gene. Estimated copy numbers greater than 1 were modelled as having the copy number of the estimate’s floor, plus the remainder treated as a probability in the same way. Application of the predictors to these copy numbers then yielded a prediction of its aerobicity. This process was repeated 100 times, noting the number of times the prediction was aerobic vs anaerobic. For node 1963, 97 out of 100 iterations were predicted “aerobic”, providing further evidence of this node being aerobic.

The second test was motivated by the observation that all cyanobacteria in the gold standard are marked as aerobic. We therefore reasoned that perhaps the predictor was actually inferring that node 1963 was cyanobacterial, and therefore aerobic, rather than arriving at an aerobic prediction without regard to the node’s taxonomy. To exclude this possibility, we remade the predictors using the same dataset as previously, except removing all Cyanobacteria from the gold standard. Most predictors including XGBoost maintained their prediction of the node being aerobic, albeit with lower confidence (**Table S3**).

Thirdly, we tested whether the generalised bias towards lower copy numbers in more ancient nodes could be underlying the aerobic prediction. Estimated copy numbers at each node were artificially increased by values in the range 0-0.5 in increments of 0.05, and the predictors rerun each time. In all cases, the XGBoost predictor predicted the node to be aerobic. Given the results of these more detailed analyses, we conclude that node 1963, the last common ancestor of Cyanobacteriia (Oxyphotobacteria) and Vampirovibrionia, likely performed aerobic respiration. Overall, we interpret these findings to indicate that oxygenic photosynthesis and aerobic respiration evolved close together in time early in the evolution of Cyanobacteriota.

It is interesting to note that, at 2.5-2 Ga, the common ancestor of Cyanobacteriia is the oldest node on the species tree to which any kind of photosynthesis can be mapped confidently, with anaerobic photosynthesizers likely having acquired their capabilities more recently (Figure 3, see also (Oliver et al. 2021)). Therefore, if the ancestral form of photosynthesis is anaerobic, this metabolism provides a good example of the “It’s the song, not the singer” model of evolution in which metabolic capabilities may outlast the organismal lineages that originally invented them (Doolittle and Booth 2017).

| **Model** | **Dataset without Cyanobacteria** | | **Dataset with Cyanobacteria** | |
|---|---|---|---|---|
| | **prediction** | **probability** | **prediction** | **probability** |
| XGBoost | aerobe | 0.86 | aerobe | 0.97 |
| LogisticRegression | anaerobe | 0.02 | aerobe | 0.71 |
| GradientBoosting | aerobe | 0.77 | aerobe | 0.86 |
| Perceptron | anaerobe | N/A | aerobe | N/A |
| RandomForest | aerobe | 0.69 | aerobe | 0.80 |
| AdaBoostClassifier | anaerobe | 0.49 | aerobe | 0.51 |
| ExtraTrees | aerobe | 0.56 | aerobe | 0.69 |
| GaussianNB | anaerobe | 0.00 | anaerobe | 0.00 |

***Supplementary Table S3****. Predictions of aerobicity in node 1963 when the cyanobacteria were removed from the gold standard.*

#### Aerobicity prediction using a concatenate of marker genes

Our hypothesis is that some amino acids at specific positions might be indicative of an aerobic or anaerobic lifestyle. We first developed a classifier based on site-wise amino acid state in a concatenate of 1007 sequences and 14075 amino acid sites. Second, we reconstructed ancestral sequences along the species phylogeny. Third, we predicted whether ancestral nodes were aerobic or anaerobic by applying our classifier on the reconstructed ancestral sequences.

**Step 1:** Out of the 1007 species, we obtained aerobic or anaerobic lifestyle annotations for 333 species, 145 of which were aerobes, and 188 anaerobes. We performed logistic regression on site-wise amino acid states based on the sequences of these 333 species. The model can be written as follows:

$$\log\left(\frac{p(\text{aerobe})}{p(\text{anaerobe})}\right) = \alpha_0 + \sum_{\text{site } i} \sum_{\text{state } j \text{ at site } i} \alpha_{ij}\delta_{ij}$$

Each site $i$ with $k$ states j contributes $k-1$ variables to the model. $\alpha_0$ is the intercept, and $\alpha_{ij}$ is the parameter associated with observing state $j$ at site $i$. We use lasso penalization to control the number of parameters, as implemented in glmnet, and cross-validation to choose the regularization parameter $\lambda$ that minimizes classification error (named lambda.min in glmnet). The resulting model has 99 non-zero parameters.

**Step 2:** Ancestral sequences were reconstructed using IQTREE 2 (Minh et al. 2020) along a tree with model LG+F+G4, as follows:

```
iqtree2 -s aln/Concatenate.faa -nt 32 -asr -m LG+F+G -te SpeciesTree.nwk -safe -pre lgfg
```

**Step 3:** , the Ancestral nodes were classified with the classifier according to their reconstructed ancestral sequences.

### 4. Calibrations

In formulating the calibrations, we sought to follow the best practice principles set out in (Parham et al. 2012). However, these were designed with animal and plant fossil-based calibrations and not all of the principles are applicable to calibrations of microbial clades which often lack phenotypic synapomorphies, let alone diagnostic characters that are preserved in fossil remains. Furthermore, the calibrations for many clades rely on geochemical evidence of microbial metabolisms, manifest as isotope fractionation or oxidation states of redox sensitive mineral species. Consequently, we have adapted the best practice principles to suit the nature of the calibrations. Novel calibrations are justified in full; we indicate the source of calibrations that are justified elsewhere, providing notes where they have been adapted for different clades or where the dating has changed subtly with the revision of the geologic timescale.

**LUCA | 3347-4520 Ma**
**Fossil taxon and specimen:** Strelley Pool Formation, Pilbara Craton, Following the justification outlined in (Betts et al. 2018).
**Minimum age justification:** The minimum age of the Strelley Pool Formation is 3.350 Ga ± 0.003 Gyr based on a volcaniclastic tuff, at the base of the overlying Euro Basalt (Nelson, n.d.) in the Kelly Group. Hence our minimum age constraint is 3.347 Ga.
**Hard maximum age justification:** The Moon forming impact would have effectively sterilized the Earth and so it serves as an effective basis for establishing a hard maximum age constraint on LUCA. Pb-Pb dating carried out on Moon rocks, yielding a date of 4.51 Ga ± 10 Myr (Hanan and Tilton 1987) a date which has also recently been confirmed by reanalysis of the Apollo mission zircons (Barboni et al. 2017). Thus, our maximum constraint is 4.52 Ga.

**Total group Cyanobacteria | 3225-4520 Ma**
**Clade:** This equates to the traditional concept of total group Cyanobacteria, excluding Melainabacteria, Vampiriovibrionia and Sericytochromatia (Shih et al. 2017), but including the stem to this remaining clade.
**Fossil taxon and specimen:** stable Fe and U-Th-Pb isotopes in the Manzimnyama Banded Ironstone Formation, Fig Tree Group, Barberton, South Africa (Satkoski et al. 2015).
**Hard minimum age:** 3225 Ma
**Hard maximum age:** 4520 Ma
**Discussion:** there are claims and counterclaims for life in the Hadean and Archaean and while we accept that there is credible fossil, sedimentologic and isotopic evidence for life by 3347 Ma (see (Betts et al. 2018), there has been insufficient consideration of whether these records evidence the establishment of the crown clade of life - i.e. descent from LUCA. Implicitly or explicitly, most records have been attributed to the crown clade of life based on claims, direct or indirect, of oxygenic cyanobacteria, either as microfossils, sedimentary structures such as MISS or stromatolites, or Banded Ironstones. The existence of stromatolites would appear to provide evidence of phototactic bacteria, but they might otherwise represent microbial organisms competing for other nutrients within the water column (Bosak, Knoll, and Petroff 2013). Similarly, banded ironstones can be formed by reaction with oxygen from abiogenic sources like photolysis (Cairns-Smith 1978), hence, banded ironstones only become a significant proxy for life when they occur in volume (and

even then they are not necessarily linked to oxygenic photosynthesis). Therefore we follow Betts et al. (2018) who based their calibration minimum for total-group cyanobacteria on Satkoski et al. (2015) who presented evidence of stable Fe and U-Th-Pb isotopes in the Manzimnyama Banded Ironstone Formation (Fig Tree Group, Barberton, South Africa), indicating a level of free oxygen indicative of cyanobacterial activity (Betts et al. 2018; Satkoski et al. 2015).
**Hard minimum age justification:** (from Betts et al. 2018) The isotopic evidence from the Manzimnyama BIF in the Fig Tree Group, Barberton, South Africa (Satkoski et al. 2015). The age of the Fig Tree Group is well constrained with a spherule layer at its base dated at 3258 Ma ± 3 Myr (Byerly et al. 1996), and an overlying volcanic unit at its top dated at 3226 Ma ± 1 Myr (Kamo and Davis 1994). Hence, the minimum date we would assign is 3225 Myr.
**Hard maximum age justification:** The Moon forming impact would have effectively sterilized the Earth and so it serves as an effective basis for establishing a hard maximum age constraint. Pb-Pb dating carried out on Moon rocks, yielding a date of 4.51 Ga ± 10 Myr (Hanan and Tilton 1987) a date which has also recently been confirmed by reanalysis of the Apollo mission zircons (Barboni et al. 2017). Thus, our maximum constraint is 4.52 Ga.

**Crown Cyanobacteria | 2013.6-3448 Ma**
**Clade:** This equates to the extant clade of Cyanobacteriia, excluding Vampiriovibrionia and Sericytochromatia (Shih et al. 2017).
**Fossil taxon and specimen**: *Eoentophysalis belcherensis*. (Holotype) GSC type no. 42770, from the upper part of Kasegalik Formation, Belcher Supergroup, unnamed island at north end of Churchill Sound (Locality A of Hofmann, 1976).
**Phylogenetic justification**: Compared to extant *Entophysalis* (Chroococcales) by (Demoulin et al. 2019) based on its similarly sized coccoidal cells, colonial organization resulting from binary fission in three perpendicular planes, and warty (pustular) mamilate-shaped outer layers (Demoulin et al. 2019). These characteristics are convincing of cyanobacterial affinity, however, on the basis of such necessarily limited evidence, that *Eoentophysalis belcherensis* is a close relative of extant *Entophysalis*, not least since so little of traditional cyanobacterial taxonomy (based in cytological, cell division patterns and arrangements) has been corroborated by molecular phylogenetics. Indeed, we cannot be certain that *Eoentophysalis belcherensis* is a member of crown-Cyanobacteriia, however, evidence of its phenotype and development is sufficient to conclude with high probability that it is a member of crown-Cyanobacteriia and we therefore use it as a basis for establishing a soft minimum constraint. The alternative would be to rely on *Bangiomorpha pubescens* but this inference of a eukaryote plastid is, by its very nature, remote from the timing of origin of crown-Oxyphotobacteria.
**Soft minimum age justification:** The age of the Kasegalik Formation (Belcher Supergroup) has been established on the basis of U-Pb dating of tufts near the base and top (Hodgskiss et al. 2019). The latter is dated to 2015.4 Ma ± 1.8 Myr, providing for a 2013.6 Ma minimum constraint on the oldest record of *Eoentophysalis belcherensis* and, therefore, crown-Cyanobacteria.
**Soft maximum age justification:** Claims of Oxyphotobacteria in the Strelley Pool Formation can be rationalised in other ways (see total-group Bacteria, above). Will implement a soft maximum to allow for the possibility that Oxyphotobacteria were established prior to the deposition of the Strelley Pool Formation. The maximum depositional age of the Strelley Pool Formation has been dated to 3414 ± 34 Ma (Gardiner et al. 2019), yielding a maximum age interpretation of 3448 Ma.

**Total-group of Nostocales and Stigonematales: Heterocystous/akinete-forming Cyanobacteria (Cyanobacteria sections IV+V):**
***Fossil taxon and specimen:*** *Anhuithrix magna* [LIb x4B, Institute of Geology and Paleontology, Technical University of Berlin,Berlin, Germany] from the Liulaobei Formation in the Huainan region of North China
**Phylogenetic justification:** *A. magna* was originally described as a tubular organism (Steiner 1994), however Pang et al. (2018) interpreted it as a filamentous cyanobacteria based on new material. Based on the presence of binary fission, hormogonia, akinetes, and probably heterocysts, an affinity with subsections IV+V of cyanobacteria was proposed.
**Soft minimum age:** 996 Ma
**Hard maximum age:** 3261 Ma
**Soft Minimum age justification:** The age of the Liulaobei Formation has been constrained through SIMS Pb-Pb dating of phosphates to 1086 Ma ± 120 Myr, providing for a minimum of 966 Ma (Zhang et al. 2022).
**Soft maximum age justification:** The oldest credible claim of evidence of Cyanobacteriia is evidence of stable Fe and U-Th-Pb isotopes in the Manzimnyama Banded Ironstone Formation (Fig Tree Group, Barberton, South Africa), indicating a level of free oxygen indicative of cyanobacterial activity. Necessarily, this long precedes the total-group of Nostocales and Stigonematales. The age of the Fig Tree Group is well constrained with a spherule layer at its base dated at 3258 Ma ± 3 Myr (Byerly et al. 1996), yielding a maximum constraint of 3261 Ma. We implement this as a hard constraint since, a priori, we consider it remote from the timing of origin of the total-group of Nostocales and Stigonematales.
**Discussion**: (Schirrmeister, Sanchez-Baracaldo, and Wacey 2016) and (Demoulin et al. 2019) consider the many claims of fossil heterocystous cyanobacteria, some extending into the Palaeoproterozoic (e.g. (Sergeev, Sharma, and Shukla 2012)); *Anhuithrix magna* is the most convincing though Demoulin et al. (2019) question whether the differences in cell phenotype are taphonomic in nature. To allow for this possibility, we implement this as a soft minimum.

**Crown-Archaeplastida / Crown plastids | 1030-1879.6 Ma**
**Specimen and fossil taxon:** *Bangiomorpha pubescens*. (HUPC 62912, Slide HUST-1A, England Finder coordinates: O-35) from the Lower Hunting Formation, Somerset Island, Arctic Canada.
**Soft minimum age**: 1030 Ma
**Soft maximum age:** 1879.6 Ma
**Minimum age justification:** Re-Os isotopic dating of sedimentary rocks in the stratigraphic region in which *Bangiomorpha pubescens* was sampled date this fossil at 1.047 +0.013/–0.017 Ga, yielding a minimum age constraint of 1030 Ma (Gibson et al. 2018).
**Hard maximum age justification**: is based on the maximum age interpretation of the Gunflint Chert, a diverse and well-documented microbiota that is widely interpreted to be absent of eukaryotes. This has been dated to 1878.3 Ma ± 1.3 Myr (Fralick, Davis, and Kissin 2002), yielding a maximum age of 1879.6 Ma.

**Crown-Mitochondria | 1030-1879.6 Ma**
**Specimen and fossil taxon:** *Bangiomorpha pubescens*. (HUPC 62912, Slide HUST-1A, England Finder coordinates: O-35) from the Lower Hunting Formation, Somerset Island, Arctic Canada.

**Soft minimum age:** 1030 Ma
**Soft maximum age:** 1879.6 Ma
**Minimum age justification:** Re-Os isotopic dating of sedimentary rocks in the stratigraphic region in which *Bangiomorpha pubescens* was sampled date this fossil at 1.047 +0.013/–0.017 Ga, yielding a minimum age constraint of 1030 Ma (Gibson et al. 2018).
**Hard maximum age justification**: is based on the maximum age interpretation of the Gunflint Chert, a diverse and well-documented microbiota that is widely interpreted to be absent of eukaryotes. This has been dated to 1878.3 Ma ± 1.3 Myr (Fralick, Davis, and Kissin 2002), yielding a maximum age of 1879.6 Ma.

**Mitochondria total group | 1030-4520 Ma**
**Specimen and fossil taxon:** *Bangiomorpha pubescens*. (HUPC 62912, Slide HUST-1A, England Finder coordinates: O-35) from the Lower Hunting Formation, Somerset Island, Arctic Canada.
**Soft Minimum age**: 1030 Ma
**Hard maximum age:** 4520 Ma
**Minimum age justification:** Re-Os isotopic dating of sedimentary rocks in the stratigraphic region in which *Bangiomorpha pubescens* was sampled date this fossil at 1.047 +0.013/–0.017 Ga, yielding a minimum age constraint of 1030 Ma (Gibson et al. 2018).
**Hard maximum age justification**: The Moon forming impact would have effectively sterilized the Earth and so it serves as an effective basis for establishing a hard maximum age constraint. Pb-Pb dating carried out on Moon rocks, yielding a date of 4.51 Ga ± 10 Myr (Hanan and Tilton 1987) a date which has also recently been confirmed by reanalysis of the Apollo mission zircons (Barboni et al. 2017). Thus, our maximum constraint is 4.52 Ga.

**Plastid total group | 1030-4520 Ma**
**Specimen and fossil taxon:** *Bangiomorpha pubescens*. (HUPC 62912, Slide HUST-1A, England Finder coordinates: O-35) from the Lower Hunting Formation, Somerset Island, Arctic Canada.
**Soft Minimum age**: 1030 Ma
**Hard maximum age:** 4520 Ma
**Minimum age justification:** Re-Os isotopic dating of sedimentary rocks in the stratigraphic region in which *Bangiomorpha pubescens* was sampled date this fossil at 1.047 +0.013/–0.017 Ga, yielding a minimum age constraint of 1030 Ma (Gibson et al. 2018).
**Hard maximum age justification**: The Moon forming impact would have effectively sterilized the Earth and so it serves as an effective basis for establishing a hard maximum age constraint. Pb-Pb dating carried out on Moon rocks, yielding a date of 4.51 Ga ± 10 Myr (Hanan and Tilton 1987) a date which has also recently been confirmed by reanalysis of the Apollo mission zircons (Barboni et al. 2017). Thus, our maximum constraint is 4.52 Ga.

**Total-group Chromatiaceae | 1631-4520 Ma**
**Fossil taxon and specimen**: Okenane biomarker record from the Barney Creek Formation, McArthur Basin, Australia (Brocks and Schaeffer 2008).
**Phylogenetic justification**: Okenane is described as specific to Chromatiaceae (Brocks and Schaeffer, 2008). (Fournier et al. 2021) contend that it may have a more general distribution among Chlorobiaceae but this view is based solely on incompatibility among the calibrations used in that study.
**Soft Minimum age**: 1631 Ma

**Hard maximum age:** 4520 Ma
**Soft minimum age justification:** The Barney Creek Formation has recently been dated to 1642.2 Ma ± 3.9 Myr, but this is from a sample close to the base of the Formation (Munson et al. 2020). Page and Sweet (1998) provide dates from tufts higher in the sequence, the youngest of which is 1638 Ma ± 7 Myr, providing for a minimum constraint of 1631 Ma (Page and Sweet 1998).
**Hard maximum age justification:** The Moon forming impact would have effectively sterilized the Earth and so it serves as an effective basis for establishing a hard maximum age constraint. Pb-Pb dating carried out on Moon rocks, yielding a date of 4.51 Ga ± 10 Myr (Hanan and Tilton 1987) a date which has also recently been confirmed by reanalysis of the Apollo mission zircons (Barboni et al. 2017). Thus, our maximum constraint is 4.52 Ga.

**Crown Chlamydiia | 1030-4520 Ma**
**Specimen and fossil taxon:** *Bangiomorpha pubescens*. (HUPC 62912, Slide HUST-1A, England Finder coordinates: O-35) from the Lower Hunting Formation, Somerset Island, Arctic Canada.
**Phylogenetic justification:** Recent work (Stairs et al. 2020) suggests that members of the Anoxychlamydiales (including a lineage named Chlam. Bact SM23_39), a clade within Chlamydiia (and Chlamydiae), donated at least three subunits of hydrogenase maturase (E, F, G) to stem eukaryotes prior to the radiation of crown eukaryotes. These phylogenies imply that a minimum age for Anoxychlamdiales is the age of the oldest crown eukaryote.
**Soft minimum age**: 1030 Ma
**Hard maximum age**: 4520 Ma
**Minimum age justification:** Re-Os isotopic dating of sedimentary rocks in the stratigraphic region in which *Bangiomorpha pubescens* was sampled date this fossil at 1.047 +0.013/–0.017 Ga, yielding a minimum age constraint of 1030 Ma (Gibson et al. 2018).
**Hard maximum age justification:** The Moon forming impact would have effectively sterilized the Earth and so it serves as an effective basis for establishing a hard maximum age constraint on LUCA. Pb-Pb dating carried out on Moon rocks, yielding a date of 4.51 Ga ± 10 Myr (Hanan and Tilton 1987) a date which has also recently been confirmed by reanalysis of the Apollo mission zircons (Barboni et al. 2017). Thus, our maximum constraint is 4.52 Ga.

**Total Group Eukaryota | 1619.1-4520 Ma**
**Fossil taxon and specimen:** Changzhougou Formation, North China, following the justification outlined in Betts et al. (2018).
**Soft minimum age**: 1619.1 Ma
**Hard maximum age**: 4520 Ma
**Minimum age justification:** The minimum age of the Changzhougou Formation is established based on dated ashes in the overlying Chuanlinggou Formation, dated to 1625.3 ± 6.2 Myr (Li et al. 2013), thus 1619.1 Ma.
**Hard maximum age justification:** The Moon forming impact would have effectively sterilized the Earth and so it serves as an effective basis for establishing a hard maximum age constraint on LUCA. Pb-Pb dating carried out on Moon rocks, yielding a date of 4.51 Ga ± 10 Myr (Hanan and Tilton 1987) a date which has also recently been confirmed by reanalysis of the Apollo mission zircons (Barboni et al. 2017). Thus, our maximum constraint is 4.52 Ga.

**Crown Group Eukaryota | 1030-1879.6 Ma**

**Fossil taxon and specimen:** *Bangiomorpha pubescens* (HUPC 62912; Slide HUST-1A, England Finder coordinates: O-35; Paleobotanical Collections of Harvard University, USA) from the Hunting Formation in northwestern Somerset Island, Arctic Canada (Butterfield 2000).

**Phylogenetic justification:** Morphological similarity of *Bangiomorpha pubescens* with modern red algae *Bangia (Butterfield 2000)* evidences affinity to total group Rhodophyta; this is the oldest unequivocal record of a crown-eukaryote.

**Hard minimum age**: 1030 Ma

**Soft maximum age:** 1879.6 Ma

**Minimum age justification:** Re-Os isotopic dating of sedimentary rocks in the stratigraphic region in which *Bangiomorpha pubescens* was sampled date this fossil at 1.047 +0.013/–0.017 Ga, yielding a minimum age constraint of 1030 Ma (Gibson et al. 2018).

**Hard maximum age justification:** The Gunflint Chert microflora has a long history of study, including claims of eukaryotes. These include process-bearing acritarch-like cysts, such as *Germinosphaera*, suggesting the presence of an actin cytoskeleton. Nevertheless, all such claims of eukaryote affinity have generally been rejected (Agić 2021). The Gunflint Chert has been dated to 1878.3 Ma ± 1.3 Myr (Fralick, Davis, and Kissin 2002), yielding a maximum age of 1879.6 Ma.

**Crown Protostomia | *Lottia-Limulus* | 532-590.8 Ma**

**Fossil taxon and specimen:** *Aldanella yanjiahensis*, from the Dahai member of the Zhujiaqing Formation in the middle Meishucunian of China, TU Berlin collection NO. YXII02-02 (Steiner et al. 2007).

**Phylogenetic justification:** *Aldanella janjiahensis* (junior synonym *Aldanella attleborensis*) is a dextrally-coiled stem group gastropod assigned to Pelagiellida (Runnegar 1981).

**Hard minimum age:** 532 Ma

**Soft maximum age:** 590.8 Ma

**Minimum age justification:** *Aldanella yanjiahensis* is associated with *Watsonella crosbyi* and *Oelandiella korobkovi* in the Dahai member in the middle Meishucunian of China (Steiner et al. 2007). Chemostratigraphic correlation places this unit in the Nemakit Daldynian within the interval 534-532 Ma (Maloof et al. 2010).

**Soft maximum justification:** Established based on the Weng'an biota which may contain total group metazoans (Yin, Sun, Liu, et al. 2022; Yin, Sun, Reitner, et al. 2022), but there is no convincing evidence of crown metazoans. (Yang et al. 2021) establish a 590.8 Ma maximum age for the Weng’an Biota.

**Embryophyta: Bryophyta – Tracheophyta | 430.54-515.5 Ma**

**Fossil taxon and specimen:** *Cooksonia barrandei* (D 552a,b, National Museum, Prague) from the Motol Formation at Loděnice, Špičatý vrch-Barrandovy Jámy, Czech Republic (Libertín et al. 2018).

**Phylogenetic justification**: Dichotomously branching axes with terminal sporangia containing trilete spores, all of which are total-group tracheophyte characters at the least (Clarke, Warnock, and Donoghue 2011).

**Hard minimum age:** 430.54 Ma

**Soft maximum age:** 515.5 Ma

**Minimum age justification:** the holotype was recovered from *Monograptus belophorus* Biozone, Motol Formation, Middle Sheinwoodian, Wenlock, Silurian. The top of the *M.*

*belophorus* Biozone is dated to 430.54 Ma (Melchin, Sadler, and Cramer 2020).
**Maximum age justification:** A soft maximum age constraint for crown group Embryophyta must encompass all total group Embryophyta, including all cryptospores (sensu stricto (Paul K. Strother 2016)). The oldest deposits to yield probable non-marine palynomorphs include the Rogersville Shale and Pumpkin Valley Shale of the Conasauga Group of east Tennessee (P. K. Strother and Beck 2000), and the Bright Angel Shale of the Tonto Group of Arizona (Baldwin et al. 2004), correlated to the Glossopleura and Ehmaniella Laurentian trilobite Biozones, dated to Stage 5 of Series 3, the oldest part of the Mid Cambrian (Paul K. Strother 2016). Cryptospores have recently been reported from the Rome Formation (Paul K. Strother 2016) that underlies the Conasauga Group and is correlated to the Olenellus trilobite Biozone of Stage 4 of Series 2, the latest part of the Early Cambrian. The base of the Olenellus Biozone is dated to approximately 515.5 Ma (Peng, Babcock, and Ahlberg 2020).

**Mollusca-Annelida | 532-590.8 Ma**

Our minimum constraint on crown-Bilateria follows the justification for crown-Mollusca in (Benton et al. 2015).

**Hard minimum age:** 532 Ma

**Soft maximum age:** 590.8 Ma

**Minimum justification:** *Aldanella janjiahensis*, or *Aldanella attleborensis* is a widely accepted stem group gastropod (Runnegar 1981), the oldest record of which is dated to 532 Ma (Steiner et al. 2007).

**Soft maximum justification:** Weng'an biota (Yang et al. 2021) which may contain total group metazoans (Yin, Sun, Reitner, et al. 2022; Yin, Sun, Liu, et al. 2022), but there is no convincing evidence of crown metazoans.

**Mandibulata: Insect-Chelicerate | 514-543 Ma**

Our minimum constraint on crown-Mandibulata follows the justification in (Benton et al. 2015) and (Wolfe et al. 2016).

**Hard minimum age:** 514 Ma

**Soft maximum age:** 543 Ma

**Minimum justification:** *Yicaris dianensis* and *Wijicaris muelleri* both have limb characteristics that place them within crown Crustacea (Benton et al. 2015; Wolfe et al. 2016).

**Soft maximum justification:** *Rusophycus* trace fossils are widely accepted to be made by arthropod-grade organisms with bilateral symmetry and evidence of segmented limbs (age from (Narbonne, Myrow, and Landing 1987; Benton et al. 2015; Wolfe et al. 2016). They evidence the existence of arthropods, but there is no evidence of mandibulates at this time.

**crown-Opisthokonta | 879-1879.6 Ma**

**Fossil taxon and specimen:** *Ourasphaira giraldae* (Specimen 74639-W46,3, sample 15RAT-021A1 in the collections of the Department of Geology in the University of Liège, Belgium) from the Shale of the Grassy Bay Formation in the Brock Inlier, Northwestern Territories, Canada (Loron, Rainbird, et al. 2019).

**Phylogenetic justification:** (Loron, François, et al. 2019) establish a eukaryote affinity on the basis of a ‘combination of complex morphology, right-angle branching, multicellularity, bilayered wall ultrastructure, compositional recalcitrance and relatively large size’,

evidencing the presence of a complex cytoskeleton; combined with their FTIR spectroscopy which diagnoses the presence of chitin, they conclude a total-group Dikarya affinity for *Ourasphaira giraldae*.
**Hard minimum age:** 879 Ma
**Soft maximum age:** 1879.6 Ma
**Minimum age justification:** There are no direct dates for the Grassy Bay Formation but the overlying Boot Inlet Formation has been dated to 892 Ma ± 13 Myr (van Acken et al. 2013). This provides for a minimum age constraint of 879 Ma.
**Hard maximum age justification:** The Gunflint Chert microflora has a long history of study, including claims of eukaryotes. These include process-bearing acritarch-like cysts, such as *Germinosphaera*, suggesting the presence of an actin cytoskeleton. Nevertheless, all such claims of eukaryote affinity have generally been rejected (Agić 2021). The Gunflint Chert has been dated to 1878.3 Ma ± 1.3 Myr (Fralick, Davis, and Kissin 2002), yielding a maximum age of 1879.6 Ma.

**Viridiplantae/Chloroplastida: Chlorophyta - Streptophyta | 940.4-1879.6 Ma**
Our minimum and maximum constraints on crown-Chloroplastida follows the justification for in (Harris et al. 2022).

**Great Oxidation Event**: (Poulton et al. 2021) associate the inception of permanent oxygenation of the atmosphere with the Lomagundi carbon isotope excursion. (Martin et al. 2013) put a maximum age constraint on this as 2306 Ma ± 9 Myr. This provides for a maximum constraint of 2315 Ma.

### 5. Molecular dating analyses

We performed the molecular dating analyses using McmcDate (https://github.com/dschrempf/mcmc-date) , which has the ability to use both relative age calibriations (Harris et al. 2022; Szöllõsi et al. 2022) and node braces ((Harris et al. 2022; Mahendrarajah et al. 2023)). First, we sampled species tree branch lengths using the LG+G4 model in PhyloBayes on the fixed unrooted species tree inferred above (section 1, this document). Two chains were run until at least 19,000 iterations until. Due to computational reasons the resulting treelist of the first chain was thinned by retaining every second tree in downstream analysis. Then, we inferred dated phylogenies based on (i) the concatenation alone with a single calibration at the root corresponding to the Moon-forming impact ("Root Maximum"), (ii) the concatenation, calibrated with the fossils and geochemical calibrations summarised in section 4 above ("Fossils"), then (iii) the concatenation, fossils and the GOE soft maximum at 2.315Ga for nodes annotated as aerobic using the XGBoost classifier. (iv) the concatenation, fossils and the GOE soft maximum at 2.315Ga for nodes annotated as aerobic using the ExtraTree classifier. (v) the concatenation, fossils and the GOE soft maximum at 2.315 Ga for nodes annotated as aerobic using the Concatenate-based approach.

The analyses used McmcDate with the calibrations described above, an uncorrelated gamma relaxed molecular clock and setting the sparse multivariate normal parameter to 0.1. An example command line parametrization of McmcDate is the following:

```
mcmc-date run --preparation-name cyan28 --analysis-name XGBoostSoft --calibrations "csv ./calibrations.csv" --ignore-problematic-calibrations --braces "./braces" --relaxed-molecular-clock "UncorrelatedGamma" --likelihood-spec "SparseMultivariateNormal 0.1" +RTS -M14G -N2
```

For each analysis the burnin was 4930 iterations using a custom auto-tuning scheme which tunes the model parameters after a specific number of iterations. (E.g. for the calibration (v) after iterations: `[[10, 10, 10, 20, 30, 40, 50, 60, 70, 80, 90, 100, 110, 120, 130], [100, 120, 140, 160, 180, 200, 220, 240, 260, 280, 300, 320, 340, 360, 380, 400]]`.) After the burnin 8000 further iterations were made, the results presented here are based on the statistics of these.

After the MCMC dating process, we summarised the posterior distribution to generate timetrees and ratetrees. All the results from these analyses can be found in the Online Data Supplement in the folder Chronograms.

#### Correlation with RED values

We correlated RED values (obtained by using a fixed rooted position) and the different consensus chronograms from the different molecular clock analyses. We computed the Spearman rho values for all correlations. In all cases, we see that the number is well above 0.93. These results suggest that the RED values may be an adequate proxy to be used for taxonomic studies.

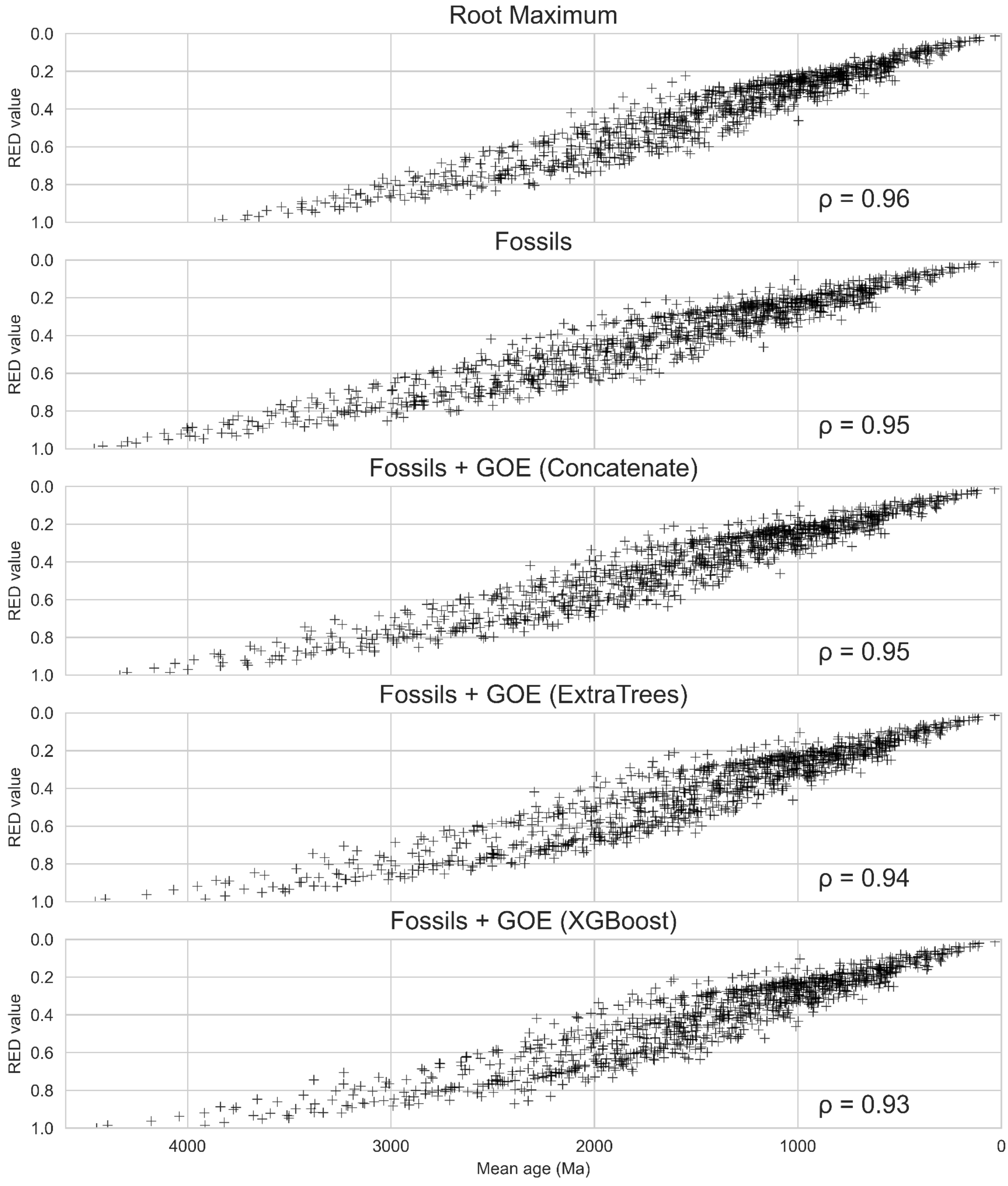


***Figure S6: Correlation between the mean age of ancestral nodes inferred using the relaxed molecular clock and RED values.*** *Under all analysis conditions, the correlation between RED values and ages estimated using the molecular clock are highly correlated, suggesting both methods capture a congruent signal as to the age of ancestral nodes.*

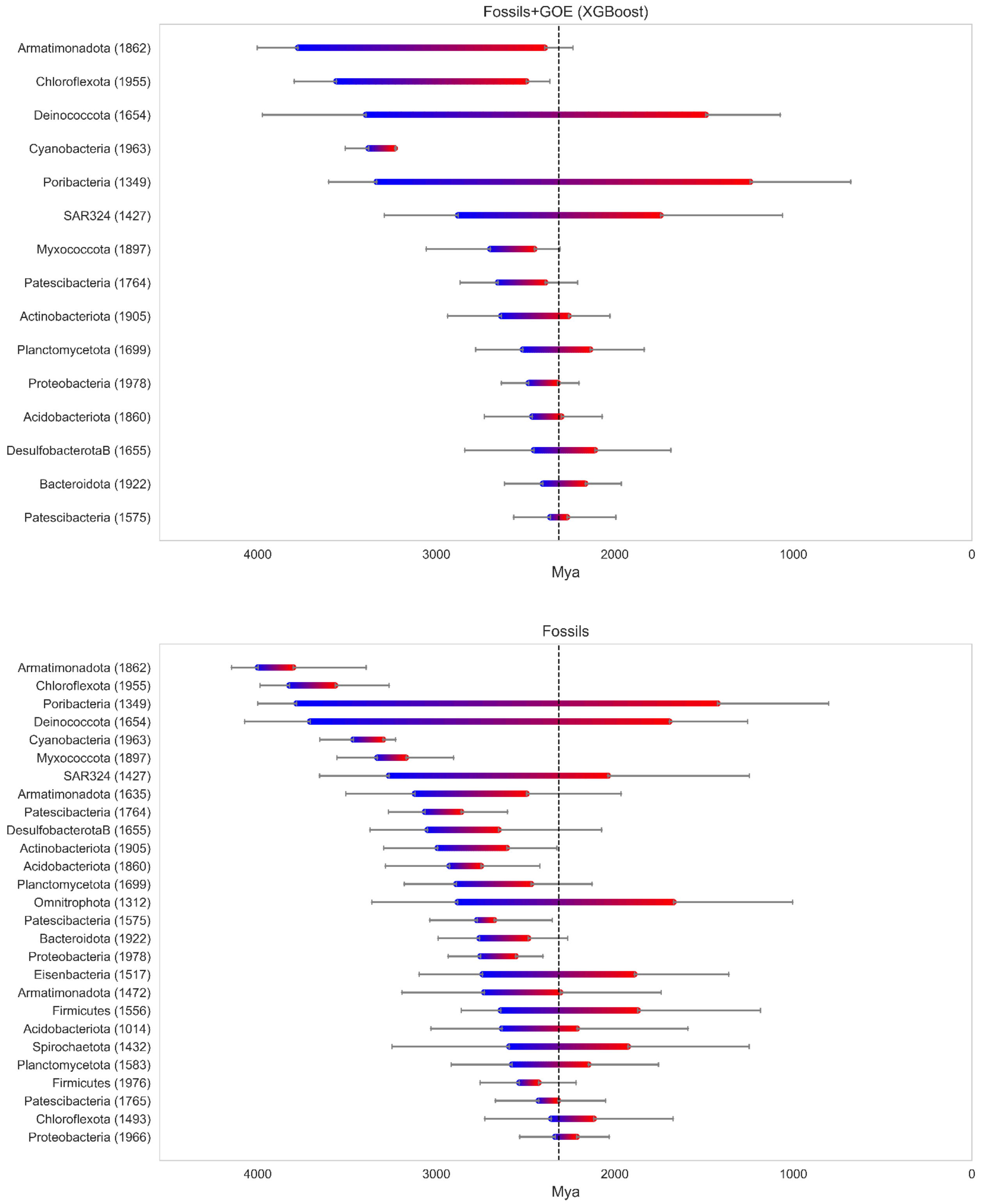


***Figure S7: Oldest transitions to aerobicity.*** *In these plots, we represent the oldest branches transitioning to aerobicity, selecting those whose starting node is older than the GOE and that the middle point of the branch is at least 2 Ga. The numbers between parentheses correspond to the name of the node. The first name is the most common phyla that diversified below that branch. The top panel plots the oldest branches for the dating analysis using Fossils+GOE (XGBoost). The lower panel plots the oldest branches for the dating analysis using only fossils.*

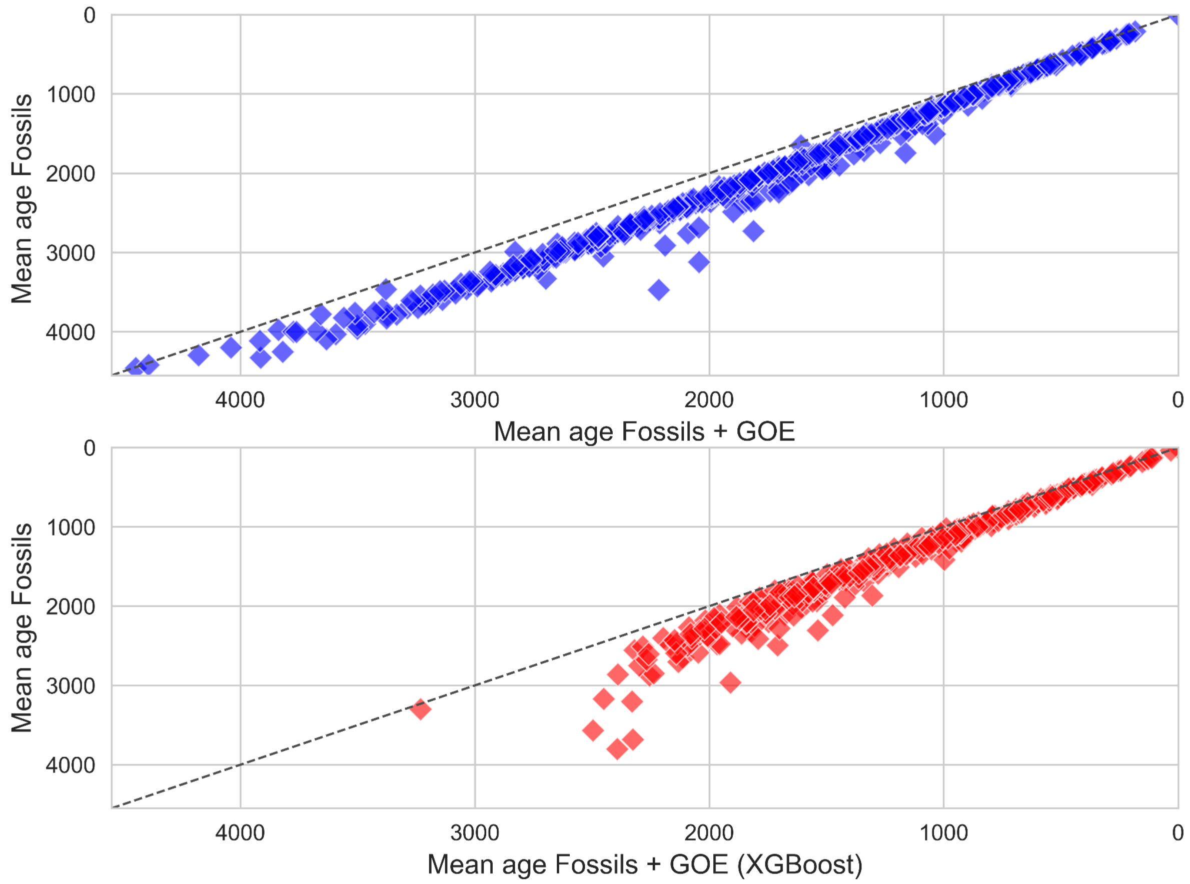


***Figure S8. Impact on node ages of the GOE constraint.*** *As expected, the maximum GOE constraint effectively pushes most nodes to be younger. Anaerobic nodes across the tree are (on average) 247 my younger in the Fossils+GOE analysis, while aerobic nodes are 195 my younger.*

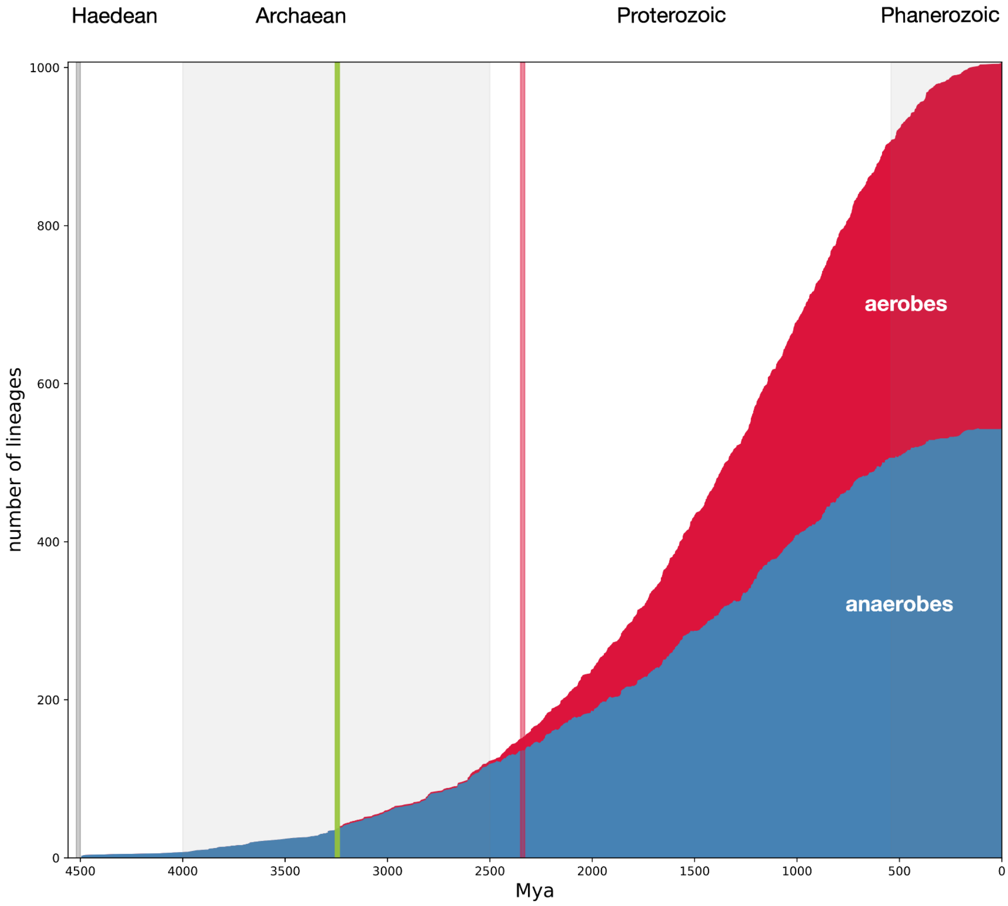


***Figure S9: Lineages through time.*** *The number of aerobic and anaerobic lineages through time. The grey line shows the Moon-forming impact at ~4.52 Ga; the green line at ~3.23 Ga reflects the presence of fossil and isotopic evidence for oxygenic photosynthesis; the red line at ~2.33 Ga reflects the end of the GOE (see calibrations above).*

### 6. Analyses with time-constrained reconciliations

In order to evaluate the impact of distinct sources of temporal information - fossils and the GOE constraint - on the accuracy of our inferred time trees, we used time-constrained phylogenetic reconciliations. We implemented the  time-constrained gene tree-specioes tree model, an extension of the undated model in ALE (https://github.com/ssolo/ALE) which describes how a gene tree evolves under a rooted species tree and the probabilities of duplication, loss, and horizontal gene transfer events. This model is an extension of the undated DTL model, described in (Morel et al. 2020). However, unlike the undated DTL, the time-constrained DTL model takes into account the relative chronological order of speciation events to constrain the horizontal gene transfers, by prohibiting “transfers to the past” — that is, transfers from a species which originates at time t1 to a species that ends at time t2, with t1 being more recent than t2. Transfers “to the future” are allowed, because they could be the result of a transfer from an unsampled or extinct species (Szöllosi et al. 2013).

We refer to this new model subsequently as the time-constrained gene tree-species tree reconciliation model.

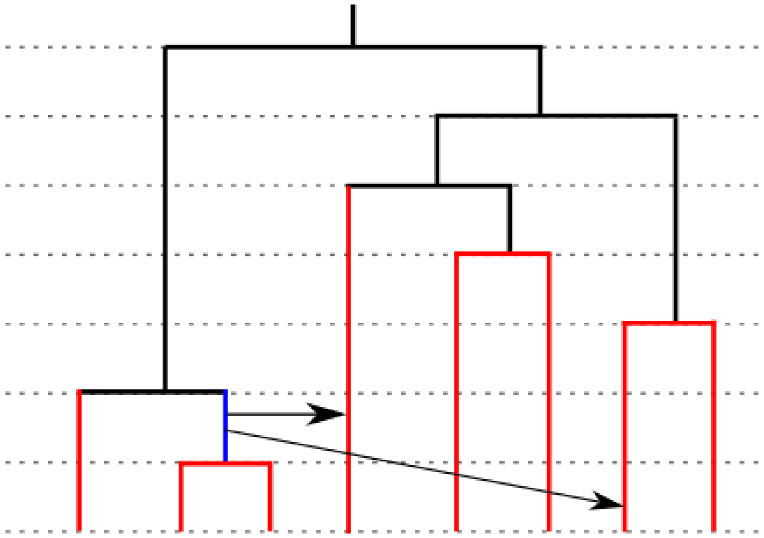

***Figure S10: Illustration of the constraints on gene transfers permitted under the time-constrained model.*** *The red branches are the species that are allowed to receive a transfer from the blue branch. The black branches are not allowed to receive a transfer from the blue branch because those transfers are impossible timewise. The arrows show examples of possible transfers.*

#### Using the time constrained model to compare dated species trees

We reconciled the gene families (described in section 3 above) with the dated species trees described in section 5, using the command:

```
ALEml_undated
 SpeciesTree ALEfile fraction_missing:fraction_missing.txt reldate
```

We inferred the reconciliations for 4731 for every of the dated species trees. The summed gene family log-likelihoods for the 5 candidate dated species trees are summarised in Table 1.

#### Simulations suggest that transfer events can identify the most accurate dated species tree

We conducted a simulation analysis to validate the ability of the time-constrained DTL model to differentiate among various dated trees. Our objective was to mimic real gene families within a known phylogeny and known dates. This setup allowed us to reconcile these gene families with different dated trees and ascertain if the reconciliations could correctly identify the original dated tree.

Initially, we derived the maximum likelihood parameters of several lognormal distributions that fit the distribution of duplications, transfers, and losses found in the real dataset, which can be found in the Online Data Supplement `MasterTableEvents.tsv`; see also Figure S11. The units are the number of events per gene per million years.

| Parameter | Mean | Standard deviation |
|---|---|---|
| Duplications | -11.382 | 1.296 |
| Transfers | -6.735 | 0.658 |
| Losses | -7.21 | 0.997 |

***Table S4. Most likely parameters for gene content evolution events (lognormal distributions).***

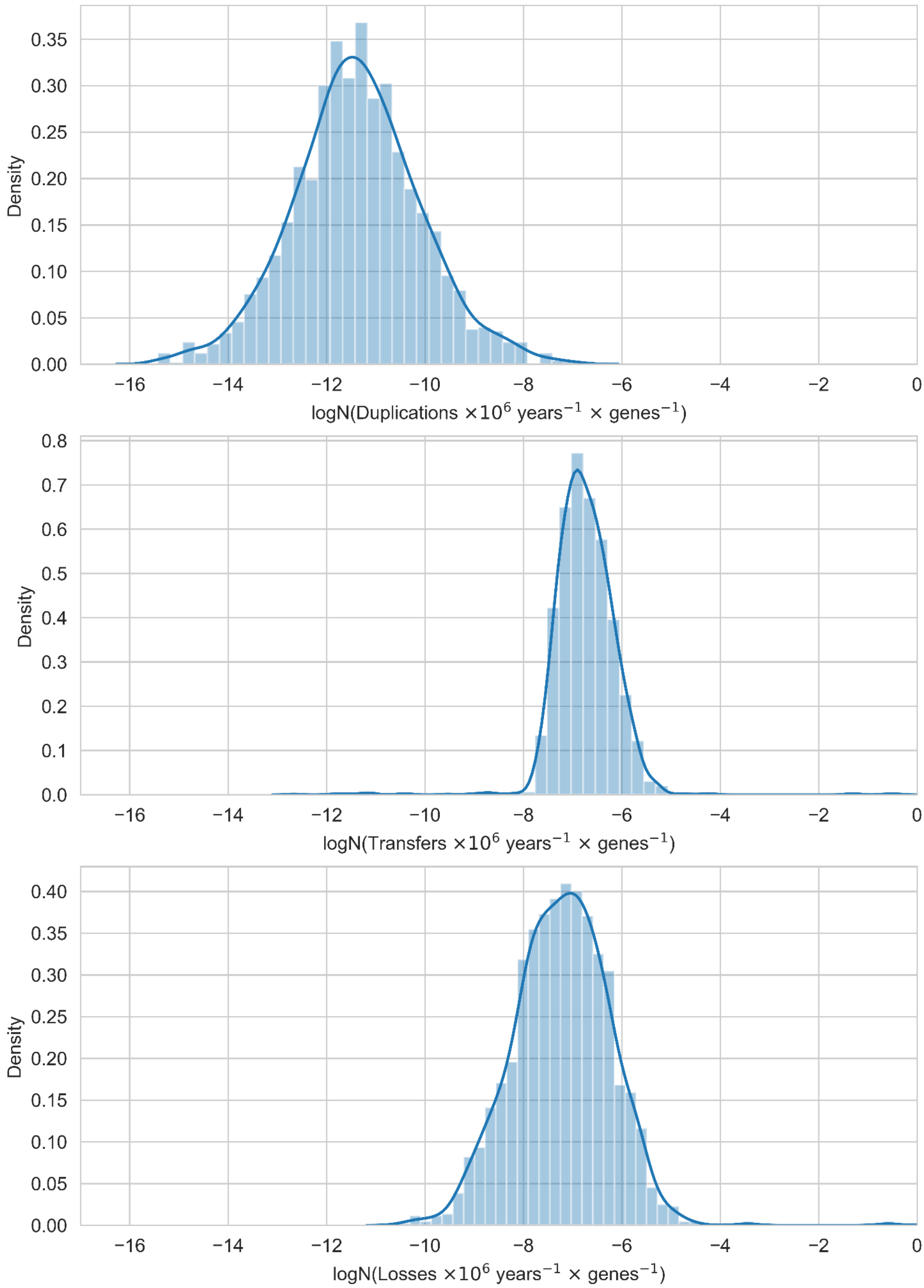


***Figure S11. Distribution of duplication, transfer and loss events in real gene families.*** *Inferred numbers of duplications, transfers and losses (per gene family and gene copy, per million years, note the log-scale) observed in the empirical gene families used in this study. To align our simulations as closely as possible to the real data, we fit lognormal distributions to these data and used the estimated parameters to set the duplication, transfer and loss rates in our simulations.*

We then simulated 100,000 families using Zombi (Davín et al. 2020) under the Gm model) and the primary tree shown in Figure 3 in the main text (Fossils + GOE (XGBoost)) as a species tree (using the Ti mode).

For each simulated family, we created a vector of length 1007, with each entry corresponding to the copy numbers of every genome for every gene family. We compared each of these simulated gene family with the largest 500 gene families in the real dataset, using the Euclidean distance and the corresponding copy number vector of the real families. We picked, for every real gene family the closest simulated family, resulting in 500 simulated families that resembled the real ones. We focused simulations on larger families because, as might be expected, these have previously been shown to contain most of the signal for distinguishing between species trees using reconciliations (Coleman et al. 2021).

Subsequently, we sampled 10 trees from the posterior distribution of the molecular clock analyses for each of the following experiments: Root Maximum (corresponding to the age of Earth), Fossils, and Fossils + GOE (XGBoost). The 30 sampled trees, along with the species tree used for gene tree simulation (the consensus mean tree from the Fossil+GOE (XGBoost) molecular clock analysis), were reconciled with the 500 simulated gene families. As a result of problems with numerical precision for large families in the current ALE implementation we were only able to calculate the likelihood of 409 of the 500 simulated gene trees for all dated trees. The resulting likelihoods from each reconciliation were summed to generate the plot in FigureSX. We compared the d2 distance (Kim, Rosenberg, and Palacios 2020) of every tree with the reference tree, that measures how different two trees are based on the order of speciations. There is a clear correlation (Figure S12) between both measures and the difference in LL inferred by the time-constrained DTL model with the reference tree. This indicates that the time-constrained DTL model can successfully distinguish between different dated trees.

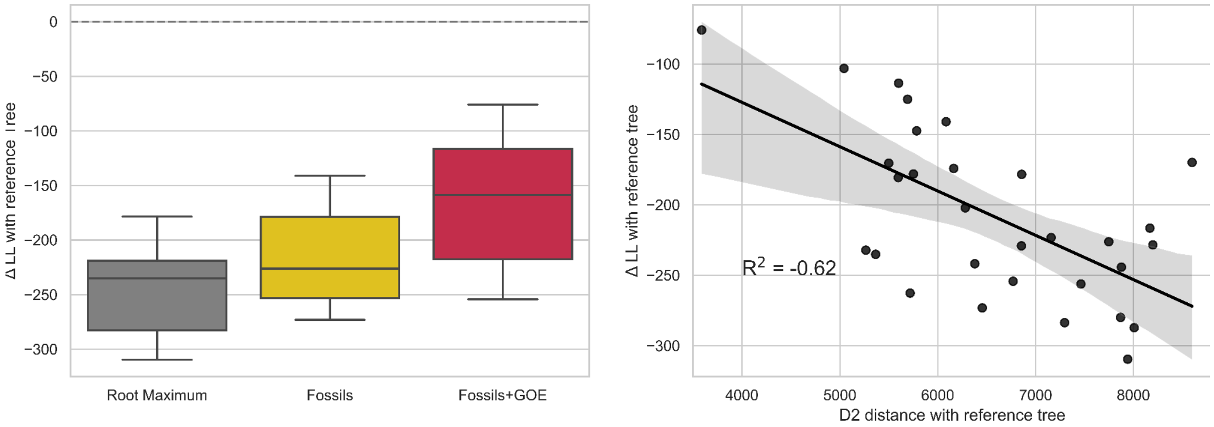


***Figure S12. Gene family log likelihoods under the time-constrained DTL model provide a measure of the accuracy of a dated species tree.*** *(a) We simulated gene families on the reference dated species tree and reconciled them against a series of sampled trees from the posterior distribution of trees in the analysis with real data. A total of 31 trees were used, 10 from each condition (Root Maximum, Fossils, Fossils+GOE(XGBoost)) plus the consensus tree of the posterior distribution of Fossils+GOE (XGBoost). The families were simulated on the consensus tree. The likelihood difference was computed for every tree compared to the reference tree. The results indicate that the time-constrained DTL model correctly identifies the reference tree and also that the likelihood increases when adding first Fossils and then the GOE constraint. (b) Correlation between d2 distance and LL. The more accurate the candidate dated species tree (that is, the closer the candidate dated species tree was to the simulation tree (d2 distance, x-axis)), the higher the summed gene family log-likelihood. This implies that gene family LLs can be used to determine the most accurate candidate-dated species tree. Based on this criterion, the dated tree inferred using both fossil calibrations and the GOE constraint is the most accurate in the empirical analysis.*

#### Robustness analyses

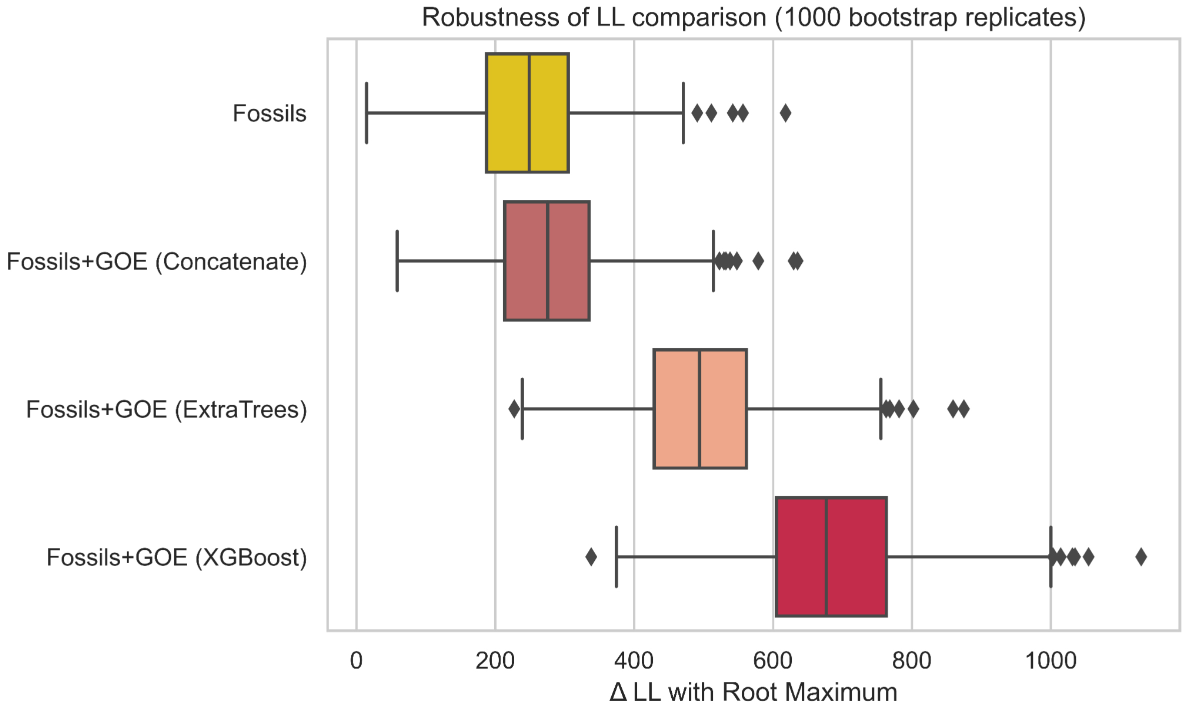


***Figure S13: Likelihood differences under the time-constrained DTL model robustly distinguish between dated trees.*** *We took the 4731 gene families used to distinguish between the different chronograms and performed a bootstrap analysis in which we resampled a given fraction of all the families and computed the difference in log-likelihood with the baseline Root Maximum tree calibrated only with the Moon-forming impact. The results indicate that the observed difference in gene family log-likelihoods between dated species trees is robust to resampling. This implies that the observed differences are not driven by differences in a small minority of families.*

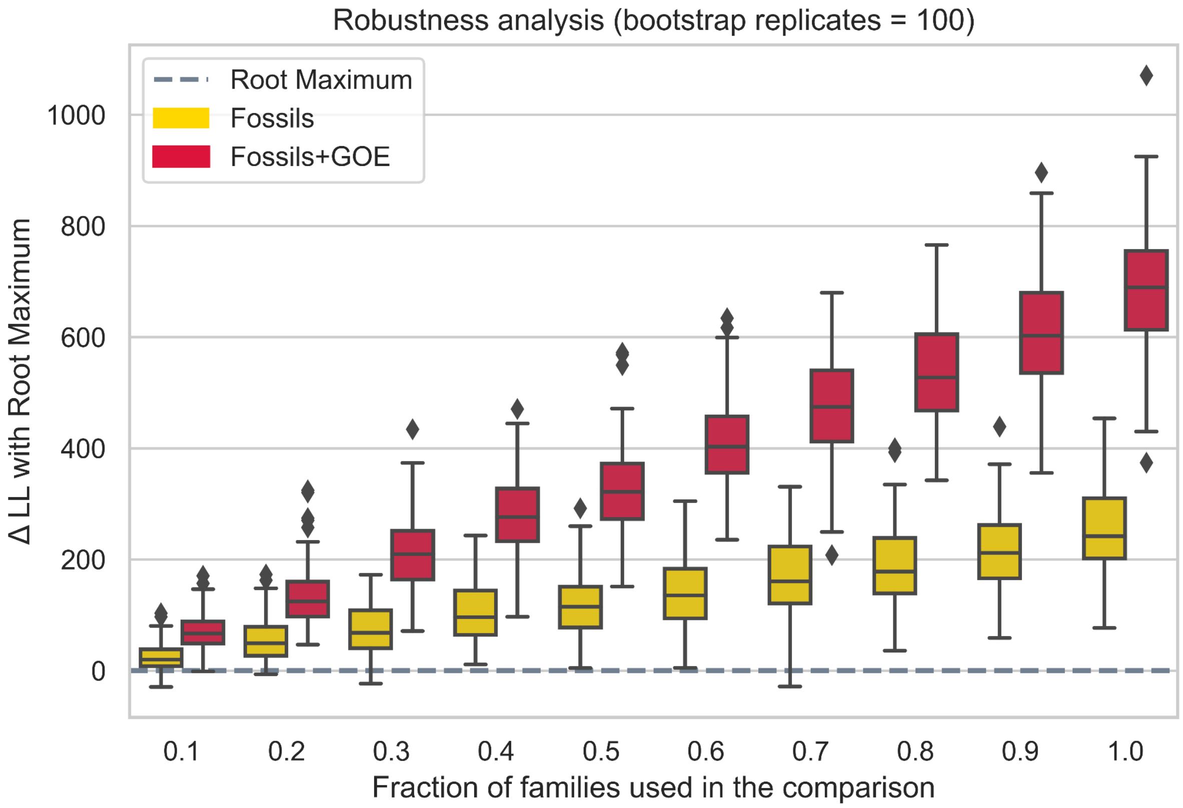


***Figure S14: A small proportion of gene families are already sufficient to distinguish between dated species trees using the time-constrained DTL model.*** *The difference in log likelihood is summarised between fossil calibrated and uncalibrated (only Root Maximum at the Moon-forming impact), and Fossil + GOE (XGBoost) calibrated and uncalibrated chronograms for n=100 bootstrap replicates over an increasing number of all the 4731 gene families used in this study. We can see that the preference of the different gene families for the chronogram dated using fossils and the GOE calibration still remains even when using a small fraction of families.*

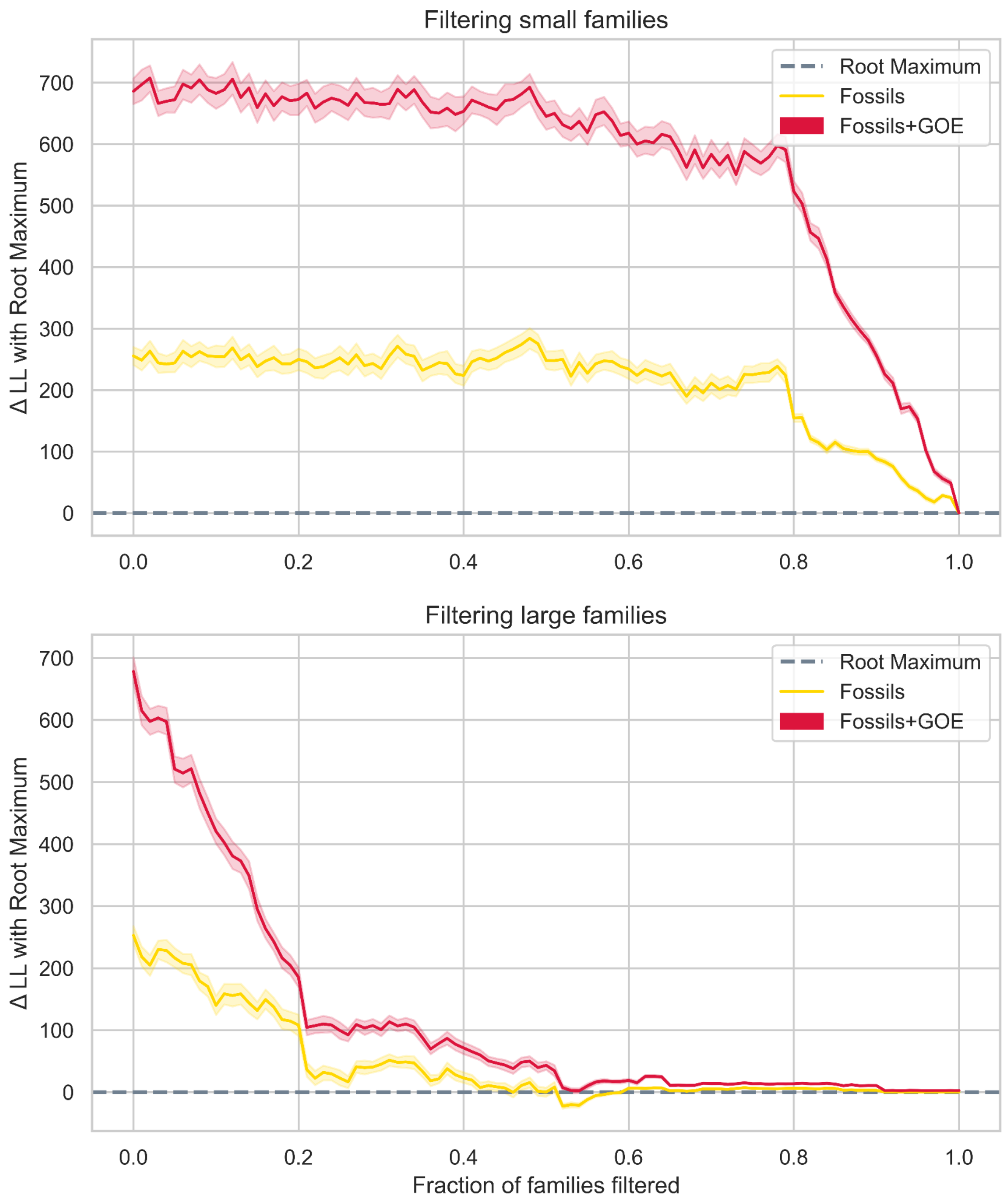


***Figure S15: Large gene families contain most of the signal for distinguishing between dated species trees using the time-constrained DTL model.*** *We filtered an increasing number of families, removing first the smallest families (on top) or first the largest families (bottom). Removing the smallest families has little effect on the LL difference. Most of the signal comes from the 20% largest families. Family size is defined as the number of different genes that share the same homologous cluster. In every step we do a resampling analysis (100 times) and obtain the total LL difference with the dated species tree calibrated only with the root maximum corresponding to the Moon-forming impact.*

#### Patterns of gene content evolution

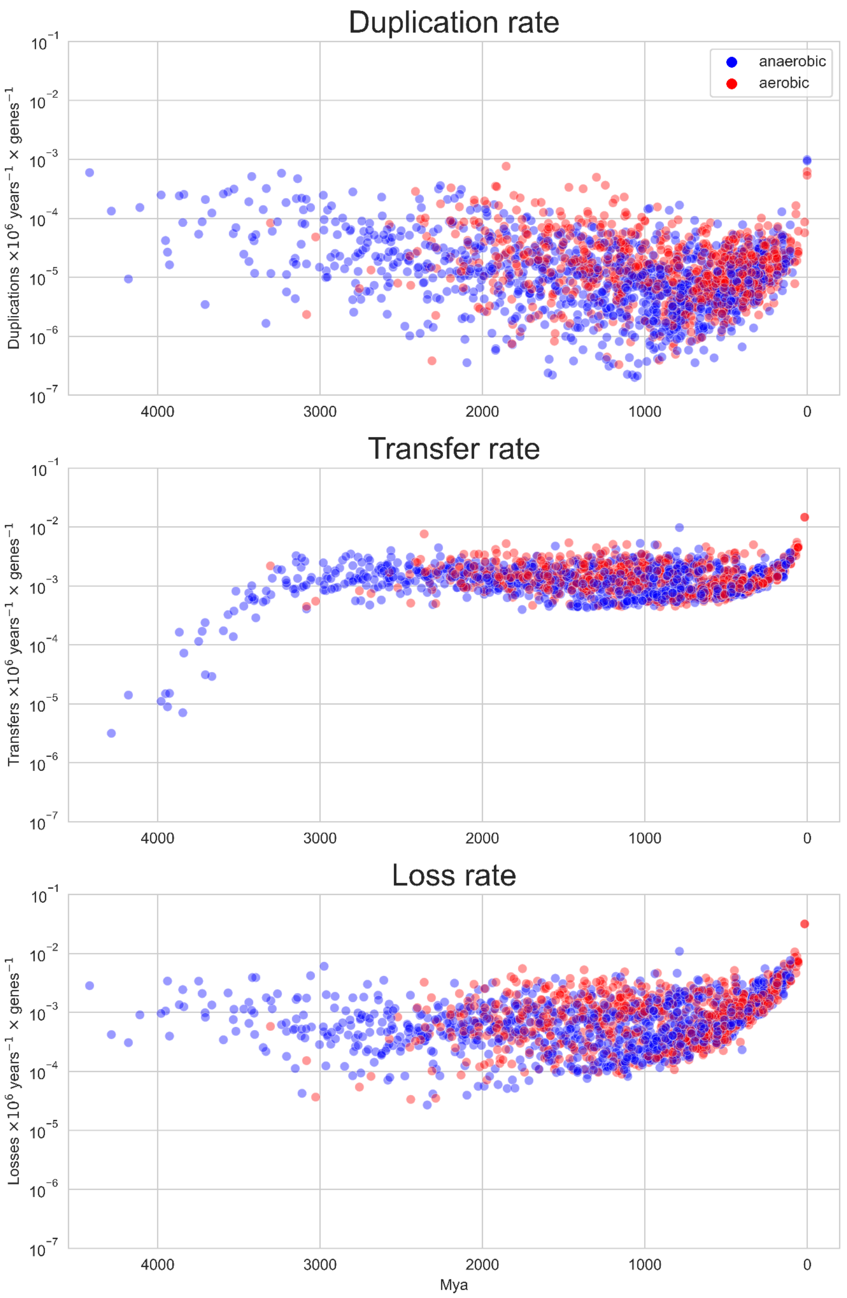


***Figure S16: Rates of gene duplication, transfer and loss in Bacteria estimated under the time-constrained DTL model.*** *Consistent with previous work (Treangen and Rocha 2011; Fernando D. K. Tria and Martin 2021; Williams et al. 2023), the rate of gene duplication (in events per gene per year) is much lower than the inferred transfer and loss rates - roughly two orders of magnitude in this analysis. The inferred transfer rate (b) is lower for early branches of the species tree, likely because of the small number of extant lineages that can be traced back to that time. This makes it more difficult to infer transfer events because only those HGTs that cross a sufficiently large phylogenetic distance to be seen as disagreeing with the species tree can be detected from the gene tree topology and therefore the reconciliation.*

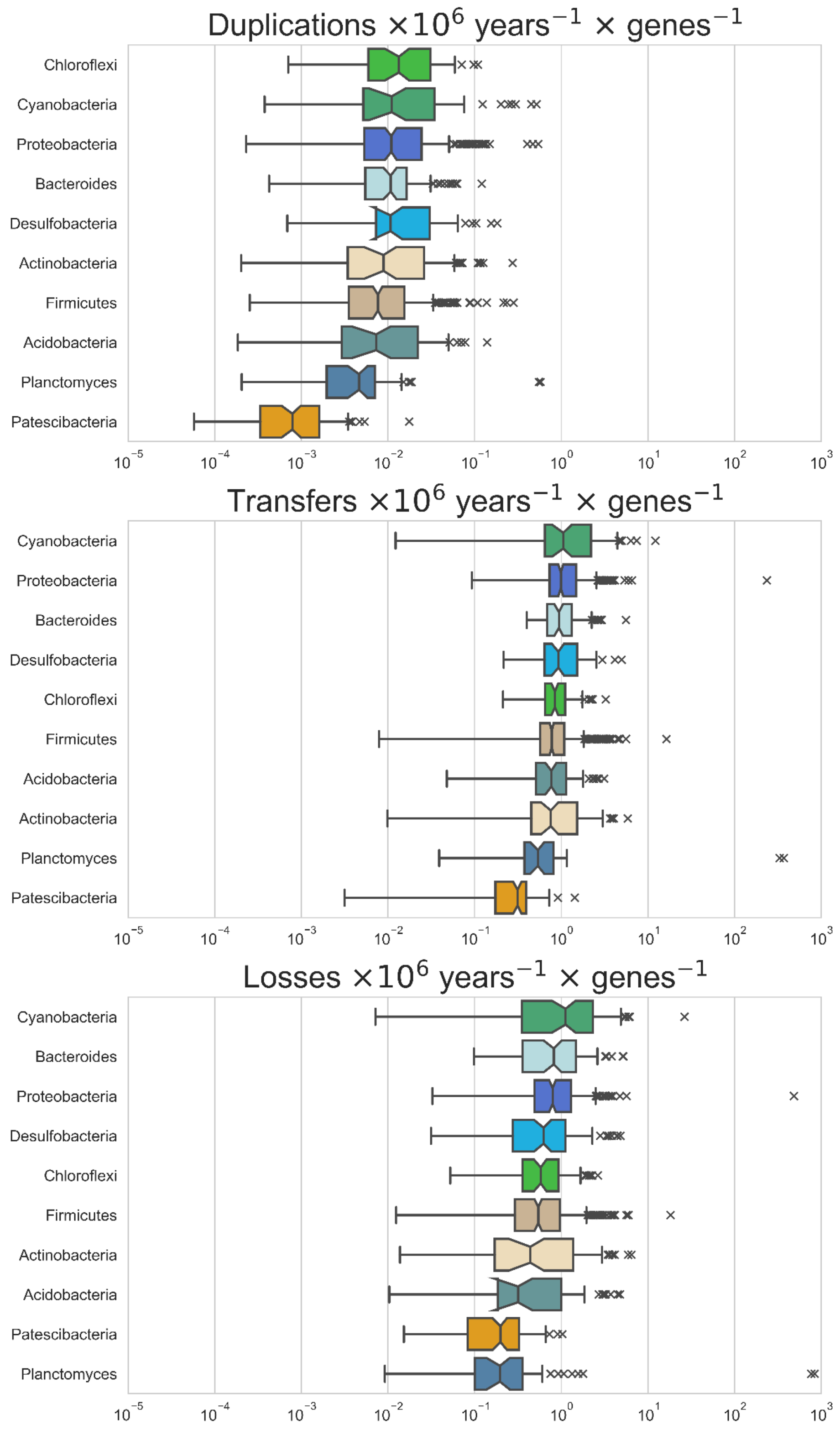


***Figure S17: Inferred rates of gene duplication, transfer and loss (inferred events per gene family per million years) in major bacterial phyla.*** *Patterns within individual phyla reflect the overall finding of a low rate of gene duplication relative to gene transfers and losses in Bacteria.*

### 7. Enrichment analysis

Every gene family was annotated with GO terms using Interproscan 5.62 (Jones et al. 2014). We used the phylogenetic reconciliations obtained with the time-constrained DTL model in ALE to measure the number of acquisitions in every branch, with acquisitions being defined as the sum of transfers and originations. We considered that a gene family is seen as acquired in a given branch if the acquisitions $\leq 0.5$.

For each branch, we recorded the frequency of a GO term being acquired. Following this, a one-sided Fisher test of proportions was used to test for significance. Our testing covered two conditions: the enrichment of GO terms for those branches transitioning to aerobic metabolism compared to anaerobic branches, and the enrichment of GO terms for branches reverting to anaerobic metabolism compared to all the aerobic branches.

After applying a Bonferroni correction for multiple testing, 23 GO terms were found significant for the first condition and only 2 for the second condition. The results can be found in the Online Data Supplement in the file `GO_enrichments.xlsx`.

### 8. Origin of core photosynthesis genes

Our gene tree-species tree reconciliation analyses indicate that the oldest node on the species tree to which oxygenic photosynthesis can be mapped is the common ancestor photosynthetic Cyanobacteria, as expected based on the distribution of these genes in extant taxa (Figure S18-S20).

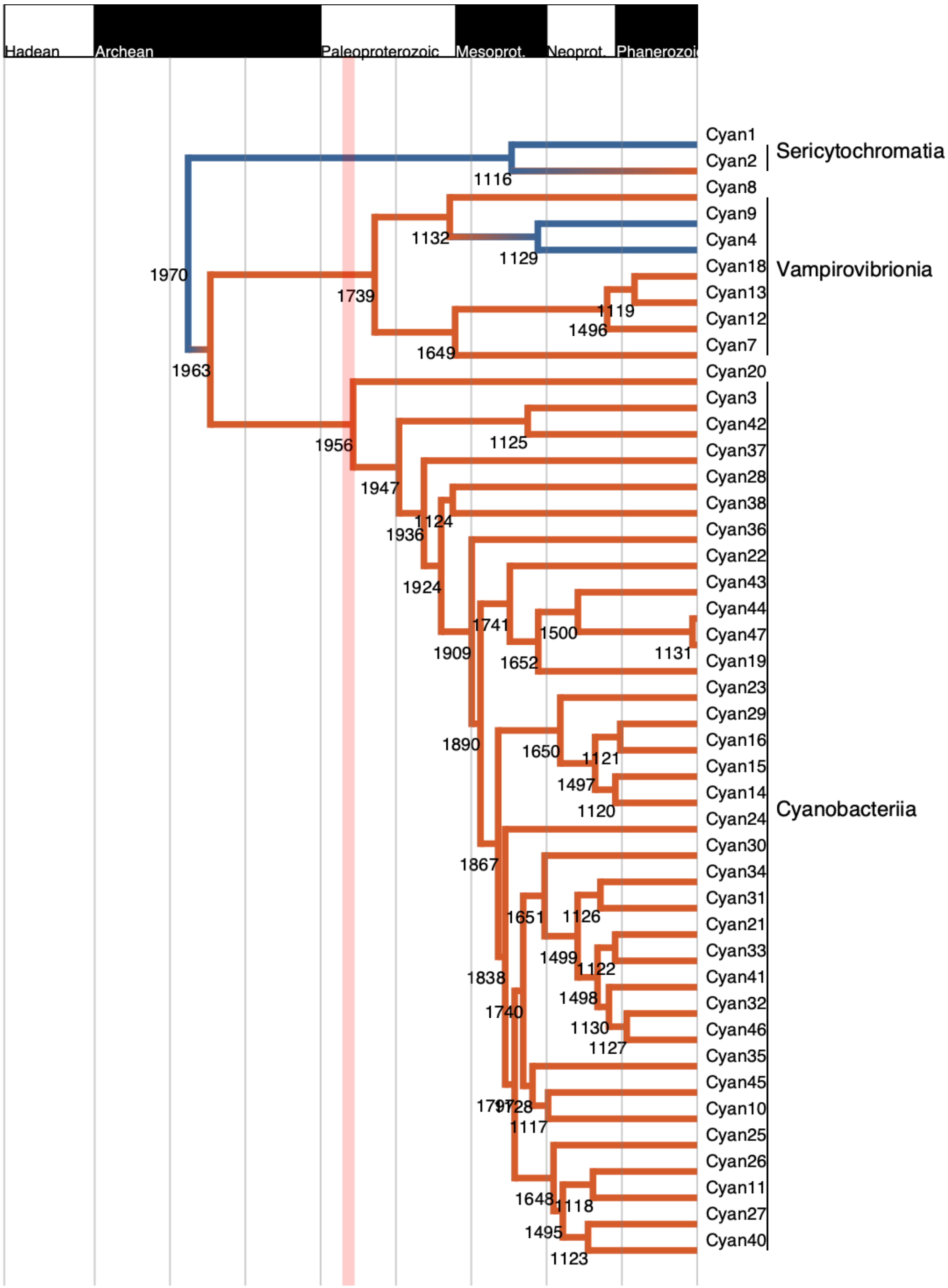


***Figure S18. Zoom on the Cyanobacteria genomes.*** *In this figure we show only the 3 classes within the Cyanobacteria phyla. Node 1963 is inferred to be the oldest aerobic node in the tree.*

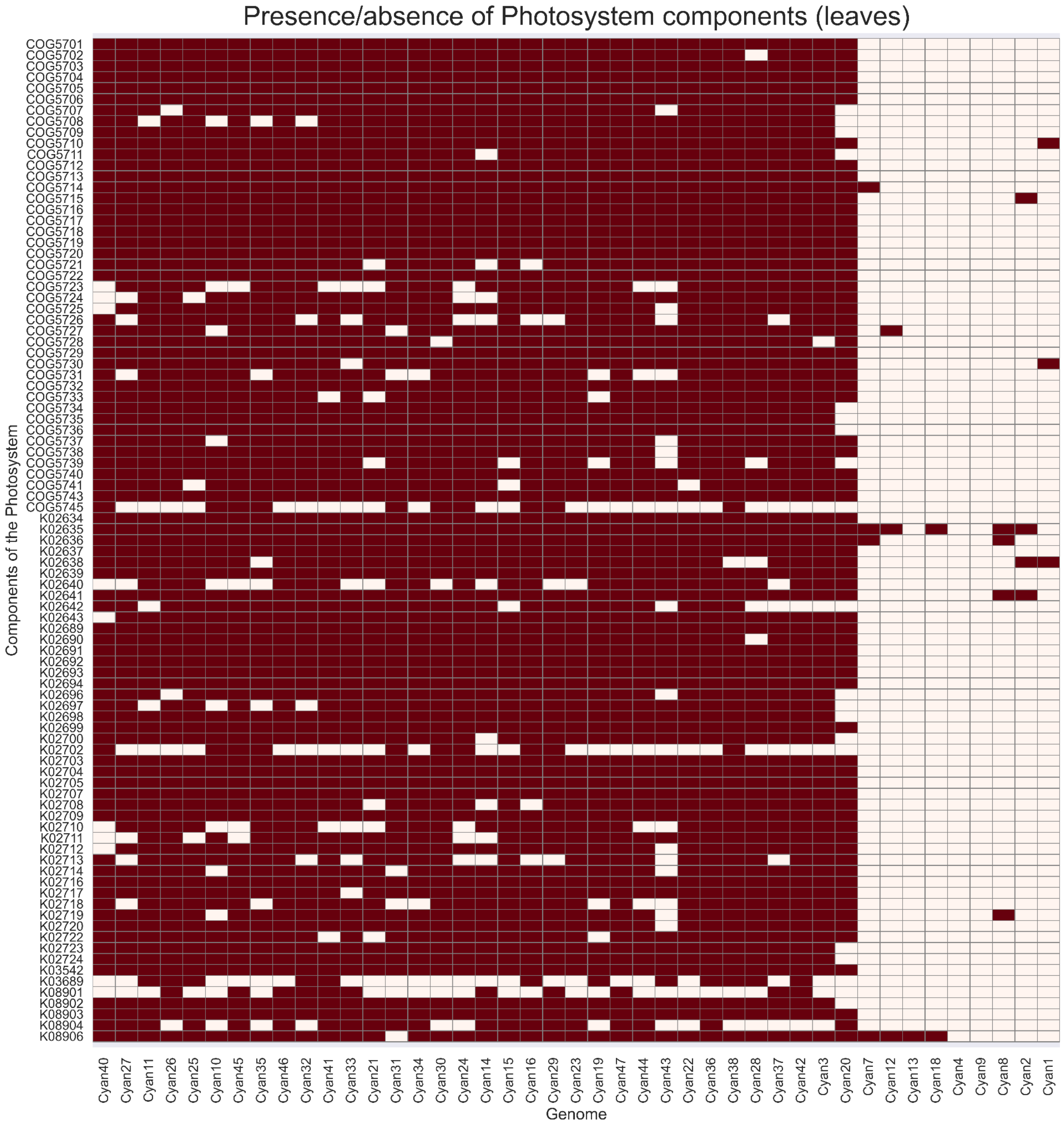


***Figure S19. Presence/absence heatmap for gene families involved in photosynthesis.*** *In this matrix we represent only the presence/absence values for extant genomes.*

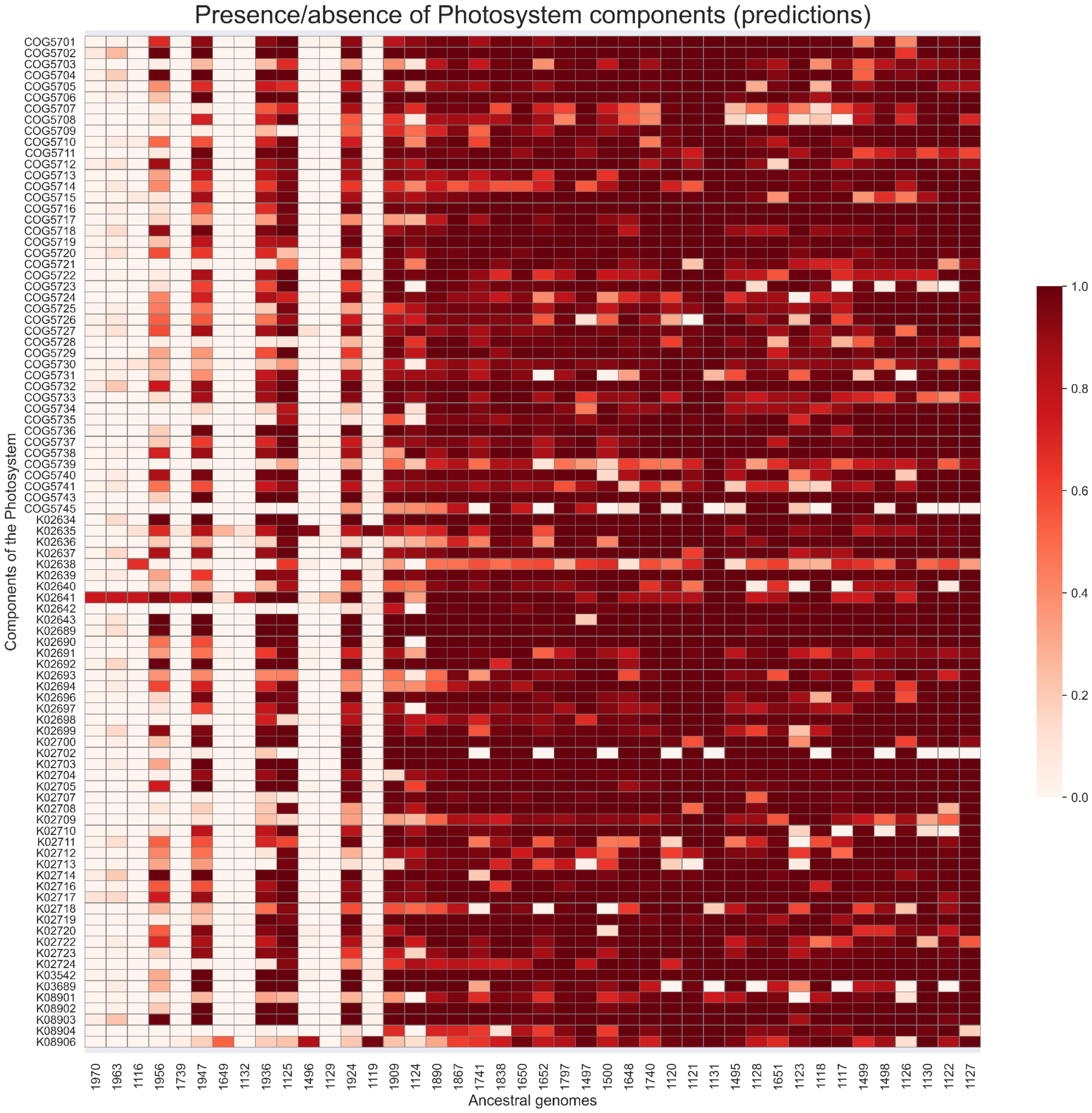


***Figure S20. Presence/absence heatmap for gene families involved in photosynthesis.*** *In this matrix we represent only the presence/absence values for the ancestral genomes, reconstructed with the time-constrained DTL model implemented in ALE.*

### 9. Analysis of bacterial disparity through time

We made use of our timescale to investigate the evolution of bacterial disparity through time. Disparity, or phenotypic diversity, is a concept from palaeontology that refers to the range of phenotypes exhibited by a clade (Guillerme et al. 2020). For single-celled organisms, variation in gene content is a good proxy for phenotypic variation. To evaluate how bacterial gene repertoires have evolved over time, we estimated the genome content distance based on the euclidean distance derived from a matrix of gene family presence or absence among the tips and the internal nodes of the phylogeny and applied applied non-metric multidimensional scaling to quantify the disparity among the gene content of modern and ancestral Bacteria, using the reconstructed ancestral gene repertoires as ancestral states. Plotting disparity through time (Figure S21) provided clear support for an “early burst” model, in which most variation in bacterial gene repertoires was already established prior to 3 Ga (Figure S21(a), which is also Figure 3(a) of the main text). The initial increase in disparity >3.5Ga is associated with the radiation of the Terrabacteria and Gracilicutes clades. This result is consistent with the previous proposal of an Archaean “genetic expansion” (David and Alm 2010), but similar patterns have been reported for other quite different clades at different times in Earth's history including plants and animals. Thus, a phenomenon in which most of the evolutionary innovation in a clade occurs early in its history may be quite general.

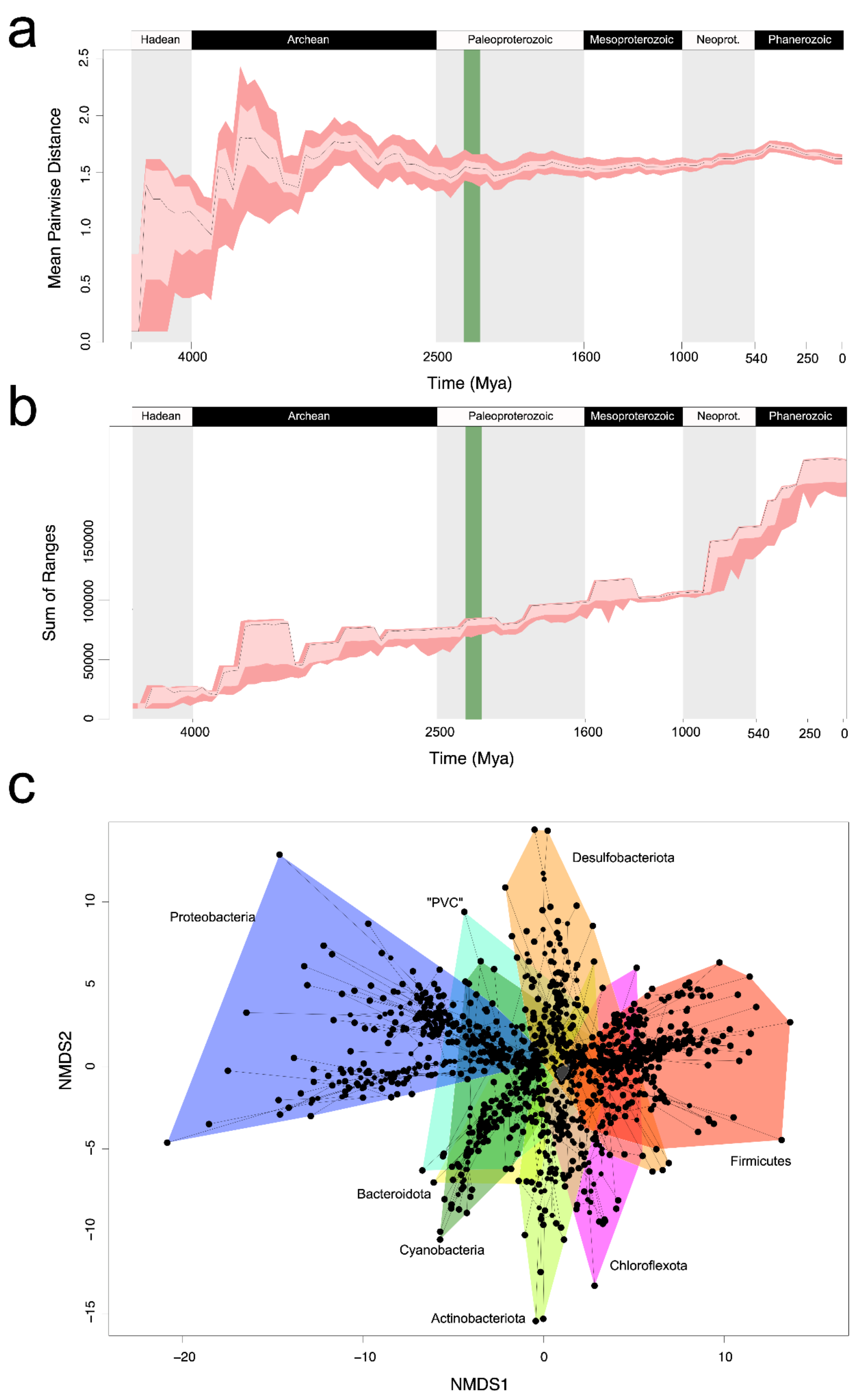


***Figure S21: Evolution of bacterial gene content disparity through time.*** *(a) Mean pairwise distance between the gene content profiles of contemporary lineages through time. The analysis demonstrates that gene content disparity evolved early in bacterial evolution, with a level of variation comparable to sampled extant bacteria already achieved prior to 3Ga. (b) The sum of ranges between the gene content profiles of contemporary lineages through time. The analysis shows a continued expansion of gene content diversity, with an uptick beginning at 0.8 Ga, coinciding with the origin of orders and families.* (c) *the genome space formed by the distance among the gene contents of the major bacterial lineages (n > 10), demonstrating a continuously occupied space with some distinction among lineages.*

### 10. Analysis of aerobic vs anaerobic diversification

Reconstructing diversification rates from dated phylogenies is difficult due to the confounding effect of sampling, as a result of which a given chronogram is compatible with multiple diversification histories (Louca and Pennell 2020). As result we did not attempt to infer diversification patterns directly, but only compared relative diversification rates between aerobic (henceforth branches labelled $1$) and anaerobic lineages (henceforth branches labelled $0$).
To do so we used two metrics each calculated using stochastic resampling based on the posterior density of nodes according to sampled chronograms in 100 My wide sliding windows every 10 Mys. The first metric, that we term "*diversification advantage*", was based on branch lengths in each window and calculated as the ratio of the inverse of branch lengths for branches of type $1 \rightarrow 1$ and type $0 \rightarrow 0$, as a proxy for apparent diversification rate. The second metric, that we call "*innovation advantage*", was based on node density in each window and calculated as the ratio of i) the number of branches of type $0 \rightarrow 1$ divided by the number of branches of type $0 \rightarrow 0$, as a proxy of the rate of transition per lineage to aerobicity, and ii) the number of branches of type $1 \rightarrow 0$ divided by the number of branches of type $1 \rightarrow 1$, as a proxy of the rate of reversion per lineage to anaerobicity.
We also evaluated the possibility that the diversification advantage for aerobic lineages identified using both proxies might arise as a result of our over-sampling of aerobes in the 1007 species dataset. However, comparing the proportion of aerobes in our dataset to the 57,085 species of GTDB 08-RS214 revealed that our dataset was slightly under-sampled for aerobes (44% vs 53% in the GTDB). As an additional test of the impact of sampling on diversification metrics, we randomly dropped 25% (~100 genomes) of aerobic genomes in several replicates (**Figure S22**). We found that the general pattern of two waves of diversification coinciding with sustained increases in atmospheric oxygen levels was robust to this subsampling, with innovation advantage dropping below unity due to the systematic thinning of aerobic lineages reducing the density of nodes of type $1 \rightarrow 1$, to which the metric is proportional.

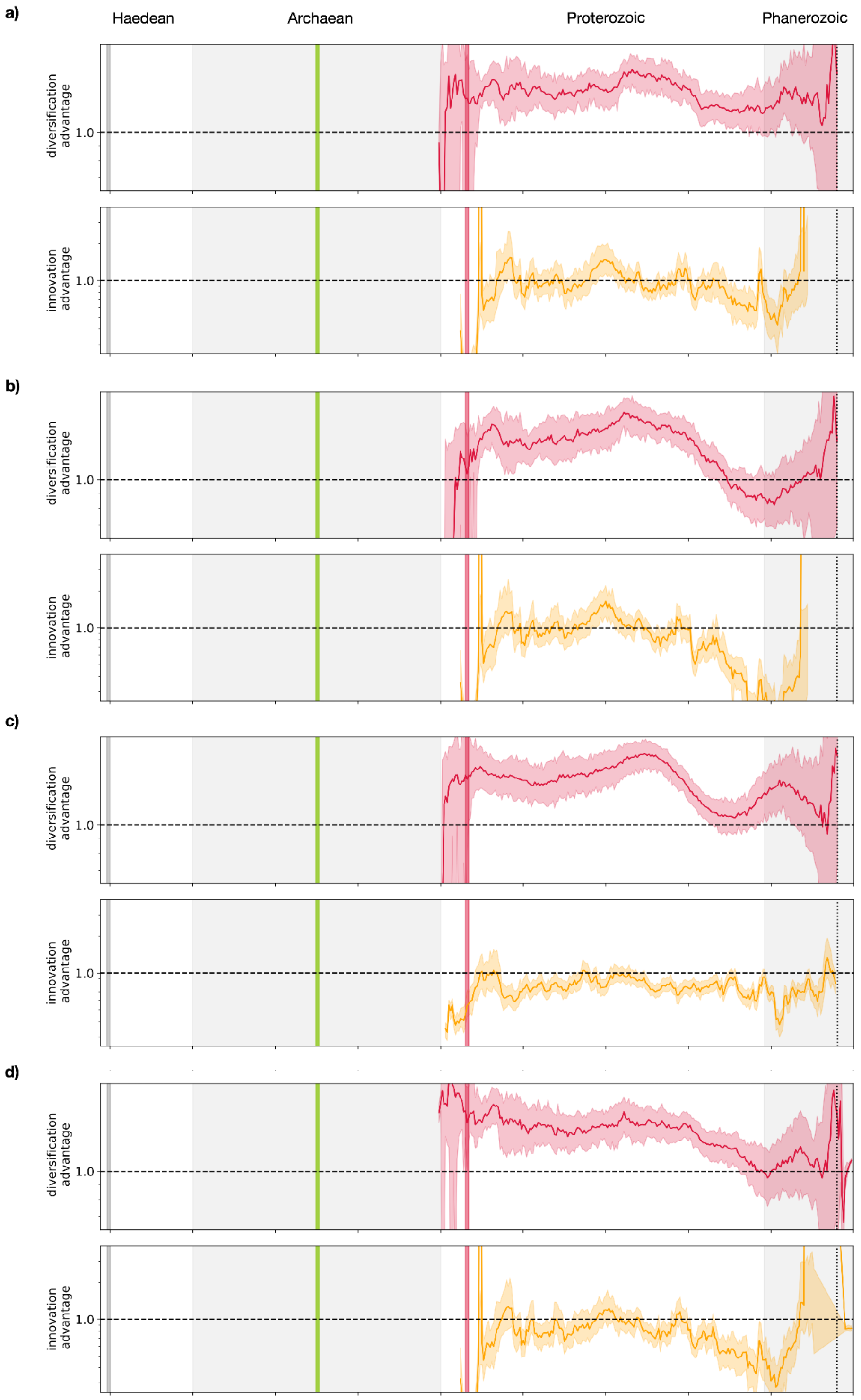


***Figure S22: Random subsampling of aerobic lineages*** *(a-d) we randomly dropped 25% (~100 genomes) of aerobic genomes and recalculated the diversification advantage and innovation advantage metrics of Figure 3. The general pattern of two waves of diversification coinciding with sustained increases in atmospheric oxygen levels was robust to this subsampling, with innovation advantage dropping below unity due to the systematic thinning of aerobic lineages reducing the density of nodes of type* $1 \rightarrow 1$*, to which the metric is proportional. The grey line shows the Moon-forming impact at ~4.52 Ga; the green line at ~3.23 Ga reflects the presence of fossil and isotopic evidence for oxygenic photosynthesis; the red line at ~2.33 Ga reflects the end of the GOE (see calibrations above).*

Coder, David Martin. 1981. *The Chlorellavorus Bacterium: An Antagonistic Symbiont of the*

*Green Alga Chlorella*.
Coleman, Gareth A., Adrián A. Davín, Tara A. Mahendrarajah, Lénárd L. Szánthó, Anja Spang, Philip Hugenholtz, Gergely J. Szöllősi, and Tom A. Williams. 2021. “A Rooted Phylogeny Resolves Early Bacterial Evolution.” *Science* 372 (6542). https://doi.org/10.1126/science.abe0511.
Criscuolo, Alexis, and Simonetta Gribaldo. 2010. “BMGE (Block Mapping and Gathering with Entropy): A New Software for Selection of Phylogenetic Informative Regions from Multiple Sequence Alignments.” *BMC Evolutionary Biology* 10 (January): 210.
David, Lawrence a., and Eric J. Alm. 2010. “Rapid Evolutionary Innovation during an Archaean Genetic Expansion.” *Nature* 469 (7328): 93–96.
Davín, Adrián A., Théo Tricou, Eric Tannier, Damien M. de Vienne, and Gergely J. Szöllősi. 2020. “Zombi: A Phylogenetic Simulator of Trees, Genomes and Sequences That Accounts for Dead Linages.” *Bioinformatics* 36 (4): 1286–88.
Demoulin, Catherine F., Yannick J. Lara, Luc Cornet, Camille François, Denis Baurain, Annick Wilmotte, and Emmanuelle J. Javaux. 2019. “Cyanobacteria Evolution: Insight from the Fossil Record.” *Free Radical Biology & Medicine* 140 (August): 206–23.
Doolittle, W. Ford, and Austin Booth. 2017. “It’s the Song, Not the Singer: An Exploration of Holobiosis and Evolutionary Theory.” *Biology & Philosophy* 32 (1): 5–24.
Dunn, Anne K. 2023. “Alternative Oxidase in Bacteria.” *Biochimica et Biophysica Acta (BBA) - Bioenergetics* 1864 (1): 148929.
Eddy, Sean R. 2011. “Accelerated Profile HMM Searches.” *PLoS Computational Biology* 7 (10): e1002195.
Finn, Robert D., Jody Clements, and Sean R. Eddy. 2011. “HMMER Web Server: Interactive Sequence Similarity Searching.” *Nucleic Acids Research* 39 (Web Server issue): W29–37.
Fournier, G. P., K. R. Moore, L. T. Rangel, J. G. Payette, L. Momper, and T. Bosak. 2021. “The Archean Origin of Oxygenic Photosynthesis and Extant Cyanobacterial Lineages.” *Proceedings. Biological Sciences / The Royal Society* 288 (1959): 20210675.
Fralick, Philip, Don W. Davis, and Stephen A. Kissin. 2002. “The Age of the Gunflint Formation, Ontario, Canada: Single Zircon U‐Pb Age Determinations from Reworked Volcanic Ash.” *Canadian Journal of Earth Sciences* 39 (7): 1085–91.
Gardiner, Nicholas J., David Wacey, Christopher L. Kirkland, Tim E. Johnson, and Heejin Jeon. 2019. “Zircon U–Pb, Lu–Hf and O Isotopes from the 3414 Ma Strelley Pool Formation, East Pilbara Terrane, and the Palaeoarchaean Emergence of a Cryptic Cratonic Core.” *Precambrian Research* 321 (February): 64–84.
Gibson, Timothy M., Patrick M. Shih, Vivien M. Cumming, Woodward W. Fischer, Peter W. Crockford, Malcolm S. W. Hodgskiss, Sarah Wörndle, et al. 2018. “Precise Age of Bangiomorpha Pubescens Dates the Origin of Eukaryotic Photosynthesis.” *Geology*. https://doi.org/10.1130/g39829.1.
Guillerme, Thomas, Natalie Cooper, Stephen L. Brusatte, Katie E. Davis, Andrew L. Jackson, Sylvain Gerber, Anjali Goswami, et al. 2020. “Disparities in the Analysis of Morphological Disparity.” *Biology Letters* 16 (7): 20200199.
Hanan, B. B., and G. R. Tilton. 1987. “60025: Relict of Primitive Lunar Crust?” *Earth and Planetary Science Letters*. https://doi.org/10.1016/0012-821x(87)90171-3.
Harris, Brogan J., James W. Clark, Dominik Schrempf, Gergely J. Szöllősi, Philip C. J. Donoghue, Alistair M. Hetherington, and Tom A. Williams. 2022. “Divergent Evolutionary Trajectories of Bryophytes and Tracheophytes from a Complex Common Ancestor of Land Plants.” *Nature Ecology & Evolution* 6 (September): 1634–43.
Hodgskiss, Malcolm S. W., Peter W. Crockford, Yongbo Peng, Boswell A. Wing, and Tristan J. Horner. 2019. “A Productivity Collapse to End Earth’s Great Oxidation.” *Proceedings of the National Academy of Sciences of the United States of America* 116 (35): 17207–12.
Huerta-Cepas, Jaime, Kristoffer Forslund, Luis Pedro Coelho, Damian Szklarczyk, Lars Juhl Jensen, Christian von Mering, and Peer Bork. 2017. “Fast Genome-Wide Functional Annotation through Orthology Assignment by eggNOG-Mapper.” *Molecular Biology and*

*Evolution* 34 (8): 2115–22.
Jabłońska, Jagoda, and Dan S. Tawfik. 2019. “The Number and Type of Oxygen-Utilizing Enzymes Indicates Aerobic vs. Anaerobic Phenotype.” *Free Radical Biology & Medicine* 140 (August): 84–92.
Jones, Philip, David Binns, Hsin-Yu Chang, Matthew Fraser, Weizhong Li, Craig McAnulla, Hamish McWilliam, et al. 2014. “InterProScan 5: Genome-Scale Protein Function Classification.” *Bioinformatics* 30 (9): 1236–40.
Kamo, S. L., and D. W. Davis. 1994. “Reassessment of Archean Crustal Development in the Barberton Mountain Land, South Africa, Based on U-Pb Dating.” *Tectonics* 13 (1): 167–92.
Katoh, Kazutaka, and Daron M. Standley. 2013. “MAFFT Multiple Sequence Alignment Software Version 7: Improvements in Performance and Usability.” *Molecular Biology and Evolution* 30 (4): 772–80.
Kim, Jaehee, Noah A. Rosenberg, and Julia A. Palacios. 2020. “Distance Metrics for Ranked Evolutionary Trees.” *Proceedings of the National Academy of Sciences of the United States of America* 117 (46): 28876–86.
Kramer, Oliver. 2016. “Scikit-Learn.” In *Machine Learning for Evolution Strategies*, edited by Oliver Kramer, 45–53. Cham: Springer International Publishing.
Libertín, Milan, Jiří Kvaček, Jiří Bek, Viktor Žárský, and Petr Štorch. 2018. “Sporophytes of Polysporangiate Land Plants from the Early Silurian Period May Have Been Photosynthetically Autonomous.” *Nature Plants* 4 (5): 269–71.
Li, Huaikun, Songnian Lu, Wenbo Su, Zhenqun Xiang, Hongying Zhou, and Yongqing Zhang. 2013. “Recent Advances in the Study of the Mesoproterozoic Geochronology in the North China Craton.” *Journal of Asian Earth Sciences*. https://doi.org/10.1016/j.jseaes.2013.02.020.
Loron, Corentin C., Camille François, Robert H. Rainbird, Elizabeth C. Turner, Stephan Borensztajn, and Emmanuelle J. Javaux. 2019. “Early Fungi from the Proterozoic Era in Arctic Canada.” *Nature* 570 (7760): 232–35.
Loron, Corentin C., Robert H. Rainbird, Elizabeth C. Turner, J. Wilder Greenman, and Emmanuelle J. Javaux. 2019. “Organic-Walled Microfossils from the Late Mesoproterozoic to Early Neoproterozoic Lower Shaler Supergroup (Arctic Canada): Diversity and Biostratigraphic Significance.” *Precambrian Research*. https://doi.org/10.1016/j.precamres.2018.12.024.
Louca, Stilianos, and Matthew W. Pennell. 2020. “Extant Timetrees Are Consistent with a Myriad of Diversification Histories.” *Nature* 580 (7804): 502–5.
Lundberg, Scott M., Gabriel Erion, Hugh Chen, Alex DeGrave, Jordan M. Prutkin, Bala Nair, Ronit Katz, Jonathan Himmelfarb, Nisha Bansal, and Su-In Lee. 2020. “From Local Explanations to Global Understanding with Explainable AI for Trees.” *Nature Machine Intelligence* 2 (1): 56–67.
Mahendrarajah, Tara A., Edmund R. R. Moody, Dominik Schrempf, Lénárd L. Szánthó, Nina Dombrowski, Adrián A. Davín, Davide Pisani, et al. 2023. “ATP Synthase Evolution on a Cross-Braced Dated Tree of Life.” *bioRxiv*. https://doi.org/10.1101/2023.04.11.536006.
Maloof, Adam C., Jahandar Ramezani, Samuel A. Bowring, David A. Fike, Susannah M. Porter, and Mohamed Mazouad. 2010. “Constraints on Early Cambrian Carbon Cycling from the Duration of the Nemakit-Daldynian–Tommotian Boundary δ13C Shift, Morocco.” *Geology*. https://doi.org/10.1130/g30726.1.
Martin, Adam P., Daniel J. Condon, Anthony R. Prave, and Aivo Lepland. 2013. “A Review of Temporal Constraints for the Palaeoproterozoic Large, Positive Carbonate Carbon Isotope Excursion (the Lomagundi–Jatuli Event).” *Earth-Science Reviews* 127 (December): 242–61.
Melchin, M. J., P. M. Sadler, and B. D. Cramer. 2020. “The Silurian Period.” *Geologic Time Scale 2020*. https://www.sciencedirect.com/science/article/pii/B9780128243602000218.
Minh, Bui Quang, Heiko A. Schmidt, Olga Chernomor, Dominik Schrempf, Michael D. Woodhams, Arndt von Haeseler, and Robert Lanfear. 2020. “IQ-TREE 2: New Models and Efficient Methods for Phylogenetic Inference in the Genomic Era.” *Molecular*

*Biology and Evolution* 37 (5): 1530–34.
Moody, Edmund R. R., Tara A. Mahendrarajah, Nina Dombrowski, James W. Clark, Celine Petitjean, Pierre Offre, Gergely J. Szöllősi, Anja Spang, and Tom A. Williams. 2022. “An Estimate of the Deepest Branches of the Tree of Life from Ancient Vertically Evolving Genes.” *eLife* 11 (February). https://doi.org/10.7554/eLife.66695.
Morel, Benoit, Alexey M. Kozlov, Alexandros Stamatakis, and Gergely J. Szöllősi. 2020. “GeneRax: A Tool for Species-Tree-Aware Maximum Likelihood-Based Gene Family Tree Inference under Gene Duplication, Transfer, and Loss.” *Molecular Biology and Evolution* 37 (9): 2763–74.
Muñoz-Gómez, Sergio A., Sebastian Hess, Gertraud Burger, B. Franz Lang, Edward Susko, Claudio H. Slamovits, and Andrew J. Roger. 2019. “An Updated Phylogeny of the Alphaproteobacteria Reveals That the Parasitic Rickettsiales and Holosporales Have Independent Origins.” *eLife*. https://doi.org/10.7554/elife.42535.
Munson, T. J., S. W. Denyszyn, J. M. Simmons, and M. Kunzmann. 2020. “A 1642 Ma Age for the Fraynes Formation, Birrindudu Basin, Confirms Correlation with the Economically Significant Barney Creek Formation, McArthur Basin, Northern Territory.” *Australian Journal of Earth Sciences*. https://doi.org/10.1080/08120099.2020.1669708.
Murali, Ranjani, Robert B. Gennis, and James Hemp. 2021. “Evolution of the Cytochrome Bd Oxygen Reductase Superfamily and the Function of CydAA’ in Archaea.” *The ISME Journal* 15 (12): 3534–48.
Murali, Ranjani, James Hemp, and Robert B. Gennis. 2022. “Evolution of Quinol Oxidation within the Heme-copper Oxidoreductase Superfamily.” *Biochimica et Biophysica Acta, Bioenergetics* 1863 (8): 148907.
Narbonne, G. M., P. M. Myrow, and E. Landing. 1987. “A Candidate Stratotype for the Precambrian–Cambrian Boundary, Fortune Head, Burin Peninsula, Southeastern Newfoundland.” *Canadian Journal of*. https://cdnsciencepub.com/doi/abs/10.1139/e87-124?casa_token=GC6isEa6X_cAAAAA:gRsty72DDcW14iKKgODsivvF6Hi0Hs9H0txgmlfE5_JxeR5bzIXdZnrkomhkw3Npzu6k6txzPCxfebg.
Nelson, D. R. n.d. “178043: Lithic-Quartz Sandstone, Strelley Pool; Geochronology Dataset 565.” In *Compilation of Geochronology Data, June 2007 Update*. Geological Survey of Western Australia.
Oliver, Thomas, Patricia Sánchez-Baracaldo, Anthony W. Larkum, A. William Rutherford, and Tanai Cardona. 2021. “Time-Resolved Comparative Molecular Evolution of Oxygenic Photosynthesis.” *Biochimica et Biophysica Acta, Bioenergetics* 1862 (6): 148400.
Page, R. W., and I. P. Sweet. 1998. “Geochronology of Basin Phases in the Western Mt Isa Inlier, and Correlation with the McArthur Basin∗.” *Australian Journal of Earth Sciences*. https://doi.org/10.1080/08120099808728383.
Parham, James F., Philip C. J. Donoghue, Christopher J. Bell, Tyler D. Calway, Jason J. Head, Patricia A. Holroyd, Jun G. Inoue, et al. 2012. “Best Practices for Justifying Fossil Calibrations.” *Systematic Biology* 61 (2): 346–59.
Parks, Donovan H., Maria Chuvochina, Pierre-Alain Chaumeil, Christian Rinke, Aaron J. Mussig, and Philip Hugenholtz. 2020. “A Complete Domain-to-Species Taxonomy for Bacteria and Archaea.” *Nature Biotechnology* 38 (9): 1079–86.
Parks, Donovan H., Maria Chuvochina, David W. Waite, Christian Rinke, Adam Skarshewski, Pierre-Alain Chaumeil, and Philip Hugenholtz. 2018. “A Standardized Bacterial Taxonomy Based on Genome Phylogeny Substantially Revises the Tree of Life.” *Nature Biotechnology* 36 (10): 996–1004.
Peng, S. C., L. E. Babcock, and P. Ahlberg. 2020. “The Cambrian Period.” In *Geologic Time Scale 2020*, edited by Felix M. Gradstein, James G. Ogg, Mark D. Schmitz, and Gabi M. Ogg, 565–629. Elsevier.
Poulton, Simon W., Andrey Bekker, Vivien M. Cumming, Aubrey L. Zerkle, Donald E. Canfield, and David T. Johnston. 2021. “A 200-Million-Year Delay in Permanent Atmospheric Oxygenation.” *Nature* 592 (7853): 232–36.

Reimer, Lorenz Christian, Joaquim Sardà Carbasse, Julia Koblitz, Christian Ebeling, Adam Podstawka, and Jörg Overmann. 2022. “BacDive in 2022: The Knowledge Base for Standardized Bacterial and Archaeal Data.” *Nucleic Acids Research* 50 (D1): D741–46.
Reimer, Lorenz Christian, Anna Vetcininova, Joaquim Sardà Carbasse, Carola Söhngen, Dorothea Gleim, Christian Ebeling, and Jörg Overmann. 2019. “BacDive in 2019: Bacterial Phenotypic Data for High-Throughput Biodiversity Analysis.” *Nucleic Acids Research* 47 (D1): D631–36.
Runnegar, Bruce. 1981. “Muscle Scars, Shell Form and Torsion in Cambrian and Ordovician Univalved Molluscs.” *Lethaia*. https://doi.org/10.1111/j.1502-3931.1981.tb01104.x.
Satkoski, Aaron M., Nicolas J. Beukes, Weiqiang Li, Brian L. Beard, and Clark M. Johnson. 2015. “A Redox-Stratified Ocean 3.2 Billion Years Ago.” *Earth and Planetary Science Letters* 430 (November): 43–53.
Schirrmeister, Bettina E., Patricia Sanchez-Baracaldo, and David Wacey. 2016. “Cyanobacterial Evolution during the Precambrian.” *International Journal of Astrobiology* 15 (3): 187–204.
Schrempf, Dominik, Nicolas Lartillot, and Gergely Szöllösi. 2020. “Scalable Empirical Mixture Models That Account for across-Site Compositional Heterogeneity.” *Molecular Biology and Evolution*, September. https://doi.org/10.1093/molbev/msaa145.
Sergeev, V. N., Mukund Sharma, and Yogmaya Shukla. 2012. “Proterozoic Fossil Cyanobacteria.” *The Journal of Physiological Sciences: JPS* 61 ((1-2)): 189–358.
Shih, P. M., J. Hemp, L. M. Ward, N. J. Matzke, and W. W. Fischer. 2017. “Crown Group Oxyphotobacteria Postdate the Rise of Oxygen.” *Geobiology* 15 (1): 19–29.
Soo, Rochelle M., James Hemp, Donovan H. Parks, Woodward W. Fischer, and Philip Hugenholtz. 2017. “On the Origins of Oxygenic Photosynthesis and Aerobic Respiration in Cyanobacteria.” *Science* 355 (6332): 1436–40.
Stairs, Courtney W., Jennah E. Dharamshi, Daniel Tamarit, Laura Eme, Steffen L. Jørgensen, Anja Spang, and Thijs J. G. Ettema. 2020. “Chlamydial Contribution to Anaerobic Metabolism during Eukaryotic Evolution.” *Science Advances* 6 (35): eabb7258.
Steiner, Michael, Guoxiang Li, Yi Qian, Maoyan Zhu, and Bernd-Dietrich Erdtmann. 2007. “Neoproterozoic to Early Cambrian Small Shelly Fossil Assemblages and a Revised Biostratigraphic Correlation of the Yangtze Platform (China).” *Palaeogeography, Palaeoclimatology, Palaeoecology*. https://doi.org/10.1016/j.palaeo.2007.03.046.
Strother, Paul K. 2016. “Systematics and Evolutionary Significance of Some New Cryptospores from the Cambrian of Eastern Tennessee, USA.” *Review of Palaeobotany and Palynology* 227 (April): 28–41.
Strother, P. K., and J. H. Beck. 2000. “Spore-like Microfossils from Middle Cambrian Strata: Expanding the Meaning of the Term Cryptospore.” In *Pollen and Spores: Morphology and Biology*, edited by M. M. Harley and Morton C M Blackmore. Royal Botanic Gardens, Kew.
Szöllõsi, Gergely J., Sebastian Höhna, Tom A. Williams, Dominik Schrempf, Vincent Daubin, and Bastien Boussau. 2022. “Relative Time Constraints Improve Molecular Dating.” *Systematic Biology* 71 (4): 797–809.
Szöllõsi, Gergely J., Wojciech Rosikiewicz, Bastien Boussau, Eric Tannier, and Vincent Daubin. 2013. “Efficient Exploration of the Space of Reconciled Gene Trees.” *Systematic Biology* 62 (6): 901–12.
Szöllosi, Gergely J., Eric Tannier, Nicolas Lartillot, and Vincent Daubin. 2013. “Lateral Gene Transfer from the Dead.” *Systematic Biology* 62 (3): 386–97.
Treangen, Todd J., and Eduardo P. C. Rocha. 2011. “Horizontal Transfer, Not Duplication, Drives the Expansion of Protein Families in Prokaryotes.” *PLoS Genetics* 7 (1): e1001284.
Tria, Fernando D. K., and William F. Martin. 2021. “Gene Duplications Are At Least 50 Times Less Frequent than Gene Transfers in Prokaryotic Genomes.” *Genome Biology and Evolution* 13 (10). https://doi.org/10.1093/gbe/evab224.
Tria, Fernando Domingues Kümmel, Giddy Landan, and Tal Dagan. 2017. “Phylogenetic

Rooting Using Minimal Ancestor Deviation." *Nature Ecology & Evolution* 1: s41559–017.
Wang, Huai-Chun, Bui Quang Minh, Edward Susko, and Andrew J. Roger. 2018. "Modeling Site Heterogeneity with Posterior Mean Site Frequency Profiles Accelerates Accurate Phylogenomic Estimation." *Systematic Biology* 67 (2): 216–35.
Weimann, Aaron, Kyra Mooren, Jeremy Frank, Phillip B. Pope, Andreas Bremges, and Alice C. McHardy. 2016. "From Genomes to Phenotypes: Traitar, the Microbial Trait Analyzer." *mSystems* 1 (6). https://doi.org/10.1128/mSystems.00101-16.
Wickham, Hadley. 2016. *ggplot2: Elegant Graphics for Data Analysis*. Springer.
Williams, Tom A., Adrian A. Davin, Benoit Morel, Lénárd L. Szánthó, Anja Spang, Alexandros Stamatakis, Philip Hugenholtz, and Gergely J. Szöllősi. 2023. "Parameter Estimation and Species Tree Rooting Using ALE and GeneRax." *Genome Biology and Evolution*, July. https://doi.org/10.1093/gbe/evad134.
Wolfe, Joanna M., Allison C. Daley, David A. Legg, and Gregory D. Edgecombe. 2016. "Fossil Calibrations for the Arthropod Tree of Life." *Earth-Science Reviews* 160 (September): 43–110.
Yang, Chuan, Alan D. Rooney, Daniel J. Condon, Xian-Hua Li, Dmitriy V. Grazhdankin, Fred T. Bowyer, Chunlin Hu, Francis A. Macdonald, and Maoyan Zhu. 2021. "The Tempo of Ediacaran Evolution." *Science Advances* 7 (45): eabi9643.
Yin, Zongjun, Weichen Sun, Pengju Liu, Junyuan Chen, David J. Bottjer, Jinhua Li, and Maoyan Zhu. 2022. "Diverse and Complex Developmental Mechanisms of Early Ediacaran Embryo-like Fossils from the Weng'an Biota, Southwest China." *Philosophical Transactions of the Royal Society of London. Series B, Biological Sciences* 377 (1847): 20210032.
Yin, Zongjun, Weichen Sun, Joachim Reitner, and Maoyan Zhu. 2022. "New Holozoans with Cellular Resolution from the Early Ediacaran Weng'an Biota, SW China." *Journal of the Geological Society* 179 (4). https://doi.org/10.1144/jgs2021-061.
Zhang, Shujing, Rong Cao, Zhongwu Lan, Zhensheng Li, Zhuoya Zhao, Bin Wan, Chengguo Guan, and Xunlai Yuan. 2022. "SIMS Pb-Pb Dating of Phosphates in the Proterozoic Strata of SE North China Craton: Constraints on Eukaryote Evolution." *Precambrian Research* 371 (April): 106562.
